# Manifold learning for olfactory habituation to strongly fluctuating backgrounds

**DOI:** 10.1101/2025.05.26.656161

**Authors:** François X. P. Bourassa, Paul François, Gautam Reddy, Massimo Vergassola

## Abstract

Animals rely on their sense of smell to survive, but important olfactory cues are mixed with confounding background odors that fluctuate due to atmospheric turbulence. It is unclear how the olfactory system habituates to such stochastic backgrounds to detect behaviorally important odors. Here, we explicitly consider the high-dimensional nature of odor coding, the natural statistics of odor fluctuations and the architecture of the early olfactory pathway. We show that their combination favors a manifold learning mechanism for olfactory habituation over alternatives based on predictive filtering. Manifold learning is implemented in our model by a biologically plausible network of inhibitory interneurons in the early olfactory pathway. We demonstrate that plasticity rules based on IBCM or online PCA are effective at implementing this mechanism in turbulent conditions and outperform previous models relying on mean background subtraction. Interneurons with an IBCM plasticity rule acquire selectivity to independently varying odors. This manifold learning mechanism offers a path towards distinguishing plasticity rules in experiments and could be leveraged by other biological circuits facing fluctuating environments.

## Introduction

Most of us have experienced an odor fading to imperceptibility after prolonged exposure. Habituation is a basic building block of sensory cognition, allowing us to pay attention to weak but important cues relevant for survival [1, 2]. Across sensory modalities, numerous mechanisms for sensory adaptation and habituation filter out irrelevant information; these mechanisms must be considered in light of the statistical features of natural scenes [3–7]. Though olfactory habituation to regular stimuli is behaviorally well-characterized [8], less is known about its computational basis in neural circuits facing naturalistic environments.

The physics of odor transport poses a difficult habituation problem, challenging simple models such as mean background subtraction. Unlike in vision and audition, olfactory signals are transported by a physical medium that is turbulent at the spatial scales relevant for behavior. Wind velocities in such environments have complex spatial and temporal fluctuations, which segregate air into patches of odor and clean air (Fig. 1A). An olfactory sensory apparatus thus receives a highly intermittent sensory signal, where clumps of intense odor detections (‘whiffs’) are separated by seconds to minutes of relatively clean air (‘blanks’) (Fig. 1B) [9–11]. These strongly non-Gaussian statistics make it non-trivial for olfactory circuits to identify new odors mixed with a dominant, fluctuating background.

**Fig. 1.**
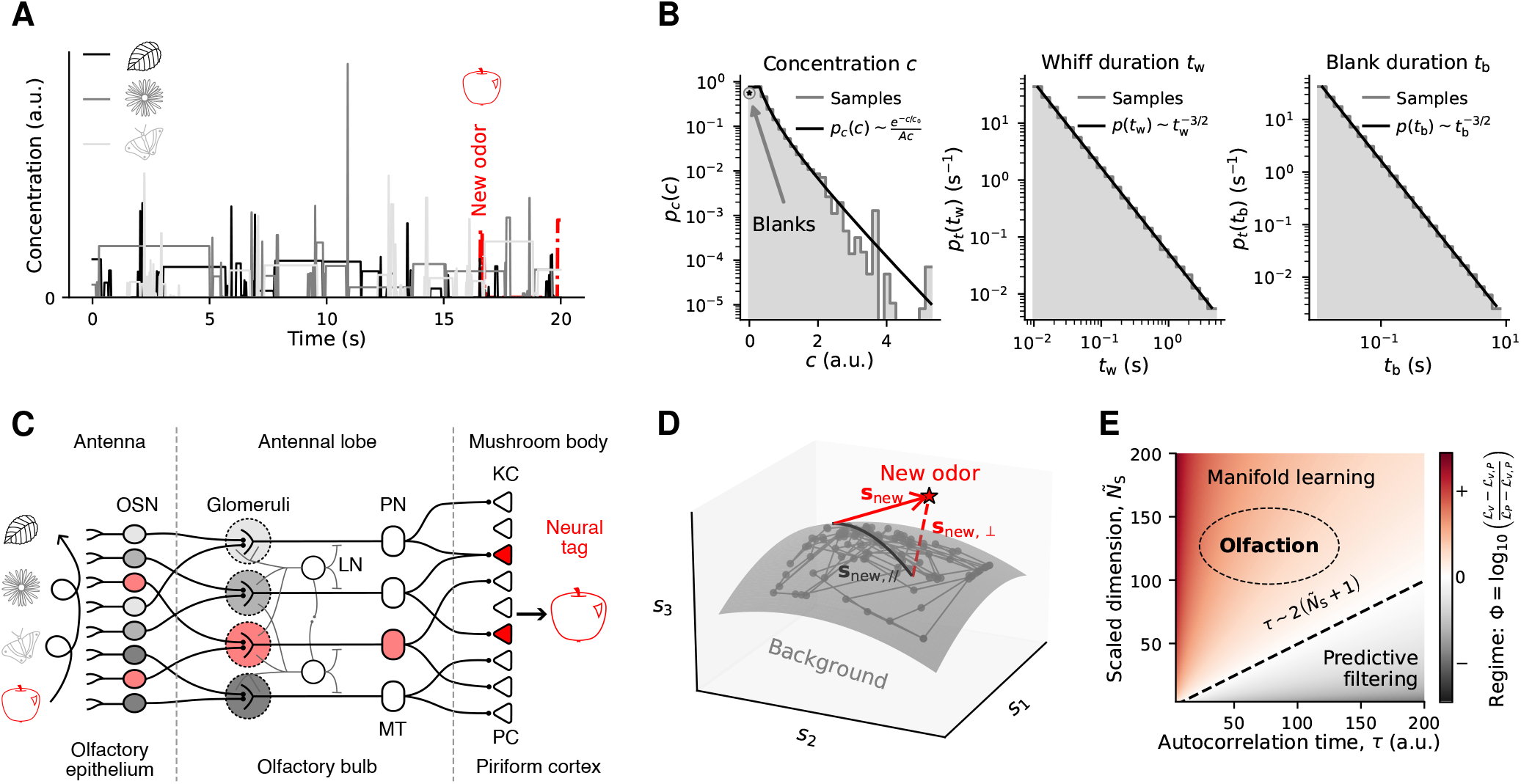
Olfactory systems face stochastic, turbulent odor mixture inputs, for which manifold learning may be an optimal habituation strategy. (**A**) Illustration of background and new odor concentrations time series, strongly fluctuating in a series of whiffs and blanks, according to the turbulent atmosphere statistics derived in [9]. (**B**) Stationary probability distributions of whiff concentrations and whiff and blank durations. (**C**) Structure of early layers in the olfactory network, annotated with fly (top) and mouse (bottom, when different from fly) anatomical regions and cell types. (**D**) Illustration of a hypothetical low-dimensional subspace spanned by background odors in the space of OSN activities *s*_*i*_, with sample mixtures generated by log-normal odor concentrations. A new odor, **s**_new_, generally has a component, **s**_new,⊥_, lying outside of the background manifold. (**E**) Log-ratio of the difference in loss functions for new odor recognition between manifold learning (*ℒ*_*P*_), predictive filtering (*ℒ*_*v*_), or the combination of both strategies (*ℒ*_*v,P*_). For a sensory system tasked to detect new odors within fast background fluctuations in a high-dimensional space, as it is the case with olfaction, manifold learning is the dominant strategy (loss *ℒ*_*P*_ *ℒ*_*v,P*_). The olfactory space dimensionality is rescaled by the relative variance of background and new odors: 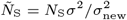.

The neurobiology of early olfaction outlines the underlying circuit structure solving this habituation problem across animal species (Fig. 1C). In insects, odors are first detected by olfactory receptors (ORs) located on olfactory sensory neurons (OSNs) in the antennae. Each OSN often expresses one olfactory receptor type (among ∼ 50 different receptor types in the fruit fly). Axons from OSNs expressing the same receptor project to distinct locations called glomeruli in the antennal lobe. Projection neurons (PNs) integrate signals from a few glomeruli and project to higher order processing centers, including the mushroom body (where associations are formed) and the lateral horn (which drives innate behaviors) [12, 13]. A strikingly similar architecture is present in mammals: glomeruli are located in the olfactory bulb where OSN input is processed and broadcast to diverse subcortical and cortical regions [14].

A characteristic feature of ORs is their broad tuning to many odor molecules, such that each OSN is activated by multiple odors [15]. Thus, OSN activity induced by a behaviorally relevant odor appearing in a naturalistic environment is masked by possibly many background odors [16]. Biophysical mechanisms for adaptation in a single OSN allow for adapting the dynamic range of spiking output to the statistics of receptor activity [17–20]. However, these single-neuron mechanisms do not, on their own, disentangle contributions from a new and relevant odor from those of irrelevant backgrounds, suggesting that background subtraction occurs at the neural population coding level, in a later stage of the olfactory pathway [21–24].

The antennal lobe (AL) in insects and the olfactory bulb (OB) in mammals are likely candidate regions for olfactory habituation. Since most olfactory receptors are promiscuous, the population activity of glomeruli in these regions represents an efficient combinatorial code for odors [25, 26]. The AL and OB further contain extensive local networks of inhibitory interneurons that mediate inter-glomerular crosstalk before signals are broadcast to downstream processing centers [27–30]. Consistent with this picture, plasticity mechanisms in *Drosophila* antennal lobe inhibitory interneurons (called lateral neurons (LNs)) have been linked to the formation of olfactory memories during habituation [6, 8, 22, 31–35]. Building on these results, Shen *et al*. proposed a neural model for olfactory habituation where LNs learn and subtract a time-averaged background signal by integrating glomerular activity over timescales of minutes [36]. While a mean filtering mechanism is plausible when backgrounds are stable, it cannot filter out the strong fluctuations of naturalistic olfactory scenes (*Supp. Materials*, sec. 3), hinting that other mechanisms are at play.

Here, we propose a conceptually distinct model for olfactory habituation to broadly activating backgrounds that fluctuate on physically relevant timescales. As in previous proposals, we assume that background subtraction is mediated by plasticity mechanisms in inhibitory interneurons within the AL and OB. Our model is motivated by the fact that the representation of a fluctuating odor traverses a one-dimensional manifold (*i*.*e*., a curve) in a much higher-dimensional glomerular activity space [23]. Consequently, a mixture of backgrounds spans a manifold of dimensionality equal to the number of independently varying odors in the mixture. A network that learns this low-dimensional manifold can thereby subtract out background activity by projecting instantaneous activity to the low-dimensional background manifold, highlighting components that are orthogonal to it (Fig. 1D). Our proposed ‘manifold projection’ mechanism thus relies on the high-dimensional nature of olfactory coding, and is still applicable when odors fluctuate on timescales comparable to timescales of neural signal propagation. Hence, the characteristics of olfactory stimuli and circuits outline a distinct habituation mechanism at the level of neural population codes, which could also be leveraged by other biological circuits facing fluctuating backgrounds in high-dimensional input spaces.

The structure of the paper is as follows. We first use a minimal mathematical model to delineate the physical and sensory coding regimes where a manifold projection strategy outcompetes a predictive filtering mechanism for new signal recognition among strong background fluctuations. Next, we propose a biologically plausible model for manifold learning implemented by inhibitory interneurons in the early olfactory system. We consider two local plasticity rules (IBCM and BioPCA), which find the linear subspace spanned by the background odors. Interneurons equipped with either rule perform considerably better against fluctuating backgrounds than prior models and perform comparably against each other. A detailed mathematical analysis of the IBCM rule shows that interneurons acquire selectivity to independent odors in the mixture. Finally, we show that both plasticity rules are robust across a range of physiologically relevant physical and computational regimes.

## Results

### Regimes of predictive filtering and manifold learning

We begin by delineating the physical and computational regimes in which a *manifold learning* strategy for habituation outperforms a *predictive filtering* strategy. We consider an olfactory system habituating for time *T* to a fluctuating background, **b**(*t*). The components of these vectors represent the glomerular activations corresponding to each OR type, and thus reflect the coordinates of an odor in an *N*_S_-dimensional olfactory coding space. As in the rest of the paper, the backgrounds **b**(*t*) are mixtures of *N*_B_ odor vectors,

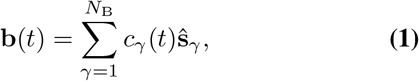

where the concentration of the *γ*th odor at time *t* is *c*_*γ*_(*t*). Here, we have assumed additive odor mixtures to keep our analysis tractable; our conceptual argument should also extend to non-additive odor mixture coding [17, 37–39] when combined with algorithms for curved manifolds (*Discussion*). The system subsequently responds at time *T* to a mixture of the fluctuating background **b**(*T*) and a new, behaviorally relevant target odor **s**_new_ which the target aims to recognize; the total input is **b**(*T*) + **s**_new_. An idealized circuit subtracts a vector **u**_*T*_ from the target-background mixture, where

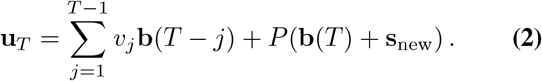

The first term represents a weighted average over the background’s history, corresponding to the predictive filtering strategy, where scalar coefficients *v*_*j*_ are set by the second-order statistics of background fluctuations. The second term corresponds to a simplified manifold learning strategy, where the matrix *P* projects the current stimulus to the subspace spanned by the background. The parameters *v*_*j*_ and *P* are learned during habituation such that the target odor **s**_new_ is recovered (in the mean squared error sense) by subtracting **u**_*T*_ from the current input.

We analytically optimized *v*_*j*_ and *P* to delineate how the two strategies contribute to recovering the target odor **s**_new_ from the mixture with **b**(*T*), which depends on background statistics (*Supp. Materials*, sec. 1). Fig. 1E illustrates the optimal reconstruction error for different autocorrelation timescales of background fluctuations (*τ*), and dimensionalities of the olfactory coding space (*N*_S_, Fig. S1A-B). While the combined strategy is by construction always better than each individual strategy, manifold learning alone explains all the performance when fluctuations are fast and the dimensionality is large (small *τ*, large *N*_S_).

The spectrum of turbulent fluctuations is dominated by brief whiffs and blanks that can reach down to ∼10 ms (Fig. 1B); predictive filtering, since it acts as a change detector, would constantly respond to these whiffs. Taking the filtering time step to be the smallest olfactory delay functionally perceptible to mice and human (30-60 ms) [40, 41], we estimate that *τ <* 100 for typical turbulent backgrounds (Fig. S1C). The number of OR types spans from a few tens to thousands across different animals. The olfactory space is high-dimensional compared to the typical number of independent odor sources that might prevail in a natural landscape. Thus, physics and neurobiology together indicate that olfaction lies within the regime where a manifold learning strategy is most effective.

### Models of manifold learning in the early olfactory circuit to improve new odor recognition

We now develop biologically plausible models of olfactory habituation that rely on manifold learning. Following [36, 42], we formulate a mathematical description (Fig. 2A) of the early olfactory circuit (Fig. 1C). We use *Drosophila* cell types for conciseness, but the model generalizes to other organisms. The key component for habituation is a layer of *N*_I_ lateral interneurons (LN) which receive inputs **s** from olfactory sensory neurons (OSN) via synaptic weights *M* and are coupled with lateral connections *L*, thus having activities 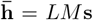. OSNs excite projection neurons (PNs) with unit synaptic weights and LNs inhibit PNs with synaptic weights *W*. The net activity of the PNs is thus 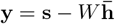.

**Fig. 2.**
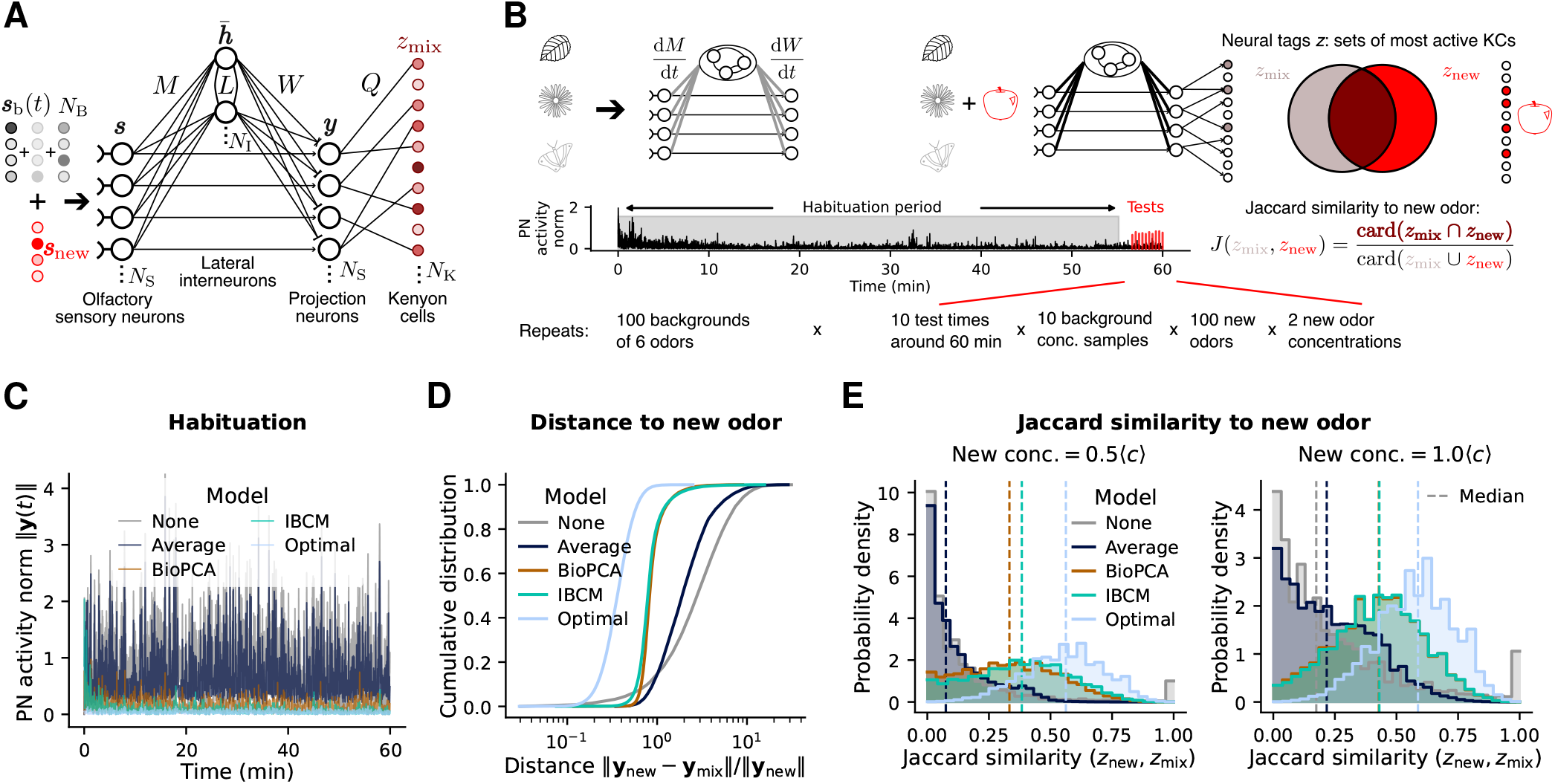
Recognition of new odors after habituation to a background with different learning models. (**A**) Mathematical description of the olfactory network. (**B**) Schematic of the numerical experiments performed to assess habituation performance. We generate a set of background odors randomly, then integrate the network’s synaptic plasticity equations for 60 minutes of habituation to a simulated background time series, where odor concentrations fluctuate according to the turbulent stochastic process illustrated in Fig. 1A-B (see *Supp. Materials* sec. 2 for simulation methods). We then compute the network’s response to a new odor **s**_new_ mixed with the background. The recognition performance is quantified by the Jaccard similarity between the neural tag of the mixture, *z*_mix_, and the neural tag (pre-habituation) of the new odor alone, *z*_new_. This procedure is repeated for several test times, random backgrounds, samples of each background, random new odors, and different new odor concentrations, for each habituation model (none, average subtraction, BioPCA, IBCM). Parameter values are listed in the *Methods*. (**C**) Sample time series of the norm of PN activity, to illustrate the extent of habituation (decrease in PN activity when exposed to the background) in each model. “Optimal”: response with the optimal manifold learning matrix *P* (no predictive filtering) derived for Fig. 1E. (**D**) Cumulative distribution of the Euclidean distance between new odors **s**_new_ and each model’s PN response to new odors mixed with the background, **y**_mix_, after habituation, across all background, odors, and new odor concentrations tested. (**E**) Distribution of Jaccard similarities *J*(*z*_n_, *z*_mix_) of the various models, across all backgrounds and odor samples tested.

Intuitively, in our model, interneurons learn to project inputs onto the low-dimensional subspace of background odors, and subtract these projections from PNs. Interneurons can therefore perform (linear) manifold learning with projection matrix *W LM*. Lastly, PN activities are projected on a large layer of *N*_K_ Kenyon cells (KC) by sparse random connections (fixed, not learned). The condition *N*_K_ ≫ *N*_S_ ensures that the 5 % most active KCs represent a distinct neural tag *z* for each possible odor. This dimensional expansion from PNs to KCs implements locality-sensitive hashing of input identity [42].

Next, we consider biologically realistic synaptic plasticity rules for weights *M, L*, and *W* to achieve adequate manifold learning within this network. The optimal manifold projection matrix *P* derived for Fig. 1E involves non-local terms and moments of the new odor distribution inaccessible to the network (eq. 9). Instead, we postulate a simple, unsupervised, local learning rule for the inhibitory weights *W* : they evolve during habituation to minimize the norm of the PN activity **y** by using interneuron activities 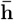. This optimization principle results in simple Hebbian dynamics (see *Methods*),

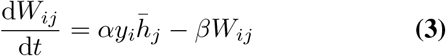

with learning rate *α* and a regularization rate *β*. This simple update rule allows us to compare different models for the projection weights *M* and *L*. We consider two models: (a) the Intrator, Bienenstock, Cooper, and Munro (IBCM) model of synaptic plasticity [43–46], and (b) a biologically plausible online implementation of principal components analysis (BioPCA) [47] (for full model definitions, see *Methods*). The IBCM model was proposed to explain neuronal selectivity to specific stimulus components [43]. Its connections with independent component analysis (ICA) [48, 49] suggest that it could provide a biologically meaningful basis for learning the manifold of non-Gaussian backgrounds.

We perform initial numerical simulations, outlined in Fig. 2B, to assess the performance of different habituation schemes. We compare these rules to the average subtraction mechanism proposed in [36] (“Average”), as well as with the absence of habituation (“None”) and the optimal manifold learning matrix *P* derived in the previous section (see *Methods*). The dynamical equations for each plasticity rule are integrated in the presence of a background as in Eq. (1), with concentrations fluctuating according to the turbulent stochastic process of Fig. 1A-B. After this habituation period, we present the network with mixtures of the background and new odors, **s**_mix_ = **s**_b_(*t*) + **s**_new_. To assess how well that odor is decoded from the mixture, we compare its output with the neural tag *z*_new_ of the new odor alone.

We find that while average subtraction cannot inhibit the strong fluctuations of turbulent backgrounds (*Supp. Materials*, sec. 3), both the IBCM and BioPCA networks significantly reduce PN activity in response to the background (Fig. 2C) comparably to the optimal manifold learning matrix *P*. This confirms that both models provide adequate projections on the background subspace, allowing the Hebbian rule for *W* (Eq. (3)) to achieve its function of minimizing PN activity.

Moreover, we compare the models’ performance for new odor recognition after habituation at the level of PN activities (Fig. 2D) and neural tags (Fig. 2E). For both metrics, average subtraction provides a very limited improvement compared to recognition without habituation, due to the strong background fluctuations caused by turbulence. In contrast, manifold learning implementations significantly improve odor recognition, with both IBCM and BioPCA networks performing similarly well. With respect to the distance in PN activity between the new odor alone and mixed with the background (Fig. 2D), these models result in a ~ 3-fold improvement, but fall short of the optimum by a similar factor. This is not surprising, since *P* is fine-tuned for the distribution of new odors, which is unknown to the IBCM and BioPCA networks. Nonetheless, in terms of the Jaccard similarity between neural tags of the mixture and the new odor (Fig. 2B, right), the IBCM and BioPCA networks perform within 15 % similarity of the optimum (Fig. 2E), producing responses much more similar to the new odor than to background odors (Fig. S2). These models thus recover, from a mixture with the background, roughly 50 % of the KCs which are most activated by the new odor alone and define its identity, even when the new odor is present at just half the average whiff concentration (Fig. 2E, left). These results prompt us to investigate in more detail how the IBCM and BioPCA models learn the background manifold.

### Analysis of background habituation by the IBCM model

We first focus on the IBCM model, since its mechanisms are less intuitive than PCA and have classically been characterized for visual input processes alternating between a fixed set of vectors [44, 50]. In its simplest form (Fig. 3A), the IBCM model describes the slow variation of a neuron’s synaptic input weights **m** (a row in matrix *M*) as

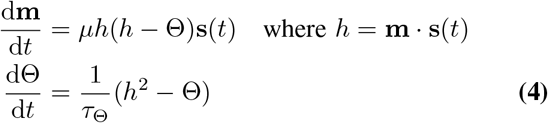

where *h* is the activity in response to input **s**(*t*), *µ* is the learning rate, and Θ is an internal threshold converging to a temporal average Θ = ⟨ *h*^2^⟩ (⟨ *·* ⟩ denotes averages over the input fluctuations). In practice, we include lateral mean-field inhibition between interneurons with coupling parameter *η*, a mild nonlinearity to *h* preventing excessively large activations, and a small − (*ε ≪* 1) decay term *εµ***m**. For simulations with turbulent backgrounds, we use a variant of the model from [45] where the learning rate is divided by Θ to speed up convergence (see *Methods*).

**Fig. 3.**
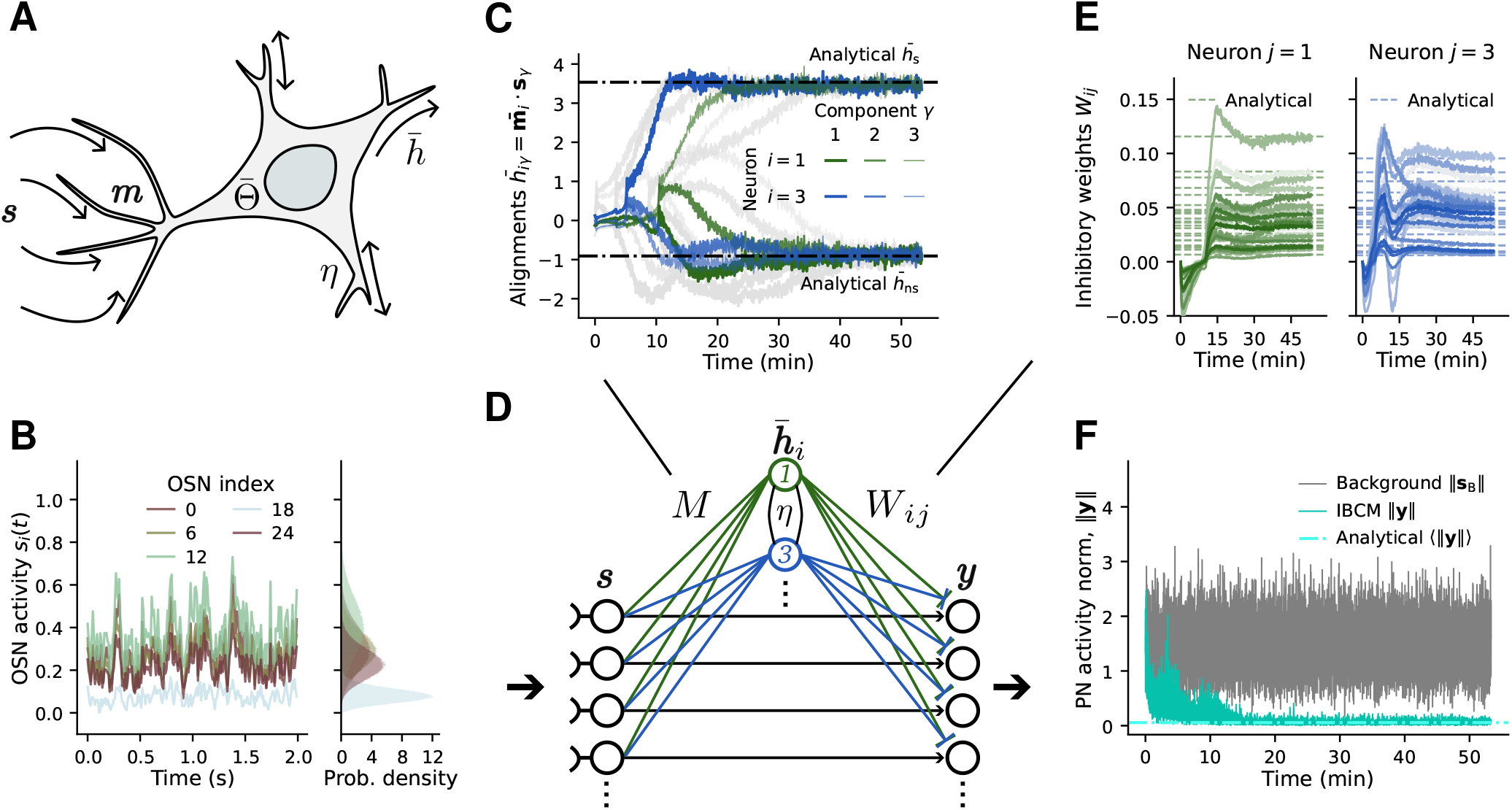
The IBCM model of synaptic plasticity learns olfactory background components. (**A**) Illustration of an IBCM neuron’s inputs from OSNs, **s**, synaptic weights **m**, activity 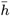, and internal threshold 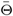. (**B**) OSN activities for a simple background process with 3 odors and weakly non-Gaussian concentration fluctuations. (**C**) Time series of the IBCM neurons’ alignment with the different background odors (two neurons are highlighted in green and blue, with line widths indicating different odors *γ*). Each neuron predominantly aligns with one odor (large dot product value 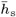 for one odor, small value 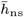 for the others). This steady-state alignment matches our analytical predictions (dashed lines, *Supp. Materials* sec. 4). (**D**) Location, in the olfactory network model, of the different weights and neurons illustrated in other panels. (**E**) Time series of the IBCM neurons’ inhibitory synaptic weights, *W*, for the neurons highlighted in (C). The steady-state values of these weights can also be derived analytically (dashed lines) and align well with the simulations. (**F**) Norm of the response of projection neurons (PN) to the background process during habituation. The analytical prediction (dashed line) over-estimates the actual inhibition, since it neglects the contribution of fluctuations in *M* and *W*.

The **m** equation introduces a competition between input patterns: inputs that cause *h >* Θ are further reinforced, while sub-threshold ones are further depressed; consequently, a neuron responds specifically to some inputs and does not respond to others [43]. This mechanism works when the threshold time scale *τ*_Θ_ is slow enough to average over fast input fluctuations **s**(*t*), yet still fast compared to the learning rate *µ*. This separation of time scales is to ensure **m** does not vary much while Θ averages over fluctuations; oscillations arise in the synaptic weights [51] without the separation *τ*_Θ_ ≪ 1*/µ*. This criterion can nonetheless be achieved during olfactory habituation over the course of 30-60 minutes.

To gain insight into how IBCM neurons learn the background subspace, we examine the fixed point equations of the model averaged over fast input fluctuations (*Supp. Materials*, sec. 4), finding exact expressions for these solutions in terms of the background concentration moments. From a linear stability analysis (*Supp. Materials*, sec. 4F and Fig. S3), we find that the only stable fixed points are those where the alignment of the synaptic weights with background odor vectors, 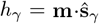, take a large positive value *h*_sp_ (“specific”) for *one* odor *γ* and a small, possibly negative value *h*_ns_ (“non-specific”) for all other background components. Hence, an IBCM neuron learns to selectively respond to one background odor: the classical specificity property of this model [44] thus extends to quite general input stochastic processes of the form given in eq. 1. From the solutions for **m** and **h**, we also derive analytical expressions for the fluctuation-averaged inhibitory *W* weights at steady-state (*Supp. Materials*, sec. 4G). Overall, our results show that a network of IBCM neurons performs manifold learning by having each neuron selectively suppress one background odor. Lateral inhibitory coupling between IBCM neurons (matrix *L*) help to push each neuron towards a different odor component [52].

To confirm our analysis, we perform numerical simulations with a simpler, weakly non-Gaussian background (Fig. 3B, see *Methods*). As expected, each IBCM neuron evolves over time to align with one background component (Fig. 3C,D), with steady-state average values of the dot products 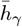 closely matching our analytical predictions *h*_sp_, *h*_ns_ (dashed lines). Different IBCM neurons (labeled by colors) become specific to different odors (line transparencies). The *W* weights also converge to steady-state average values matching our analytical results (Fig. 3E). The network of IBCM neurons performs habituation effectively, reducing both the mean and standard deviation (fluctuations) of the PN activity norm below ∼ 10% of the input levels. Characterizing further the learning dynamics, we observe that the selectivity of IBCM neurons is acquired in two phases, first approaching a saddle point before converging to a selective fixed point (*Supp. Materials* sec. 6, Fig. S4, and Fig. 3C). This selectivity is driven by skewness (non-zero third moment) in the background statistics [44, 53] (Fig. S5).

Importantly, these properties of IBCM neurons are robust across different background statistics. We observe similar specificity and learning dynamics in IBCM neurons exposed to log-normal background fluctuations (Fig. S6) and in the simulations of Fig. 2 with the full turbulent statistics (Fig. 4A-B). In the latter case, averaging over long whiffs and blanks requires slower learning rates 1*/τ*_Θ_ and *µ* (see *Methods*and Table S2); despite stronger fluctuations, IBCM neurons align with a single background odor each.

**Fig. 4.**
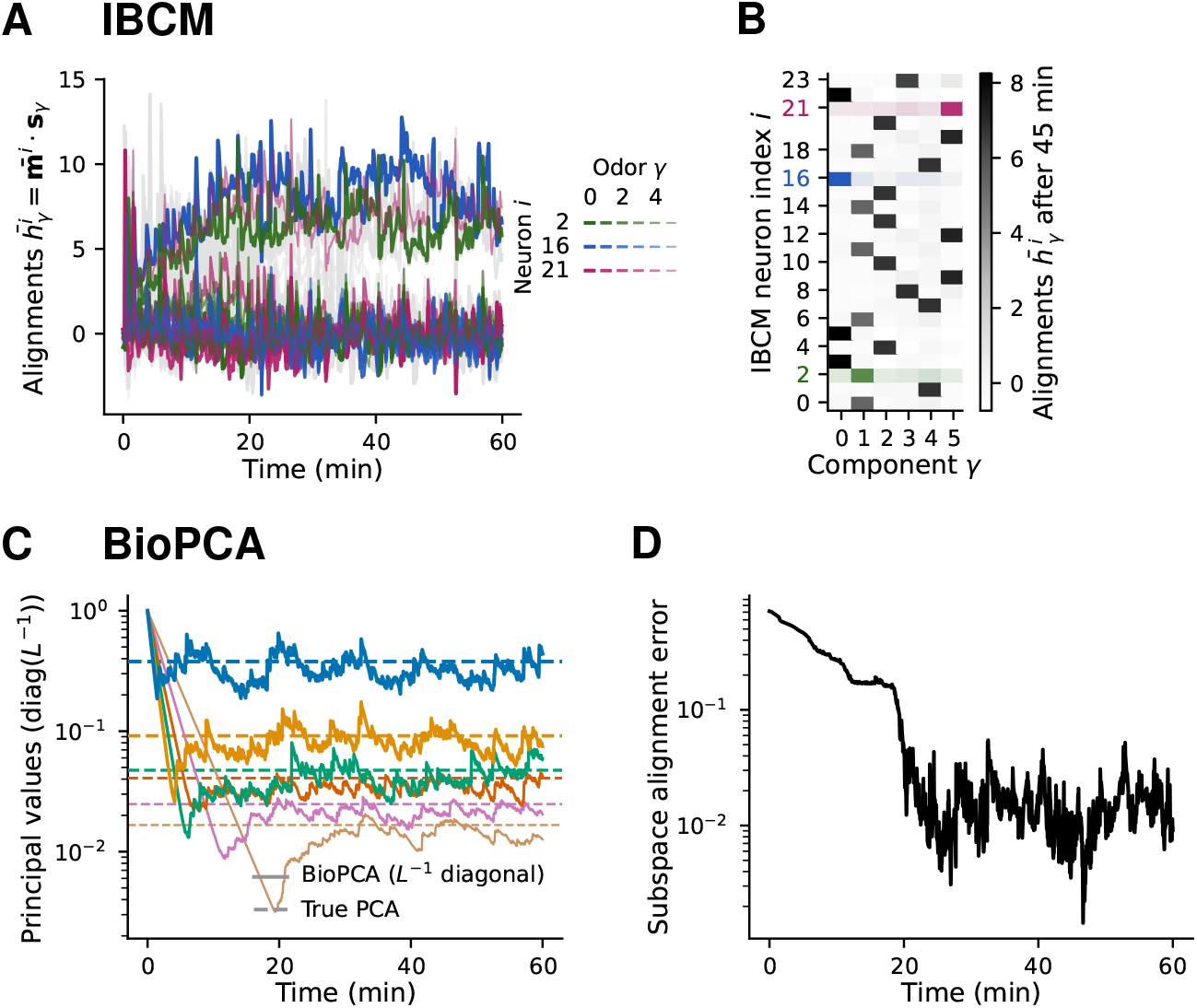
Habituation of IBCM and BioPCA neurons to turbulent olfactory backgrounds. (**A**) Time series of the IBCM neurons’ synaptic weight alignment with background odor during habituation to a six-odor background with the turbulent concentration stochastic process illustrated in Fig. 1A-B. Three neurons are highlighted with colors. (**B**) Table of each neuron’s alignment after habituation, showing that IBCM neurons becomes selective for one odor even in this strongly fluctuating background. (**C**) Time series of the principal values learned by lateral interneurons obeying the BioPCA model during habituation to the same turbulent background. These principal values are stored in the inverse of the self-coupling weights 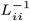 (inverse of the diagonal entries in the LN coupling matrix *L*). The principal values learned by the first *N*_*B*_ neurons converge to averages equal to the *N*_*B*_ non-zero eigenvalues of a PCA decomposition of the background (dashed horizontal lines). (**D**) Alignment error between the background subspace and the principal components learned by the BioPCA LNs (the rows of the *LM* matrix should be the principal components), confirming the model does learn the PCA decomposition of the background.

### Comparison of IBCM and BioPCA learning

As a point of comparison, we also analyze how the BioPCA network learns the background manifold in fluctuating environments. Despite strongly non-Gaussian statistics, the network converges to the expected PCA decomposition fixed point (see *Methods, Supp. Materials*, sec. 5 and sec. 6C). The matrix *L* becomes nearly diagonal, containing the principal values (Fig. 4C, Fig. S6B), while the rows of *M* converge to (scaled versions of) the principal component vectors, as evidenced by the error on their alignment decreasing to ∼ 1 % (Fig. 4D, Fig. S6C, Fig. S7).

On the whole, both models achieve similar habituation levels within ∼ 30 minutes. However, they rely on distinct mechanisms and converge to different vector bases for the background manifold. BioPCA neurons learn principal components: linear combinations of the true odors, distinguished by their variance. IBCM neurons, in contrast, rely on higher statistical moments of the inputs to select individual odor sources. Both models require in principle one neuron per background dimension, *N*_I_ = *N*_B_, to span the background subspace. Superfluous neurons have little effect for the BioPCA model, where they reach principal values *L*_*ii*_ ≈ 0. For the IBCM network, extra neurons are helpful, increasing the probability that each background odor will be selected by at least one of them. Notwithstanding these differences, both models produce very similar habituation and odor recognition performances in Fig. 2. They are also similar in their robustness to OSN noise (Fig. S8), and in their performance when combined with alternate Hebbian rules for the *W* weights based on different *L*^*P*^ norms (*Supp. Materials* sec. 7 and Fig. S9).

### Performance in various olfactory space conditions

To understand the similar performance of the IBCM and BioPCA versions of manifold learning, we investigate the effect of various olfactory space parameters on them. We perform numerical simulations analogous to those of Fig. 2B for increasingly large olfactory space dimensions (*N*_S_) and for a wider range of new odor concentrations (Fig. 5A). We consider dimensionalities ranging from half (25) that of the fruit fly (50) up to human (300) and mouse (1000) levels. While the performance of the optimal manifold learning algorithm increases with *N*_S_ up to a nearly perfect score, the IBCM and BioPCA networks reach a very similar plateau at *N*_S_ ∼ 100 (Fig. 5B). Remarkably, this plateau corresponds to the similarity between the new odor tag, *z*_new_, and the tag *z*_new,⊥_ of the new odor component orthogonal to the background, **y**_new,_ (“orthogonal” pink line, Fig. 5B). This observation clarifies why both models perform similarly well: the local rules for *W* (Eq. (3)), based on minimizing PN activity, cause the parallel component of the new odor to be subtracted at the same time as the background. As long as the *M* weights provide complete projections of the inputs, the network produces a response **y**_mix_ ≈ **s**_new,⊥_. In comparison, the optimal matrix *P* preserves some of the new odor’s parallel component, thus maximizing the recognition of **s**_new_.

**Fig. 5.**
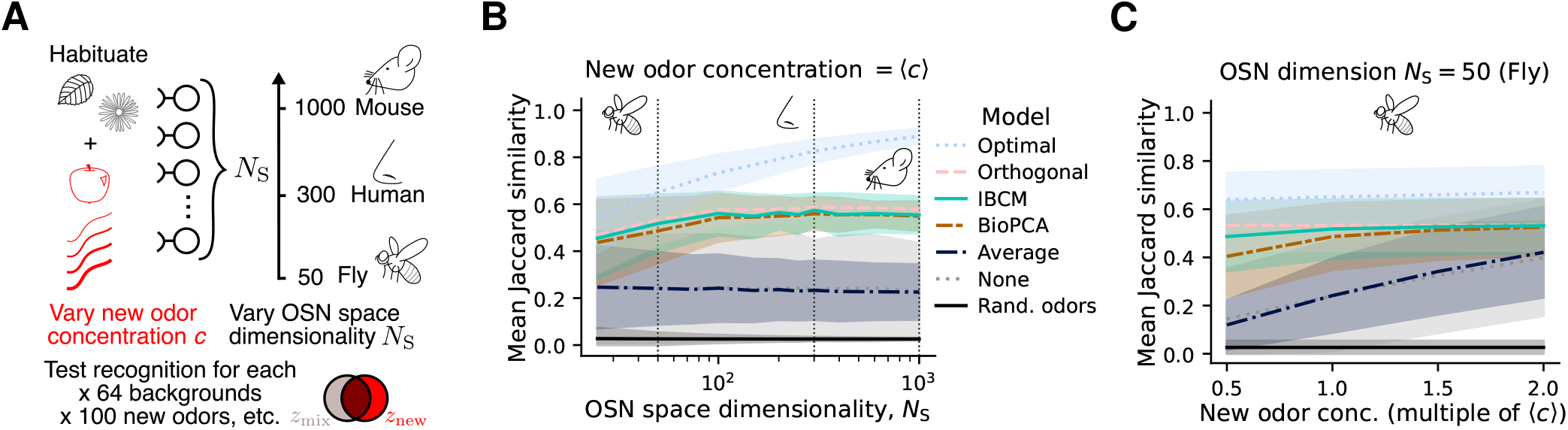
Recognition performance as a function of olfactory space dimensionality and new odor concentration. (**A**) For various olfactory space dimensionalities *N*_S_ (*i*.*e*., number of OSN types) and new odor concentrations *c*_n_ at test times, we perform simulations like those in Fig. 2: the model habituates to a six-odor turbulent background for ∼60 minutes before a new odor is introduced in the mixture. Each (*N*_S_, *c*_new_) condition is tested across 64 backgrounds, 100 new odors, 5 test times post-habituation, and 4 background samples at each test time. (**B**) New odor recognition performance, quantified by the Jaccard similarity between the new odor and the response to the mixture after habituation, as a function of *N*_S_, for different manifold learning models. Results shown here are for the new odor *c*_new_ equal to the average concentration of background odors, ⟨ *c* ⟩ (see Fig. S10 for all concentrations). “Optimal”: manifold learning matrix *W* derived in Fig. 1E. “Orthogonal”: similarity between the entire new odor and its component orthogonal to the background. “Rand. odors”: similarity between two randomly selected odors (*i*.*e*., similarity by chance). The shaded area represents one standard deviation across replicates. (**C**) Recognition performance as a function of the concentration at which the new odor is presented (multiples of ⟨ *c*⟩), for the fly case (*N*_S_ = 50; see *Supp. Materials*, Fig. S10A for all dimensions). Same legend as (B).

Still, the levels of odor recognition reached by the IBCM and BioPCA models are significant, several standard deviations above chance similarity (black line, Fig. 5B). They also perform well above the average subtraction model, which was similar to no habituation: in that model, new odor signals are masked by strong fluctuations away from the average. For higher new odor concentrations, manifold learning provides a more modest improvement (Fig. 5C and Fig. S10), because habituation is not as crucial for very strong new odor whiffs which dominate the background. Overall, both IBCM and BioPCA interneurons subtract the background manifold from olfactory inputs while preserving new odor signals in regimes that are physically and computationally relevant for biological systems.

## Discussion

Sensory adaptation to olfactory backgrounds is particularly challenging due to strong fluctuations generated by turbulent mixing in naturalistic conditions. We showed that predictive filtering strategies, which act on individual stimulus features, cannot adequately distinguish between changes in activity due to new odors and changes in activity due to fluctuations in the background. An alternative class of habituation strategies, manifold learning, could better identify new odors by learning to subtract projections of the instantaneous inputs onto the low-dimensional background manifold. We propose that inhibitory interneurons, which modulate the activity of principal neurons in early olfactory pathways, implement a manifold learning strategy for habituation. We explore two classes of synaptic plasticity rules, each of which combines a Hebbian-like rule and a linear projection learning rule (IBCM or BioPCA). Our analysis shows that these simplified linear manifold learning strategies are near-optimal for a range of physiologically relevant parameters, including when background odors display strong fluctuations such as those encountered in turbulent environments. Both plasticity rules show comparable performance on a habituation task, but learn distinct stimulus features. Notably, IBCM neurons select biologically relevant projections corresponding to independently varying components in the background mixture.

The biological underpinnings of our proposed model for olfactory background manifold projection are supported by previous experimental and theoretical studies. All connections in our network structure (OSN to PN, OSN to LN, LN to PN) are abundant in the connectome [12]. Habituation on the time scale of minutes has been shown to occur predominantly at the level of PNs in flies [22, 54] or M/T cells in mice [29]. Several studies found lateral inhibitory signals (GABA, glutamate) and their receptors for such signals (GABA-A, NDMA) to be essential to habituation [6, 27, 34, 35]. Our model relies on odor-specific PN inhibition by PN-to-LN plasticity for habituation, as observed in *Drosophila* [8] and honeybees [55]. Of note, we neglected feedback of PNs on LNs [31] and instead considered a feed-forward network for mathematical simplicity, to illustrate our concept of manifold learning. Moreover, recent theoretical work has argued that the PN-LN connectivity pattern reflects correlations in PN activity, suggesting that the PN-LN circuit whitens odor representations in the antennal lobe [56]. However, the authors focused on hardwired computations pre-adapted to a given set of odors (*i*.*e*., offline), whereas we addressed a different problem altogether, showing that online PCA is one plausible set of synaptic plasticity rules to achieve background subtraction in fluctuating environments.

Our theory provides salient predictions that could be tested experimentally. The simplest observable feature is the decrease in both the mean and variance of PN (or M/T cells in mice) activity after 20-60 minutes of exposure to turbulent odor mixtures (Fig. 2C and 3F). This phenomenon has already been observed for simpler backgrounds in *Drosophila* [8], which motivated our study. It could be directly tested in mice by calcium fluorescence imaging of glomeruli [37]. PN or glomerular activity should however be restored in response to new odors orthogonal to the learned background. In comparison, temporal average filtering would fail to reduce PN activity (Fig. 2C), while filtering based on recent samples (as in eq. 2) would rapidly suppress the response to new odors as well.

A more subtle feature in our proposed model is that lateral interneuron activity (LN) should, conversely, closely track background stimuli in real time to keep inhibiting PN responses (Fig. S8D-E). It may be experimentally challenging, however, to single out interneurons and record their fast fluctuations. A corollary of this model feature (which might be easier to measure) is that, since LN activity reflects projections on the learned background manifold, these neurons should become silent if the stimulus is suddenly switched to new odors with null projections on the previous subspace. Their activity should slowly recover on the time scale of habituation as they learn the new background.

A third feature of habituation by manifold learning is that new odor recognition performance decreases with the distance to the background subspace (*i*.*e*., as the norm of the orthogonal component **s**_n,⊥_ decreases; Fig. S2C-D). This dependence would not be as strong in predictive filtering strategies. This correlation between odor recognition and distance from new odors could be tested in behavioral experiments.

Further theoretical work will also be necessary to refine our proposed implementation of manifold learning in olfactory circuits, and to assess how this strategy may be coupled with other odor recognition mechanisms. Schemes more sophisticated than a Hebbian rule would be necessary to reach the optimal performance promised by manifold learning (Figure 5) or to fully exploit the biologically relevant projections learned by IBCM neurons. Also, in our study, we focused on linear background manifolds (eq. 1) that could be decomposed into linear projections (**h** = *LM* **s**); while manifold projection also applies conceptually to non-additive odor mixtures, this extension will require olfactory circuit implementations of algorithms for curved manifold learning, such as manifold tiling [57]. Moreover, by choosing to focus on early olfactory processing, we neglected neuromodulatory inputs [58–60] and feedbacks from higher-level cognitive functions [61, 62]. For instance, while we assumed background and new odors are merely defined by their order of presentation, long-term memory of odors and other computations in the piriform cortex [63] likely help mammals focus their attention on relevant cues rather than on uninformative odors for,*e*.*g*., odor trail tracking [64, 65]. Future investigation on this aspect could draw upon recent advances on attention mechanisms in artificial learning models [66]. Conversely, the concept of background manifold projection could prove useful for algorithms performing figure-ground segregation in time-varying signals, such as in video object detection [67].

Beyond olfaction, the interplay between habituation and attention also arises in other biological systems performing chemodetection in fluctuating environments [68]. For instance, in T cell antigen recognition [69], both (immune) memory and (T cell receptor) signal processing networks play important roles for pathogen detection amid a sea of irrelevant (self) antigens. Overall, we hope that our proposed model of habituation via manifold learning will motivate further theoretical and experimental efforts to clarify how living systems meet the challenge of adaptation to fluctuating backgrounds.

## Materials and Methods

### Odor vectors and concentrations

In our models, odors have a fixed, unit-normed direction, and an amplitude along that axis set by their (fluctuating) concentration: **s**(*t*) = *c***ŝ**. Except for the idealized setup of Fig. 1E, vectors for background (**ŝ**_*γ*_) and new (**ŝ**^new^) odors are drawn from the same distribution 𝒫_**ŝ**_, by sampling i.i.d. exponential elements, then normalizing each vector. New odors are tested at fixed concentrations *c*_new_. Background concentrations *c*_*γ*_ (*t*) follow a stochastic process, usually (Figs. 2, 5) the turbulent process illustrated in Fig. 1A-B. We simulate each odor concentration as a telegraph-like process, alternating blanks and whiffs with stochastic durations and whiff concentrations. The power-law distribution of whiffs and blanks durations (*t*_*w*_, *t*_*b*_) have a lower cutoff at 10 ms and upper cutoffs at 5 s (whiffs) or 8 s (blanks), respectively. The whiff concentration distribution has a scale *c*_0_ = 0.6. We also considered weakly non-Gaussian (Fig. 3) and log-normal (Fig. S6) background concentrations, by simulating a multivariate Ornstein-Uhlenbeck *{g*_*γ*_ (*t*)*}*, then transforming these variables as *c*_*γ*_ (*t*) = *g*_*γ*_ (*t*) + *νg*_*γ*_ (*t*)^2^ or 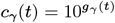, respectively. We used a short autocorrelation time *τ* = 20 ms, an average 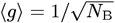, and standard deviation *σ*_*g*_ = 0.3. For the non-Gaussian case, we set *ν* = 0.2. Details of the stochastic simulation methods are provided in *Supp. Materials* sec. 2.

### Optimizing predictive filtering and manifold learning regimes

In Eq. (2), we introduced an idealized inhibitory network response combining manifold learning and predictive filtering. The objective of this network is to minimize the squared distance between its response, **b**(*T*) + **s**_n_ − **u**(*T*), and the target odor alone, **s**^n^. The corresponding loss is

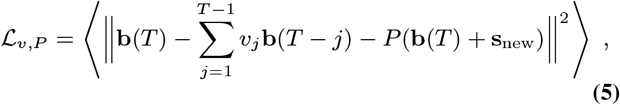

where the average is taken across samples (concentrations) from a given background and across new odors. Our goal was to determine the ideal performance of such a network, and the contribution of each habituation strategy depending on olfactory space parameters. We therefore solved for the optimal scalar coefficients *v*^*j*^ and the optimal *N*_S_ *× N*_S_ matrix *P*, as an upper bound on the mechanisms that real networks could learn during habituation. We minimized the loss by solving 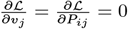. *Supp. Materials* sec. 1 details the calculation and the result; we obtained a general solution for any background mixture, as defined in Eq. (1), considering zero-mean and statistically independent concentrations.

For Fig. 1E, we considered a simpler particular case where background odors are orthogonal, new odors are drawn uniformly from the unit hypersphere, and background concentrations have an exponential autocorrelation function with time constant *τ*, ⟨ *c*_*γ*_ (*t*)*c*_*ρ*_(*t* + *s*) ⟩ = *σ*^2^*δ*_*γρ*_*e*−^|*s*|*/τ*^. The minimized loss is then

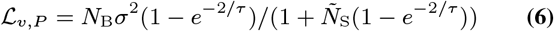

where 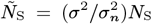 and 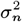 is the new odor concentration variance. In the figure, we compared ℒ_*v,P*_ with the limiting cases of pure predictive filtering (ℒ_*v*_, setting *P* = 0) and pure manifold learning (ℒ_*P*_, setting *v*_*j*_ = 0), which respectively give losses

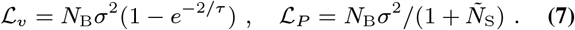

For the “optimal” strategy in Figures 2 and 5, we needed to derive the general solution (for non-orthogonal background and new odors drawn from 𝒫_**ŝ**_) for pure manifold learning (*v* = 0) when the background has a non-zero average (as was the case in our simulations). In that case, the optimal projection matrix is

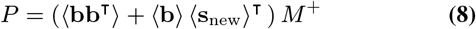

where *M*^+^ is the Moore-Penrose pseudo-inverse (or the usual matrix inverse, when it exists) of

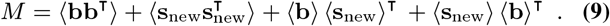

We evaluated numerically the moments ⟨**bb**^T^⟩, 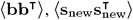, etc., by sampling unit vectors **s**_*γ*_, **s**_new_ from 𝒫_**ŝ**_ and background concentrations *c*_*γ*_ from the stationary distribution of the background process at hand – for Figs. 2 and 5, the turbulent statistics shown in Fig. 1A-B.

### Mathematical model of the olfactory network

We model the instantaneous response of the olfactory network (Fig. 2A) to a stimulus *s*(*t*) received at time *t* as the following set of neural activities in its different layers:

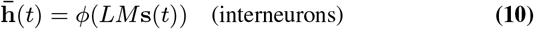

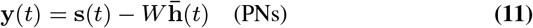

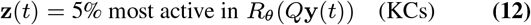

where *ϕ* is an element-wise nonlinearity and *R*_*θ*_ clips elements below threshold *θ*. We used *ϕ*(*x*) = *A*_sat_ tanh(*x/A*_sat_) for IBCM neurons to saturate their activity at a large *A*_sat_ = 50 for numerical stability; most of the time, *x ≪ A*_sat_ is in the linear part of this function, so **y ≈ s** − *WLM* **s**. We did not apply a nonlinearity for the BioPCA network, nor for the IBCM model on simpler backgrounds (Fig. 3).

Then, the neural tag *z* is computed as in [36]. First, PN activities are projected to the KC layer by the sparse *N*_K_ *× N*_S_ binary matrix *Q*. Then, *R*_*θ*_ clips Kenyon cells (KCs) with activity below threshold; we set 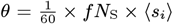, where ⟨ *s*_*i*_ is the average OSN activity in the current input **s**(*t*) and *f* = 6*/*50, the fraction of PNs forming a synapse with one KC in *Drosophila*. Finally, the neural tag *z* is the set of all non-zero KCs with activity above the 95th percentile of all KC activities.

The matrix *Q* is generated by randomly picking *f N*_S_ PNs to project to each KC (*i*.*e*., picking *f N*_S_ non-zero elements in *Q*’s row for that KC). We generated a new *Q* for each background tested in the numerical experiments in Figs. 2 and 5. When varying the olfactory space dimensionality *N*_S_ in Fig. 5, we preserved the relative size of PN and KC layers *N*_S_*/N*_K_ = 50*/*2000 found in *Drosophila*; hence, for mice with *N*_S_ = 1000 OR types, we used *N*_K_ = 40, 000 cortical cells (KC equivalent), and the *Q* matrix had *f N*_S_ = 120 M/T cells (PN equivalent) projecting to each cortical cell. This ratio aligned with experimental estimates in mice giving ∼200 glomeruli connected to a cortical cell, or 10 % sparsity in *Q* [70–72]. The other matrices (*M, W, L* in BioPCA) are slowly updated according to synaptic plasticity rules during a habituation run.

### Hebbian learning rule for *W*

The *W* weights are learned according to the Hebbian rule in Eq. (3). This rule derives from minimizing the average squared PN activity with *L*^2^ regularization on the *W*_*ij*_ :

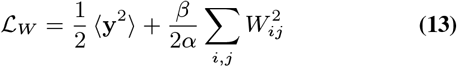

where we recall that 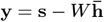. Taking the *W* dynamics to be a gradient descent on ℒ_*w*_ with rate 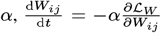, yields the aforementioned Hebbian rule. The average ⟨⟩ is replaced by time averaging over a time window 1*/α* using a slow rate *α* to implement online averaging of fast background fluctuations.

### Average subtraction model

The “negative image” subtraction model proposed in [36] is effectively a mean filtering or average subtraction model. It can be recast in the form of our network structure by having *N*_I_ = 1 interneuron with fixed activity *h* = 1, without *M* or *L* weights. The *W* weights are then a vector **w**_avg_, which is subtracted from the PN response input since *h* = 1: **y**(*t*) = **s**(*t*) − **w**_avg_. The Hebbian rule above is then

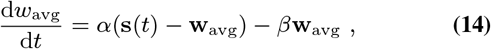

which makes **w**^avg^ align, at steady-state, with the average of the background over a time window, 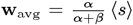 (*Supp. Materials*, sec. 3 for details).

### Network of IBCM neurons

Equation 4 presents the simplest form of the IBCM model for a single neuron. In our olfactory network, we consider *N*_I_ IBCM neurons with constant mean-field lateral inhibitory coupling, as proposed in [44], corresponding here to a matrix *L* with 1 on the diagonal and −*η* off-diagonal. Consequently, the reduced activity 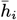 of neuron *i* (*i*.*e*., element *i* of 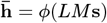) is

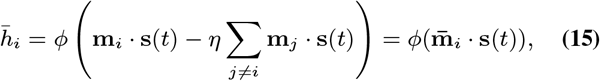

where we defined the inhibited weights 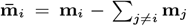, and where *ϕ*is the tanh nonlinearity introduced in Eq. (10). The complete dynamical equation for the synaptic weights **m**_*i*_ incoming into IBCM neuron *i* has additional terms due to this coupling,

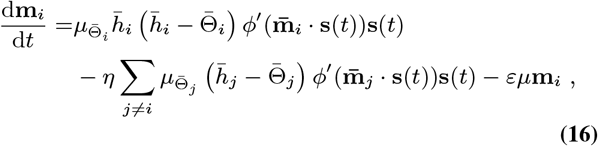

where *ϕ*^′^ is the derivative of the nonlinearity. We have also added a small decay term −*εµ***m**^*i*^ to eliminate any component orthogonal to the background manifold in the random initial weights. For simulations with turbulent background statistics, we scaled the learning rate in the first two terms as

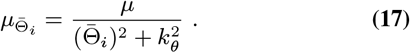

This form is similar to the variant introduced in [45], but we added a constant *k*_Θ_ in the denominator to prevent blowups at *t* = 0, where we initialize 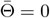. For simpler backgrounds (Figs. 3, Fig. S6), we did not include this variant and simply used *µ*_Θ_ = *µ*. Moreover, the internal threshold of each neuron, 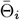, evolves as

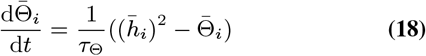

such that it tracks the reduced neuron activity 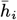 averaged on an intermediate time scale *τ*_Θ_.

### Network of BioPCA neurons

We could use the “inverse-free PSP” version of the biologically plausible online PCA (BioPCA) proposed in [47] directly for the *M* and *L* weights of the interneuron layer. The model converges to a fixed point where the *L* matrix is diagonal with the principal values in it, and where the matrix *LM* contains the principal vectors in its rows, with norms specified by the pre-defined diagonal matrix <u>Λ</u> [47, Lemma 3]. The model specifies dynamical update rules for *M* and *L*^*′*^ = *L*^1^, the inverse of *L*, rather than *L* directly. To avoid non-biological matrix inverse comnputations, the vector of interneuron activities **h** is computed with a Taylor series for *L*,

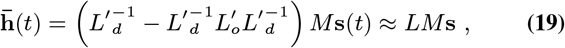

where *L*^′^_*d*_ contains the diagonal of *L*^′^, and *L*^′^_*d*_ contains the off-diagonal terms. This approximation is accurate at the fixed point where *L*^′^ is diagonal 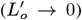. The BioPCA dynamical update rules converging to this PCA decomposition are

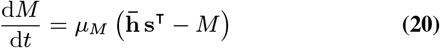

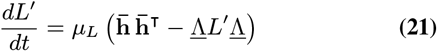

In practice, we set 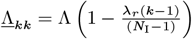, where Λ is the scaling factor for *M* weights described below to make IBCM and BioPCA perform similarly, and *λ*_*r*_ is the range of Λ values (between 0 and 1). We followed the original paper’s recommendation for the linear decrease of Λ_*kk*_ with *k* and for setting *µ*_*L*_ = 2*µ*_*M*_ For further comparison with the original paper [47], note the following equivalence between our notation theirs: *M ↔ W, L ↔ M*^−1^, **s***↔* **x**, and 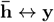.

In our simulations, we added an extra interneuron applying the average subtraction mechanism described in Eq. (14), with *β* = 0, upstream of the BioPCA model. This way, the BioPCA network learned the decomposition of the covariance matrix rather than of ⟨**ss**^T^⟩, which still includes the average ⟨ *s* ⟩. This choice did not change the model performance, but made it more interpretable.

To measure the convergence of *M* ‘s columns to the PCA vectors, in Fig. 4D, we computed, at each time point, the subspace alignment error proposed by [47],

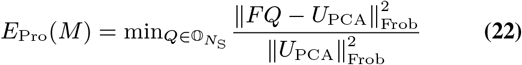

where columns of *U*_PCA_ contains the *N*_B_ PCA vectors with non-zero eigenvalues, *F* = (Λ^1^*LM*)^T^contains the eigenvectors learned in the network’s projection weights, and 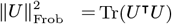 is the Frobenius matrix norm. The *Q* matrix minimizing the distance to give the alignment error solves the so-called orthogonal Procrustes problem and is *Q* = *U*_*F*_ *V*^T^ where *U*_*F*_, *V* come from the SVD of *U*_PCA_*F*^T^ = *U*_*F*_ Σ*V*^T^ [73].

### Scaling parameter Λ for M weights

In the BioPCA model, the scale parameter Λ in the <u>Λ</u> matrix (see below eq. 21) controls the magnitude of weights *M*. This scale influences the strength of habituation: larger *M* ∼ Λ weights allow smaller *W* ∼ 1*/*Λ weights that are less constrained by regularization (*β* term in eq. 13) and thus further reduce PN activity. We set Λ to the value necessary to achieve the same background reduction level as the IBCM network, as predicted by our analytical calculations for the IBCM fixed points and post-habituation PN activity; see *Supp. Materials*, sec. 8 for detailed expressions. Of note, we rescaled the *µ*_*L*_ rate to *µ*_*L*_*/*Λ^2^ in the BioPCA model (eq. 21) to preserve exactly the same learning dynamics for any Λ, just with *M* weights scaled up or down.

For comparison, we introduced a similar scale parameter Λ_IBCM_ in the IBCM model, but we generally kept it equal to 1 (its implicit value by default), since that was sufficient to achieve complete background manifold projection (Fig. S11). Similar to BioPCA, scaling of the learning rate *µ* was required for Λ_IBCM_ ≠ 1 (*Supp. Materials*, sec. 8 for details).

### Numerical simulations and model parameter values

We integrated the stochastic differential equations of the network, with the background processes simulated as described above, using an Euler scheme with time step Δ*t* = 10 ms. Below, we give rates in scaled units where this time step = 1.

By default, we performed simulations lasting 360, 000 time steps (1 hour) with *N*_S_ = 25 dimensions, *N*_K_ = 40*N*_S_ Kenyon cells, *N*_B_ = 6 background odors, *N*_I_ = 24 IBCM neurons or *N*_I_ = *N*_B_ BioPCA neurons. For *W* Hebbian learning, we used *α* = 10^−4^ and *β* = 2 *×* 10^−5^. However, for Fig. 3 and Fig. S6, we used *N*^**B**^ = 3, *N*_I_ = 6 (IBCM), *α* = 2.5 *×* 10^−4^, and *β* = 5 *×* 10^−5^.

For the IBCM weights, we used by default *µ* = 1.25 *×* 10^−3^, *τ*_Θ_ = 1600, *η* = 0.6*/N*_I_, *k*_Θ_ = 0.1, *ϵ* = 0.005, and *A* = 50 as the maximum amplitude of the nonlinearity *ϕ*. For the simple background in Fig. 3, we used *µ* = 1.5 *×* 10^−3^, *τ*_Θ_ = 200, *η* = 0.5*/N*_I_, we did not apply the *ϕ* nonlinearity or divide the learning rate by *k*_Θ_ + Θ. Also, for the simulations in higher dimensions in Fig. 5, we used a slower learning *µ* = 7.5 *×* 10^−4^ and *τ*_Θ_ = 2000. For the BioPCA model, we used by default *µ* = 10^−4^, *µ*_*L*_ = 2*µ*, and *λ*_*r*_ = 0.5 (the range of <u>Λ</u>_*kk*_ entries). For Fig. 4, we fixed Λ_PCA_ = 8 instead of the exact value making BioPCA and IBCM inhibit the background equivalently (described above). A full list of parameter definitions and values is provided in Tables S1 and S2.

The background process was initialized to a random sample from its stationary distribution. We initialized the *W* weights to zero, and the *M* weights to random i.i.d. normal samples with standard deviation 0.2 (or 0.3 for Fig. 4) for IBCM, or standard deviation 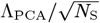 for BioPCA. For the latter model, we initialized *L* to the identity matrix (as recommended in the original paper); for IBCM, we initialized 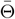 to the value of 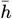 with the initial weights and background.

## Code Availability

All simulation code is available on Github: https://github.com/frbourassa/olfactory_habituation

## Author contributions

All authors designed research, FXPB performed research, and FXPB and GR wrote the paper with input from PF and MV.

## ACKNOWLEDGEMENTS

This work was supported in part by the National Science Foundation, through the Center for the Physics of Biological Function (PHY-1734030). F.X.P.B. was supported in part by an award from the Fonds de recherche du Québec – Nature et technologies. P.F. was supported by grants from the NSERC Discovery and CIHR Programs, and by a Fonds Courtois award. M.V. was supported by NIH via the Grant No. RF1NS128865-01. G.R acknowledges support from a joint research agreement between NTT Research Inc. and Princeton University.

## Supplementary Materials

### 1. Optimal models of manifold learning and predictive filtering

In this section, we detail the optimization problem we solved to delineate regimes of predictive filtering and manifold learning, as shown in Fig. 1E and Fig. S1.

#### A. Definition of the loss function

We want to minimize the loss function

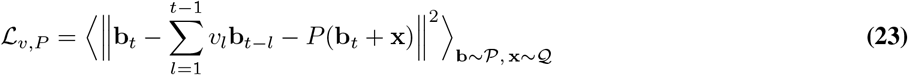

as a function of the scalar coefficients *v*_*l*_ (predictive filtering) and of the matrix *P* (manifold learning). In this section, we call 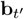 the background OSN input vector at time *t*′, and **x** the new odor (appearing at time *t*), instead of **s**_b_ and **s**_new_. The background is a linear combination of pre-defined odor vectors, **ŷ**_*ρ*_, weighted by stochastic concentrations,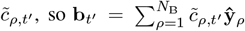. For simplicity, the concentrations are assumed i.i.d. and stationary with mean zero, variance 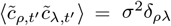, and autocorrelation function 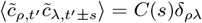, with *C*(0) = *σ*^*2*^ ; together, these concentrations statistics define the background vector distribution 𝒫. The new odor **x** comes from some distribution 𝒬 we assume has zero mean and finite covariance matrix ⟨**xx**^r^⟩.

##### A.1. Remarks on notation

In this calculation, sums and matrix products are applied over three different indices, denoting olfactory dimensions, time, and background odors, *e*.*g*., (*P* **x**)_*i*_ = *P*_*ij*_*x*_*j*_. To make notation more concise, we rewrite several sums as dot products. To clarify the indices on which these products are, we use boldface **x** on vectors in olfactory dimensions, and underlines 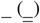 for 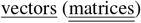 in time dimensions. We write out explicitly sums with Greek indices on background odor indices, 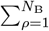.

##### A.2. Expanding the loss function terms

With the above assumptions on the background and new odor statistics, we can expand the square and write out the different terms in the loss function. First, using the statistical independence and zero mean property of **x** and **b**_*t*_, the terms to evaluate are

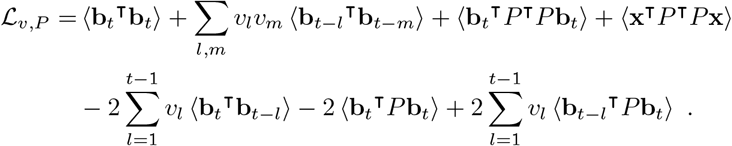

We compute these terms more explicitly by using the background statistics defined above. The loss function is thus

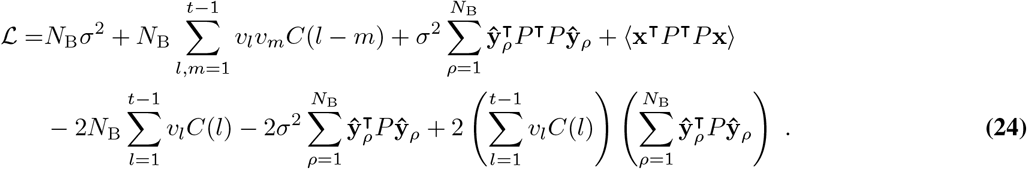

We did not need to assume that background odors were orthogonal to get this answer; the statistical independence of their concentrations 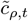 removed cross-odor terms. Most terms involve only *P* or *v*; only the last term couples the two strategies together.

#### B. Solving for the optimal *P* and *v*

##### B.1. Loss function derivatives and resulting optimum equations

We can now take the derivative of this loss function with respect to the parameters *v*_*j*_ and *P*_*ij*_. After working out the derivatives of the different terms, the result is

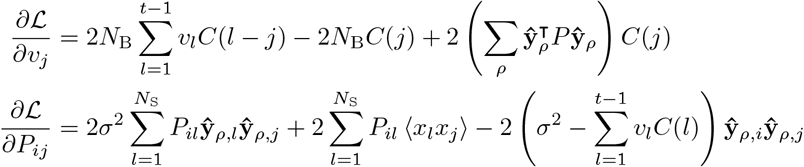

To shorten the notation of terms involving the autocorrelation function *C*(*s*), we introduce the vectors *<u>c</u>* = (*C*(1), …, *C*(*t* − 1)) and *<u>v</u>* = (*v*_1_, …, *v*_*t*−1_), and the matrix 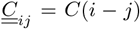. We note that 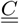 is a Toeplitz matrix (*i*.*e*., *C*_*ij*_ only depends on *i − j*), symmetric since *C*(*i − j*) = *C*(*j − i*). These properties help to express its inverse explicitly in some cases [74].

Setting the derivatives to zero to find the optimum parameters, we thus have a set of vector and matrix equations for *<u>v</u>* and *P*, respectively:

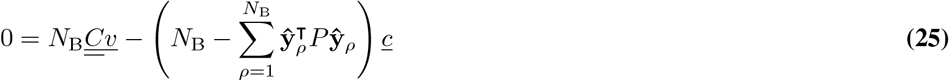

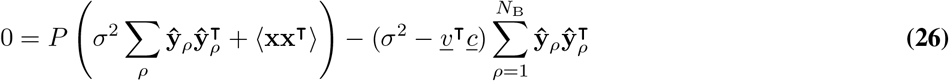

##### B.2. Solving for P in terms of <u>v</u>

The best solution path is to first solve for *P* in terms of *<u>v</u>*^*⊤*^*<u>c</u>*, then solve for *<u>u</u>*. We define the *N*_S_ *× N*_S_ symmetric matrix

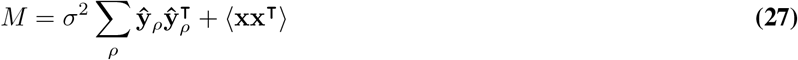

which admits a spectral decomposition *M* = *U* Σ*U*^⊤^ and a Moore-Penrose pseudo-inverse *M*^+^ = *U* Σ^+^*U*^T^, which is the actual inverse *M*^−1^ when *M* is invertible (*i*.*e*., no zero eigenvalue in Σ). Equation 26 thus takes the form 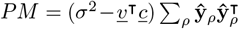,which can be inverted for *P*_*M*_ = *PU* ΣΣ^+^*U*^T^, the component of *P* in the subspace spanned by *M* ‘s eigenvectors of non-zero eigenvalues. The *P* component in the null space of *M*, if any, is not constrained by this optimization problem, so we set it to zero, and take *P* = *P*_*M*_. Hence,

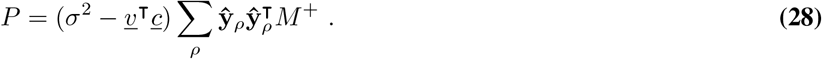

##### B.3. Solving for <u>v</u>

We can now insert the implicit solution for *P* in equation 25 for *<u>v</u>*. We first evaluate the term

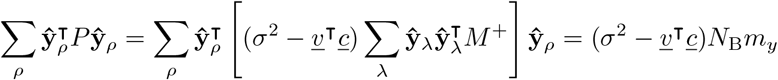

where we have defined

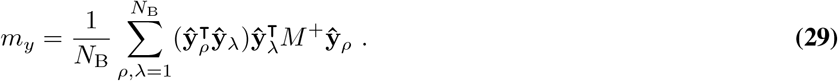

that background odor directions **ŷ**_*ρ*_ were orthogonal, but if that were the case, *m*_*y*_ would simplify to 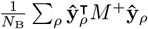. Inserting in eq. 25, dividing by *N*_B_, and isolating *<u>v</u>* by assuming that the autocorrelation matrix 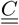 is invertible, we have

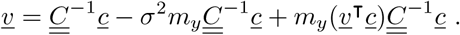

This is still an implicit expression because *<u>v</u>*^*⊤*^*<u>c</u>* appears on the right; taking the dot product of this expression with *<u>c</u>*, we can isolate *<u>v</u>*^*⊤*^*<u>c</u>*, then reinsert in the implicit equation for *<u>v</u>* to arrive at an explicit solution (*i*.*e*., in terms of 𝒫, 𝒬 parameters),

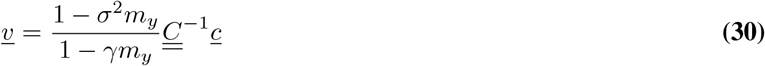

where we have defined

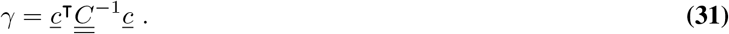

##### B.4. Replacing <u>v</u> in the P solution

Having solved for *<u>v</u>*, we can put it back in 28 to obtain an explicit solution for *P*,

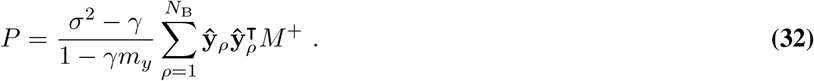

#### C. Evaluating the loss function at the optimum

For simplicity, we first rewrite the loss function in eq. 24 using the underlined vector notation for 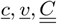, giving

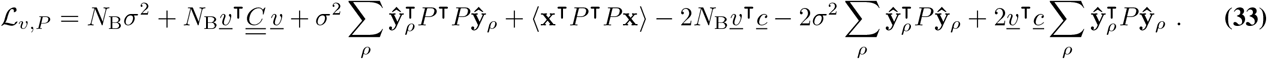

We need to evaluate multiple terms to insert the solutions for *<u>v</u>* and *P* in ℒ. We find several simplifications by using the fact that *M* and thus *M*^+^ and *P* are symmetric (eqs. 27 and 32), commuting scalars resulting from intermediate dot products, and renaming indices when appropriate. We find the following terms,

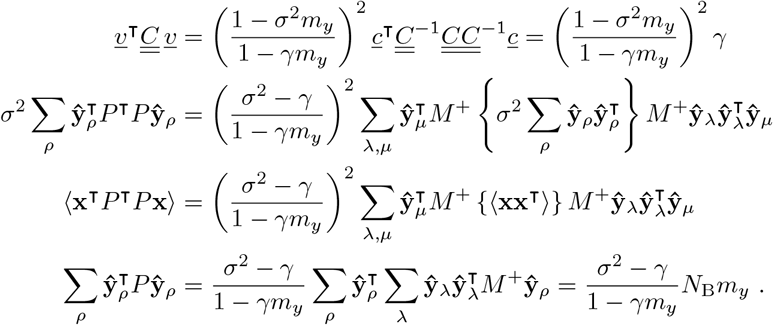

The second and third terms can be combined by noticing they have the same form with sums over odor indices *λ, µ*, and combining the bracketed terms to find 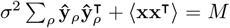. Then, using the definition of the pseudo-inverse, we have *M*^+^*MM*^+^ = *M*^+^, resulting in

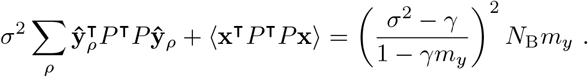

Combining these expressions in ℒ, we can cancel out a few terms with further algebra and factorize common expressions, finding a surprisingly simple form,

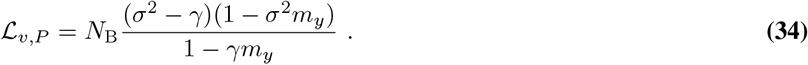

We notice that the loss seems to scale proportionally with the background subspace dimensions, *N*_B_. Terms *σ*^2^, *γ* do not depend on *N*_B_ but only on the autocorrelation and variance of odor concentrations. The only term that could depend on *N*_B_ is 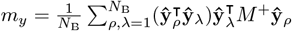, but we generally expect *N*_B_*m*_*y*_ *∼ N*_B_. This is especially clear if we assume orthogonality of the **ŷ**, such that 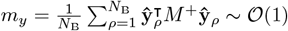.

Hence, there is no obvious tradeoff between the two strategies – predictive filtering and manifold learning – as a function of the background dimension. This makes sense *a posteriori*. Predictive filtering tries to anticipate *N*_B_ independent, identically distributed odors, hence the squared errors committed on each background component add up in variance. Meanwhile, the error in manifold learning increase with *N*_B_ because a fraction ∼ *N*_B_ of the new odor, on average, will lie in the background subspace.

However, there is a tradeoff between the strategies as a function of the autocorrelation time, encoded in the parameter *γ*, and the dimensionality of the olfactory space, which enters *m*_*y*_ through the new odor statistics ⟨**xx**^T^⟩ in *M*. We expect *γ* to increase with the autocorrelation time scale, and *m*_*y*_ to increase with the olfactory space dimension *N*_S_. Hence, as the autocorrelation time increases, *γ* diminishes the relative efficacy of manifold learning by reducing the denominator 1 − *γm*_*y*_, while increasing the importance of predictive filtering by reducing the numerator factor *σ*^2^ − *γ*. This tradeoff will be clearer in the special case studied in section F.

#### D. Summary of the general optimal solution

We recapitulate the optimization results here. The optimal *<u>v</u>* and *P* are

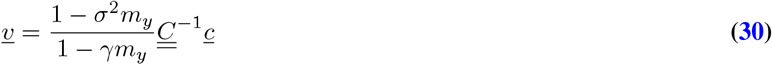

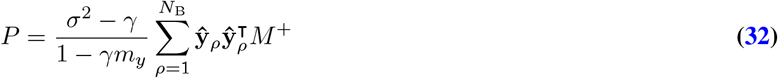

and they give a minimum loss of

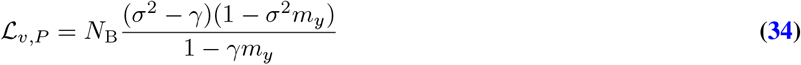

where *N*_B_ is the number of i.i.d. background odors, *σ*^2^ is the variance of each odor concentration 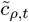, and where we have defined background parameters

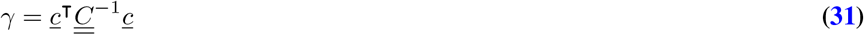

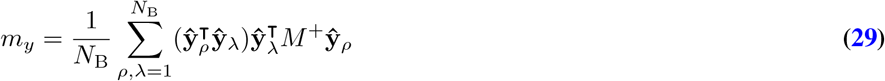

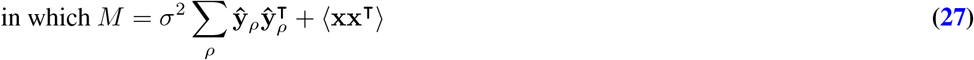

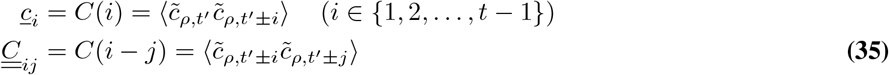

#### E. Limiting cases

*P* = 0 **and** *<u>v</u>* = 0.

##### E.1. Predictive filtering only

*P* = 0. When *P* = 0, we can directly solve eq. 25 for *<u>v</u>*, finding

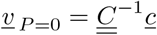

which yields a loss of

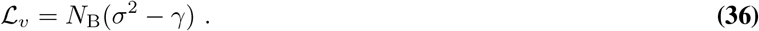

As long as *γ < σ*^2^ – which should be the case if the autocorrelation function decays with time – we have ℒ_*v,P*_ *<* ℒ_*v*_ since 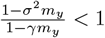 in that case.

##### E.2. Manifold learning only

*<u>v</u>* = 0. When *<u>v</u>* = 0, we can directly solve eq. 26 in terms of *M*^+^, resulting in

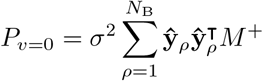

which yields a loss of

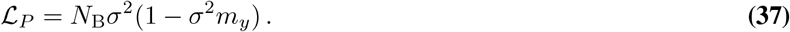

As long as 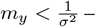 which should be the case since *M*^+^ ∼ 1*/σ*^2^ and *m*_*y*_ is some projection of it on the background subspace – then ℒ_*v,P*_ *<* ℒ_*P*_, since 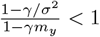 in that case.

#### F. Special case: exponential kernel, 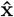 uniform on hypersphere

To make these results more concrete, we now consider a simple case of background statistics where expressions such as *γ, m*_*y*_, etc. can be computed analytically in terms of interpretable parameters. We consider an exponential autocorrelation function and new odors uniformly sampled on a hypersphere. This is the case plotted in Figs. 1E and S1.

##### F.1. Exponential autocorrelation kernel, to evaluate γ

We suppose that each odor concentration 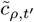 is independent of other odors and forms a Gaussian process with exponential autocorrelation kernel 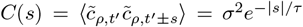 with auto-correlation time *τ* (*i*.*e*., the Ornstein-Uhlenbeck process). A small *τ* corresponds to fast fluctuations compared to the time scale of learning. In this case, the symmetric Toeplitz matrix 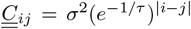 is a Kac-Murdock-Szegö matrix (form *A*_*i−j*_ = *r*^|*ij*|^, *r* ≠ 1), which has an explicit inverse, provided in [74, sec. 1.3]. This inverse is the tridiagonal matrix

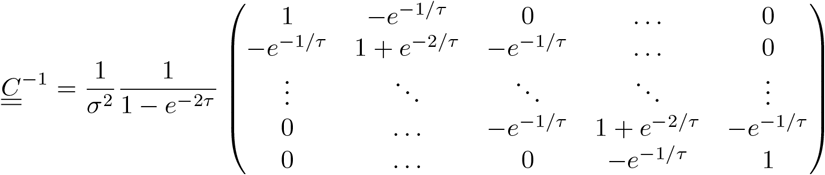

It is not hard to check that this is indeed the inverse of 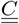. This allows us to evaluate

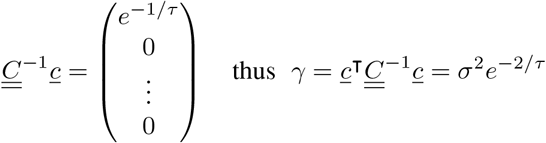

##### F.2. New odors 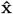 uniform, and ŷ_ρ_ orthogonal, to evaluate m_y_

Moreover, we suppose that new odors take the form 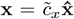, where 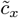 has variance 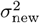 possibly different from the background odors, and where 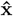 is uniformly sampled on the *N*_S_-dimensional unit hypersphere. In that case, by symmetry, the new odor covariance matrix is 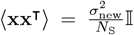, with 𝕀 the *N*_S_ *× N*_S_ identity matrix. Additionally, we assume that background odors are orthogonal to each other: 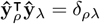.

These choices allow us to compute *M*^+^ explicitly. We first note that *M* = ⟨**xx**^⊤^⟩ + *σ*^2^ ∑_*ρ*_ **ŷŷ**_*ρ*_ has full rank, and thus *M*^+^ = *M*^−1^. We define the rescaled olfactory dimension

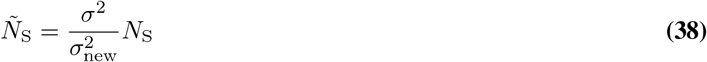

to simplify expressions (equal to *N*_S_ if the background and new odors have the same concentration variance, 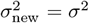). We can compute that inverse by repeatedly applying the Sherman-Morrison formula to the sequence of matrices 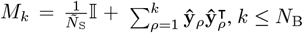. Proceeding by induction, we eventually find the inverse of the full matrix 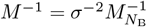,

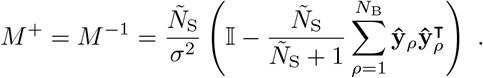

We can thus evaluate the matrix product appearing in the optimal *P* solution,

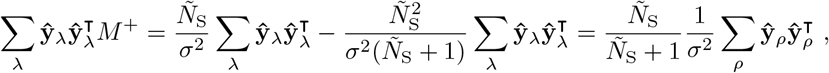

using 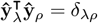, which also allows us to compute

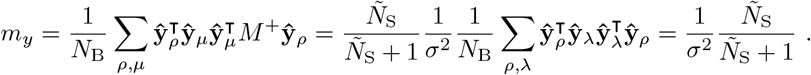

##### F.3. Optimal <u>v</u>, P and loss function in this background choice

Inserting the above expressions for *γ, M*^+^, *m*_*y*_, etc. into the general optimal solution, we find

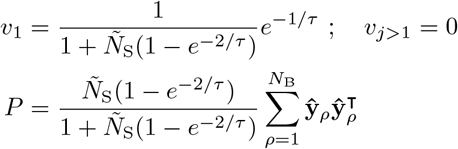

and a minimum loss of

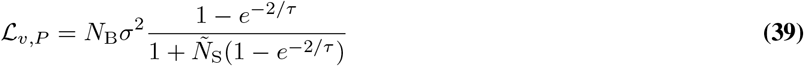

Here, we see clearly that there is no tradeoff as a function of *N*_B_: both strategies have an error that increases proportionally to *N*_B_. At least, we clearly see the transition from predictive filtering to manifold learning as 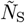 decreases or *τ* decreases. For small correlation times, 1 *e*^2*/τ*^ *≈* 1, so the main reduction of the loss comes from the 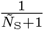 factor (also the case if *Ñ*_S_ is large). Meanwhile, for large correlation *τ*, the reduction comes from the numerator 1 − *e*^−2*/τ*^ ≈ 0, while the *Ñ*_S_ term in the denominator is rendered ineffective – the same is true if *Ñ*_S_ is small.

##### F.4. Special cases P = 0 and <u>v</u> = 0 in this background choice

For further comparison, the optimal solutions and loss function in the pure predictive filtering case (*P* = 0) with our specific background choice are:

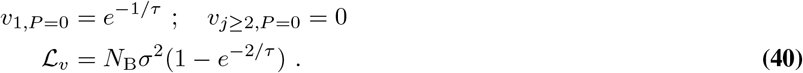

In the pure manifold learning case (*<u>v</u>* = 0), these are rather

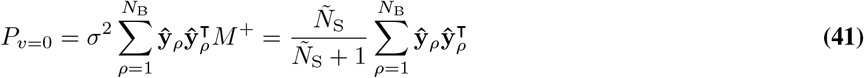

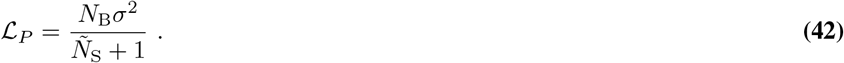

##### F.5. Plots of the loss function versus Ñ^S^ and τ for the different strategies

We notice that the loss is proportional to *N*_B_*σ*^2^ in all strategies for our special background choice, hence we can illustrate the relative efficacy of predictive filtering and manifold learning by plotting ℒ*/*(*N*_B_*σ*^2^) as a function of *Ñ*_S_ and *τ*, the two background parameters for which there is actually a transition between the two strategies. Fig. S1A-B shows the single-strategy losses ℒ_*v*_ (eq. 40) and ℒ_*P*_ (eq. 42) compared to the loss for both strategies applied simultaneously, ℒ_*v,P*_ (eq. 39).

We see that for even a small olfactory space dimension, *Ñ*_S_ = 50 as in the fruit fly, manifold learning performs much better than predictive filtering even for moderately long correlation time scales, as discussed in the main text. When ℒ_*v,P*_ is close to either ℒ_*P*_ or ℒ_*v*_, it means that the corresponding strategy contributes most of the loss reduction. We can thus draw a “phase” diagram of where habituation is dominated by manifold learning (large *Ñ*_S_, small *τ*) or by predictive filtering (small *Ñ*_S_, large *τ*). The (smooth) transition occurs when ℒ_*P*_ = ℒ_*v*_, which happens at 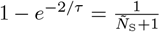, which is at *τ* ≈ 2(*Ñ*_S_ + 1) for large *Ñ*_S_. This is shown in Fig. 1E. There is a large region (in red) where manifold learning is the most effective strategy.

### 2. Simulating background odor fluctuations

#### A. Turbulent statistics

We use the statistics of whiff durations 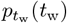, blank durations 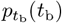, and whiff concentrations *p*_*c*_(*c*) derived in [9], with the exponents and statistics for an atmospheric boundary layer. We simulate each background odor concentration as a telegraph-like process, as illustrated in Fig. 1, by drawing a next blank duration at the end of a whiff, a next whiff duration at the end of a blank, and a concentration for each whiff from *p*_*c*_. As the simulation advances per time steps Δ *t* = 10 ms, we keep track of how much time is left in the current whiff or blank as well as of the current concentration, and update when that time runs out. This method neglects intra-whiff concentration fluctuations, as well as correlations between successive whiff and blank durations, but captures the main challenges of varying whiff concentrations and power-law (long-tailed) distributions of durations. We now describe these distributions and how we numerically sample from them.

For the durations of whiffs and blanks, the distribution is a power law with exponent −3*/*2 and a lower cutoff at *τ*_b_, *τ*_w_. For numerical stability, we prevent abnormally large durations by imposing also an upper cutoff at *T*_max,b_, *T*_max,w_, and we use sharp cutoffs, corresponding to a probability density function

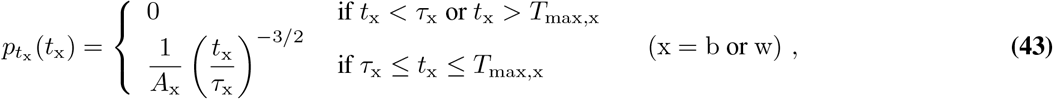

where *A*_x_ = 2*τ*_x_ (1−(*T*_max,x_*/τ*_x_)^−1*/*2^) is a normalization constant. From these distributions, the average duration of a whiff or blank is the geometric average of the limits, since

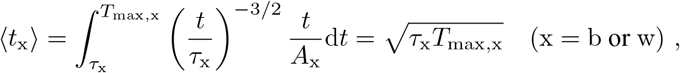

so the probability *χ* to be in a whiff is

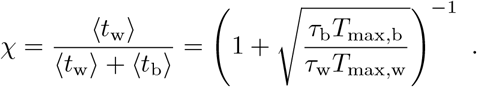

The probability distribution of whiff concentrations *c* is therefore 0 with probability 1 − *χ* (illustrated by the point at *c* = 0 in Fig. 1C, left) or, with probability *χ*, the conditional distribution *p*_*c*_ given there is a whiff. Hence, *p*_*c*_(*c*) = (1−*χ*)*δ*(*c*) + *χp*_*c*_(*c*|whiff). The conditional distribution has a tail 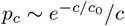, where *c*_0_ is a typical concentration scale, and a probability plateau near *c* = 0. As illustrated in Fig. 1C, left, we use a sharp transition at *α*_*c*_*c*_0_ for some *α*_*c*_ *<* 1, with a uniform probability on the range below, (0, *α*_*c*_*c*_0_], corresponding to a probability density function

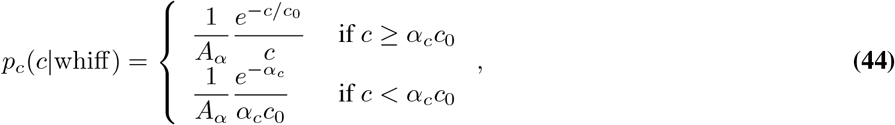

where 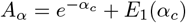 is a normalization constant and *E*_1_ is the exponential integral,

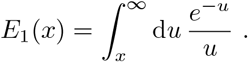

Of note, the average whiff concentration then has the analytical expression

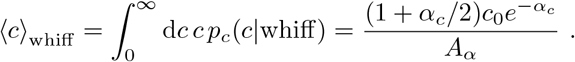

To sample from these distributions during a simulation, we use the inverse transform method: given a random uniform(0, 1) sample *r*, we generate a sample of a random variable *X* following the cumulative distribution function (cdf) *F*_*X*_ as 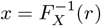. The cdf for the whiff or blank durations is

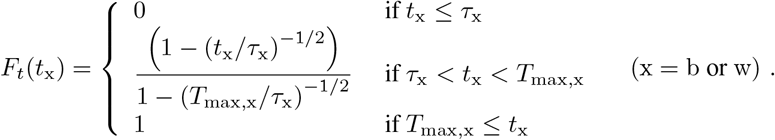

Taking the inverse, we generate *t*_x_ from uniform samples *r* as

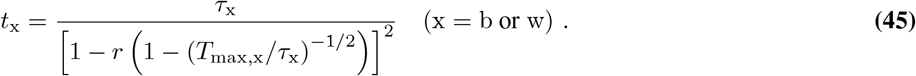

As a check, notice that *t*_x_ = *τ*_x_, the lower cutoff, when *r* = 0, and *t*_x_ = *T*_max,x_, the upper cutoff, when *r* = 1. The cdf for the conditional whiff concentrations is

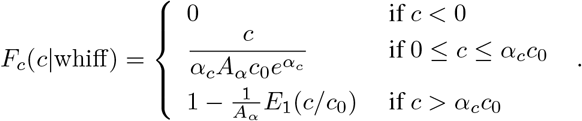

Hence, given a random uniform sample *r*, we generate a sample *c* as

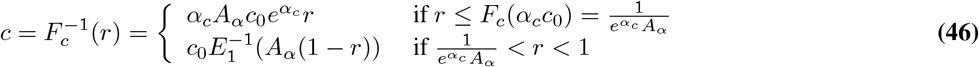

where 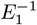 is the inverse exponential integral. This inverse function does not have an analytical closed form, so we evaluate 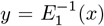 at a given *x* numerically by solving the equation *E*_1_(*y*) − *x* = 0 for *y*, using Brent’s method [75] with suitable bounds on the solution. For numerical accuracy, for *x <* 1, we solve in log scale, log(*E*_1_(*y*)) − log(*x*) = 0, to expand the range of *E*_1_(*y*) values. For larger *x, y* becomes very small (*e*.*g*., *E*_1_(2) = 0.0489), so we solve for *z* = log(*y*). For *x >* 30, *y* is small enough (*y* ∼ 10^−13^) to use the approximation *E*_1_(*y*) = −*γ* − log(*y*) − 𝒪 (*y*) [76, 6.6.1], where *γ* = 0.577 … is the Euler-Mascheroni constant, so the equation is inverted directly: *y* = *e*^− *γ*−*x*^.

Hence, overall, for each update when a whiff starts, we use two random uniform(0, 1) samples, one for *t*_w_ (using Eq. (45)), one for *c* (using Eq. (46)); only one sample is needed when a blank starts, for *t*_w_ (Eq. (45)). In our simulations, we use the following typical parameter values, the same for all background odors: *τ*_b_ = *τ*_w_ = 10 ms = 1 time step for the lower whiff or blank duration cutoff, *T*_max,w_ = 5000 ms and *T*_max,b_ = 8000 ms for the maximum whiff and blank durations respectively, *c*_0_ = 0.6 for the (arbitrary) concentration scale, and *α*_*c*_ = 0.5 for the lower whiff concentration cutoff.

Moreover, to sample a concentration from the stationary distribution, we draw a first uniform(0, 1) sample *r*_1_ to determine whether there is a whiff, with probability *χ* (whiff if *r ≤ χ*), or a blank (*c* = 0). Then, if in a whiff, draw a second uniform sample *r*_2_ to generate a concentration *c* using Eq. (46).

#### B. Univariate Ornstein-Uhlenbeck process

To simulate a univariate Ornstein-Uhlenbeck (O-U) process *ν*(*t*) numerically, we use an exact update rule for finite time steps Δ*t*, derived from the analytical solution of the O-U process. Taking the last time step as a new deterministic initial condition [77, eq. 2.47],

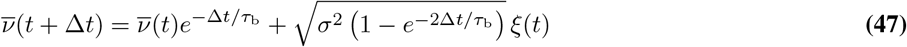

where *ξ*(*t*) is white noise. The coefficients of 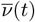 and *ξ*(*t*) can be computed in advance. This rule ensures a steady-state distribution of 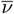 with the desired variance *σ*^2^ even when the simulation time step is on the order of *τ*_b_.

#### C. Multivariate Ornstein-Uhlenbeck process

The multivariate Langevin equation for the Ornstein-Uhlenbeck process with zero stationary mean, 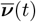, is covered in [78, sec. 4.5.6], and can be simulated exactly using the same trick as in the univariate case, Eq. (47). In practice, we used independent and identically distributed background odors in this paper, so the matrices *A* and *B*, were diagonal, and the general simulation method effectively reduced to simulating *N*_B_ zero-mean univariate processes in parallel (using Eq. (47)), then adding the desired mean vector 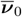 to it.

#### D. Weakly non-Gaussian fluctuating background

We simulate a multivariate Ornstein-Uhlenbeck process, **g** as in section 2C, with zero mean and identically distributed variables with stationary variance 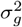. Then, we take 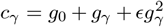, where ϵ should be chosen small and *g*_0_ is the desired zeroths-order mean concentration. Then, the concentrations have the following moments:

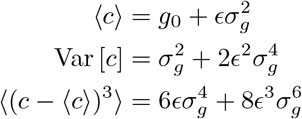

These results are straightforward to obtain by expanding *c*^2^ and *c*^3^ and using higher moments of the Gaussian distribution as required, *i*.*e*., 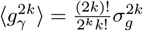. The important outcome is that the third moment is of order ϵ.

#### E. Log-normal background fluctuations

As above, we simulate identically distributed O-U variables with variance 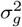, add a mean *g*_0_ to them, then use them as the log_10_ of the concentrations, transforming them according to 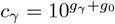. Then, from the log-normal distribution properties [79], the concentrations themselves have a log-normal distribution with moments

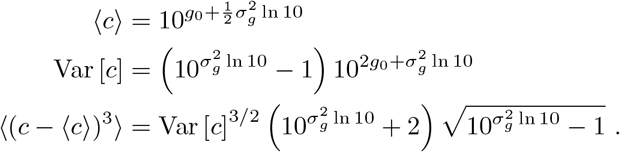

We test habituation to this background with IBCM and BioPCA networks in Fig. S6.

#### F. Numerical stability

Numerical integration of the *W* equations displays instabilities when 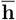 reaches large magnitudes, *e*.*g*., when increasing the scale parameter Λ (Fig. S11 and section 8). To ensure these are numerical errors rather than a true dynamical instability of the fixed points, we perform a nonlinear numerical stability analysis of the Euler integrator. This integrator, applied to the *W* matrix equation, is effectively a discrete mapping,

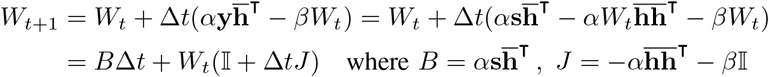

We consider the worst-case scenario, when 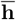 reaches its maximal magnitude encountered in a simulation, 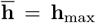 and 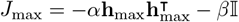, and iterate the map in this case, which gives

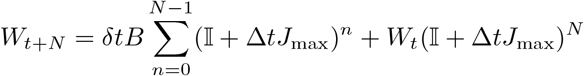

Consequently, the stability depends on the eigenvalues of *A* = 𝕀 + Δ*tJ*_max_, but not on Δ*tB* (because the latter is not raised to some power). Indeed, first note that the matrix *A* is symmetric and diagonalizable as *A* = *UDU*^*†*^ with *D* = diag(*λ*_*i*_). Then, if *A* has at least one eigenvalue with magnitude |*λ*_*j*_| *>* 1, then 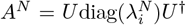 will have a diverging component as the mapping is iterated (as *N* increases). To find the threshold where this happens, we can read out the eigenvalues from the expression of 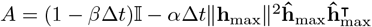,

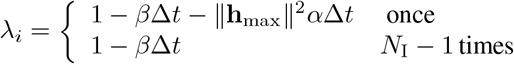

To see this, consider the rotation matrix *R* that aligns **h**_max_ with one of the unit vectors, *e*.*g*., (1, 0, … 0): *RAR*^r^ is then diagonal with *λ*_1_ = 1 *βt* ‖**h**_max_‖^2^*α*Δ*t* in one row, *λ*_2_ = 1 − *β*Δ*t* on the others. For large LN activity, *λ*_1_ is first to reach a magnitude *>* 1, by becoming negative. Hence, we can predict that numerical instabilities arise in the Euler integrator when

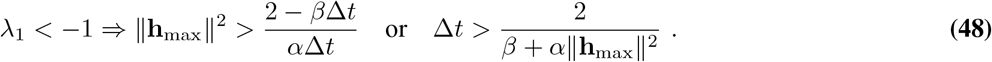

Given a simulation of the *M* weights and 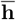 (which are not destabilized), we can thus anticipate whether the *W* integration should be unstable by extracting ‖*h*_max_‖ from the simulation. The vertical lines in Fig. S11 indicate the smallest Λ value for which this threshold is reached, in each model, in the turbulent background considered. They coincide well with the observed drops in model performance, confirming these are due to Euler integrator instabilities. A linear stability analysis of the *W* equations (computing the Jacobian of the ODE near the fixed point, etc.) confirms that the *W* is linearly stable for any Λ. Hence, the observed divergences are numerical limitations rather than true model performance drops and would be remedied by decreasing the time step, Δ*t*.

### 3. Average subtraction model

In this short section, we examine the average subtraction model from [36], and explain how it is insufficient against fluctuating backgrounds. In our notation, it corresponds to a vector **w** of inhibitory weights learned as

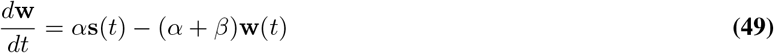

and a PN response **y**(*t*) = **s**(*t*) − **w** (the LN activity is fixed to 1). If this network is exposed to a constant background odor **s**_b,0_, the inhibition vector **w** converges to 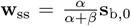, so the background is then perfectly subtracted. However, this strategy fails if the background vector **s**_b_(*t*) fluctuates randomly over time. In this case, equation 49 amounts to computing the average background over a time window of duration 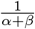 as seen from the formal solution of the stochastic equation,

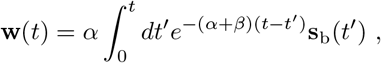

assuming **s**_b_(*t*) started at a given initial value **s**_b,0_. From the formal solution, assuming the background is a stationary process, the steady-state average value of **w** is therefore proportional to the average background,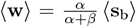. If the background process has an autocorrelation time scale much faster than the learning rate, 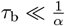, with its elements approximately obeying 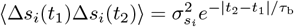, then to leading order, the inhibitory weights have a small variance

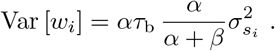

Hence, the inhibitory weights do not fluctuate much around the constant average background, ⟨ *s*_b_⟩, since the factor *ατ*_b ≪_1. Therefore, this model computes the average background, scaled by 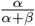, and subtracts this fixed quantity from **s**_b_(*t*) to obtain the projection neuron activity **y**(*t*) = **s**_b_(*t*) − **w**. Consequently, the variance of the PN activity, **y**(*t*), is not reduced compared to the variance of the background:

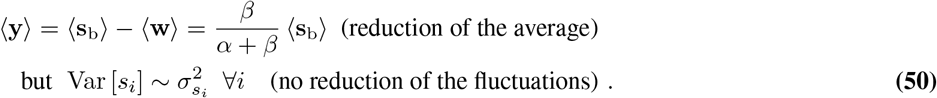

These large remaining fluctuations in PN activity mix with new odors appearing in the landscape and thus hinder their recognition; this effect explains why the average subtraction model does not provide a significant reduction in PN response to the background, or improvement in new odor recognition compared to the absence of habituation in Figures 2, 5, S2, and others.

### 4. Analytical solution of the IBCM model’s average fixed points

To understand what projections are learned by IBCM neurons, we derive analytical expressions for the synaptic weights **m** of an IBCM neuron at stationary state. We first establish approximate fixed point equations for these weights averaged over fast fluctuations of the background process **s**_b_(*t*), assuming perfect separation of time scales between **s**, Θ, and **m**. Then, we obtain exact solutions to these approximate equations.

#### A. Establishing the IBCM fixed point equations

Let’s first recall the stochastic differential equations describing the synaptic weights learning of an IBCM neuron in a network with feedforward lateral coupling (Methods). For tractability, we assume the activation function ϕ is the identity function (instead of a nonlinearity like tanh), such that 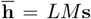. We also set the decay term −*εµ***m**_*i*_ to zero. Then, for each neuron *i*,

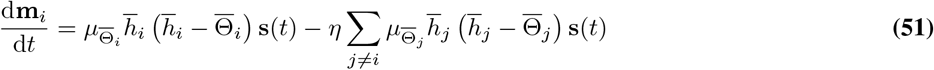

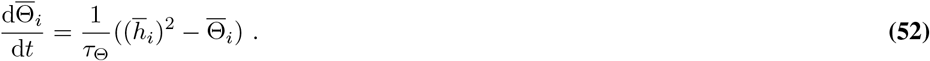

The reduced activity of interneuron is 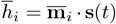, where the reduced weights 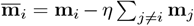. Hence, 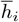 varies in time on two separate scales: rapidly with sensory inputs **s**(*t*) fluctuating on time scale *τ*_b_, and gradually as the synaptic weights 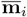 are learned.

To make analytical progress, we focus on averages over fast background fluctuations, denoted by brackets ⟨*·* ⟩, while the slow dynamical variables 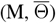 remain unchanged. Moreover, we make a quasi-static approximation on the thresholds: we assume *τ*_Θ_ is slow enough to average over background fluctuations, yet fast enough that 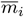 remains unchanged over the averaging time window (*i*.*e*., we neglect correlations between 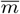 and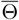). Hence, as in the original IBCM models [43], we assume a perfect separation between the background, threshold, and synaptic weight time scales: 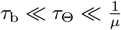.

Thus, averaging equation 52, and setting 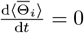, we find

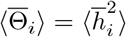

and we replace 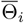 in the **m** equation 51 with this average. Averaging that equation as well, and setting 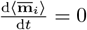, we find the IBCM fixed point equations:

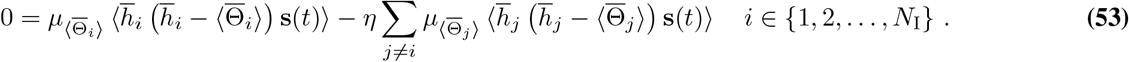

Defining terms 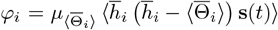 and combining them in a *N*_I_-dimensional vector *φ*, this system of equations can be written in matrix form

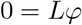

where *L* is the *N*_I_ *× N*_I_ matrix of feedforward inhibitory coupling between interneurons, with 1 on the diagonal and −*η* everywhere else. This *L* is a circulant matrix (each row is a cyclic permutation of the previous row by one element to the right); hence, except in the pathological cases *η* = −1 or 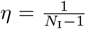 (for which some of its eigenvalues are zero), *L* is invertible and the unique solution is φ = L^−1^0 = 0. Therefore, in general, the fixed points of the IBCM network are found by setting each φ_*i*_ term to zero individually

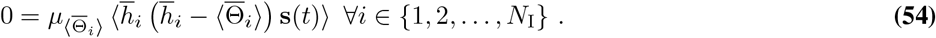

Hence, in terms of the reduced synaptic weights 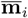 and activities 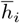, the fixed point equations for the network of IBCM neurons decouple to take the same form as that of a single IBCM neuron.

Before proceeding to solve this set of equations, we note that the actual synaptic weights **m**_*i*_ can be found from the 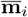 solutions by inverting the matrix equation

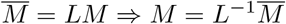

where 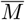 contains the reduced 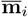 in its rows. From the eigenvectors and eigenvalues of the circulant matrix *L*, the inverse matrix elements are

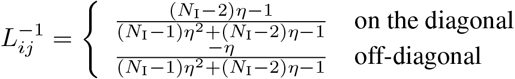

so the synaptic weights **m**_*i*_ can be recovered from the reduced ones, if necessary, as

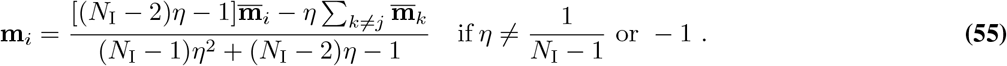

#### B. IBCM fixed point equations for i.i.d. background concentrations

We will express the solutions to the fixed point equations in terms of the alignments, or dot products, of the IBCM synaptic weights with the background odors,

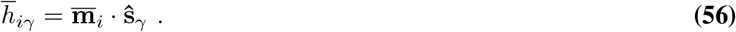

and we imply these are averaged over fast background fluctuations (*i*.*e*., we really look at 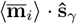), The odor vectors **ŝ**_*γ*_ do not have to be orthogonal, but we assume they form a linearly independent set. Specifying these dot products is sufficient to obtain the complete solution, since the IBCM dynamics only update weights in the background subspace of **s**(*t*) (eq. 51). The other directions are reduced to zero by the slow decay term, −*δµ***m**_*i*_, that we have added to the full dynamics (Methods). For the rest of this section, we work with reduced variables and drop the overlines on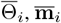, and 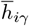 to simplify notation. We also drop the IBCM neuron index *i* since the fixed point equations eq. 54 are identical and solved independently for each neuron.

Since we have decoupled the Θ and **m** fluctuations in Eq. (53), we can divide by the learning rate and write

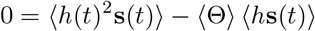

We replace ⟨ Θ ⟩ = ⟨*h*^2^⟩, using our quasi-static approximation. We now assume that the odor concentrations *h*_*γ*_(*t*) are independent, identically distributed stationary processes. We write 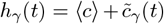 where 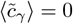. The concentrations have mean ⟨*c*⟩ variance 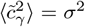 and third moment 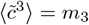. We also let 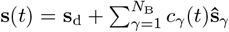 and 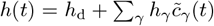 where we have defined **s**_d_ = ⟨*c*⟩ ∑_*γ*_ **ŝ**_*γ*_ and *h*_d_ = **m** *·* **s**_d_. We proceed to compute the averages appearing in the fixed point equation:

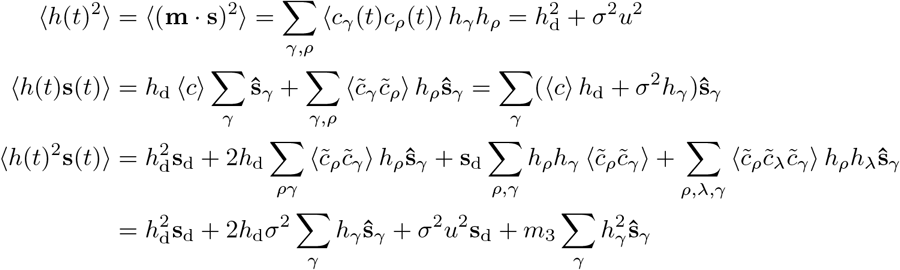

where we have defined

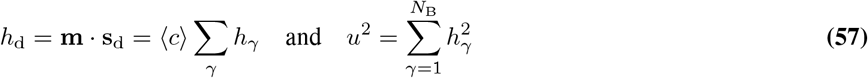

Combining and expanding **s**_d_ = ⟨*c*⟩ ∑_*γ*_ **ŝ**_*γ*_, we have

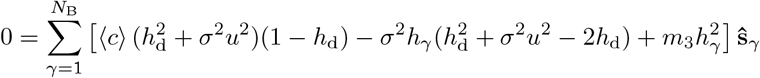

Since the **ŝ**_*γ*_ odors are linearly independent, we have, for each IBCM neuron, a set of *N*_B_ equations specifying the neuron’s alignment with each odor, *h*_*γ*_. These are the fixed point equations to solve for the *h*_*γ*_s:

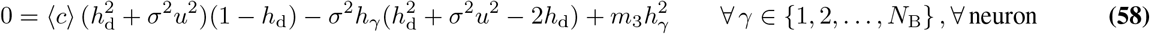

These equations are cubic polynomials in the *h*_*γ*_s, since *h*_d_ = ⟨*c*⟩ ∑_*γ*_ *h*_*γ*_ and 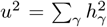. There is always a fixed point at *h*_*γ*_ = 0 ∀ *γ*, but it is unstable; trajectories initialized near the origin move away from it during habituation.

#### C. Solution for a zero third moment background processes

We first examine solutions when *m*_3_ = 0, *e*.*g*., Gaussian backgrounds, since the solutions simplify greatly in that case. The equations then have two terms,

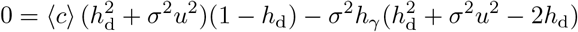

which can be made individually zero by setting *h*_d_ = 1 and *σ*^2^*u*^2^ = 1. Thus, these fixed points are defined by two constraints,

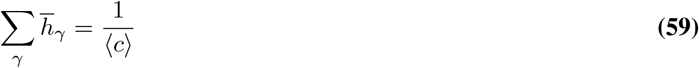

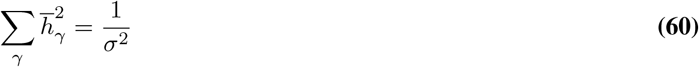

which correspond, geometrically, to the intersection of a hyperplane where coordinates sum to one, and a hypersphere of radius 1*/σ*^2^, respectively. All **m** weights on the resulting (*N*_B_ − 2)-dimensional surface in the background subspace are non-isolated fixed points when *N*_B_ *>* 2; Fig. S5A shows a three-dimensional background example, with each neuron converging to a point on the ring defined by these equations. For a two-dimensional background, there can be zero, one, or two isolated fixed points, while for a one-dimensional background, these equations do not apply.

The fixed point equation for a Gaussian background admits another set of solutions, where the two terms are not individually zero. Isolating *h*_*γ*_, we find it must have the same value for every odor, *h*_*γ*_ = *h*_0_. Then, 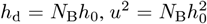, and we can solve to find

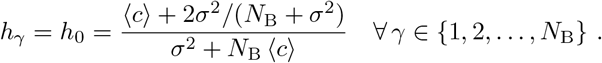

For a one-odor background, this solution would be the stable fixed point; for higher dimensions, we find in practice that it is unstable (see also D.1).

#### D. Solution for a general background with non-zero third moment

We return to solving the full fixed point equations, eq. 58, when *m*_3_ ≠ 0. Now, because of the *m*_3_*h*_*γ*_ term, setting *h*_d_ = 1 and *u*^2^ = 1*/σ*^2^ does not satisfy the equations, so we must solve for individual *h*_*γ*_s. This leads to a finite number of isolated fixed points: as shown in Fig. S5B, the third moment of the background breaks the degeneracy seen in the Gaussian case.

First, we notice that the dot products 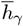 of an IBCM neuron can only take at most two different values at steady-state. To show this, we take the difference between the equation for *h*_*γ*_ and some other *h*_*α*_, which gives

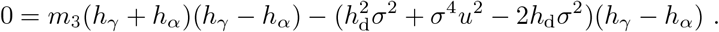

Either *h*_*γ*_ = *h*_*α*_, or if they have different values, then they are related by the constraint

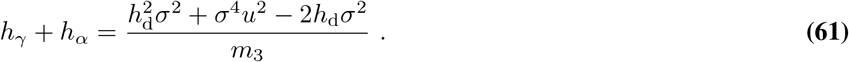

Note that constraint 61 is removed for Gaussian backgrounds *m*_3_ = 0, explaining the non-isolated fixed points in that special case. For any pair *γ, α*, the r.h.s. is the same; if we imagine a third dot product *h*_*β*_, then *h*_*γ*_ + *h*_*α*_ = *h*_*γ*_ + *h*_*β*_ ⇒ *h*_*α*_ = *h*_*β*_, implying that there cannot be a third distinct value. Therefore, either all *h*_*γ*_s are equal, or they each take one of two possible values.

##### D.1. All h_γ_s are equal

First, consider the case where all *h*_*γ*_ = *y*, a unique dot product value. Then *u*^2^ = *N*_B_*y*^2^ and *h*_d_ = *N*_B_ ⟨*c*⟩ *y*. Inserting in Eq. (58), we can factor out *y*^2^, giving

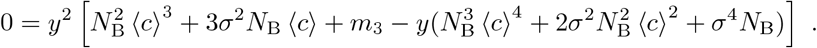

So, either *y* = 0, which is an unstable fixed point, or

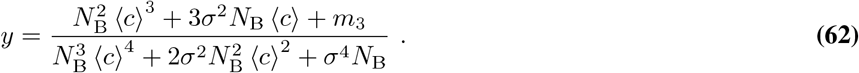

We conjecture that this fixed point is always unstable, based on the linear stability analysis of section 4F below. Fig. S3 shows that in the background process examples we considered, it is a saddle point, approached before the IBCM neuron becomes selective for a background odor.

##### D.2. Two different h_γ_ values (general case

Second, consider the case where **m** has a dot product equal to *y*_1_ with *k*_1_ odors, and equal to *y*_2_ with the remaining *k*_2_ = *N*_B_ − *k*_1_ odors. We let *y*_1_ *> y*_2_ by convention. The values *y*_1_ and *y*_2_ will depend on the repartition *k*_1_, *k*_2_, but there will be a unique pair *y*_1_, *y*_2_ for each choice of *k*_1_, *k*_2_. Moreover, in this case,

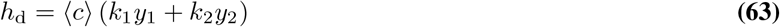

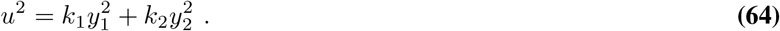

Then, the set of *N*_B_ equations 58 really reduces to two equations, one for all the *h*_*γ*_ = *y*_1_ and the other for *y*_2_, which are symmetric under 1 *↔* 2.

We start from Eq. (61) – the difference between the two values of *h*_*γ*_. We rewrite the equation in terms of *y*_1_ and *y*_2_ as

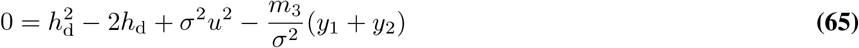

by letting one of the *h*_*γ*_s be equal to *y*_1_ and the other, to *y*_2_. We use this equation to replace, where appropriate, the following term in the fixed point equation 58,

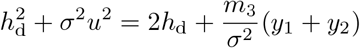

for, say, *h*_*γ*_ = *y*_1_ (using *y*_2_ would not make a difference), resulting in

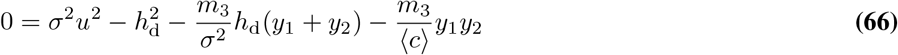

Together, equations 65 and 66 form our system of equations to solve for *y*_1_ and *y*_2_. Obtaining the latter was the crucial simplification to make, because it eliminates terms linear or cubic in *y*_*i*_, allowing us to easily isolate *y*_2_ in terms of *y*_1_ (or vice-versa)^1^. Indeed, writing *h*_d_ and *u*^2^ in terms of the *y*_*i*_ (equations 63-64), we find it takes the simple, symmetric form

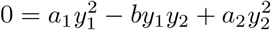

where

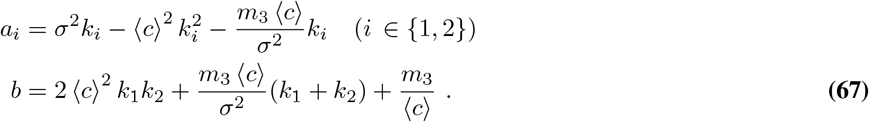

This equation makes clear the symmetry of solutions under exchange of labels 1 *↔* 2, confirming that we can keep only the roots where *y*_1_ *> y*_2_, knowing that other roots would be found by exchanging *k*_1_ and *k*_2_; in other words, the roots 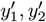 for 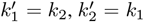, are the solutions for *k*_1_, *k*_2_ with *y*_2_ *> y*_1_ (this can be checked explicitly with the solution below). For now, we write *y*_2_ in terms of *y*_1_,

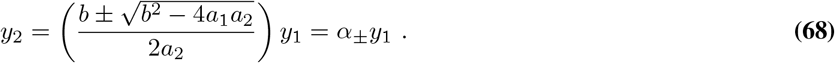

Formally, the numerator should be 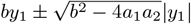, but we can absorb the absolute value into *±*, compute the solution for both *α* values, and keep the one with *y*_1_ *> y*_2_ at the end. We now insert *y*_2_ = *αy*_1_ into Eq. (65); another root *y*_1_ = *y*_2_ = 0 can be factored out, and we find the non-trivial solution

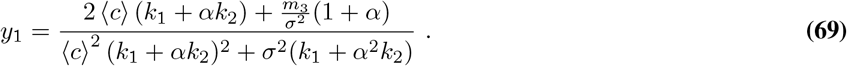

Equations 67, 68, and 69 form our analytical solution for the (approximate) fixed points of IBCM neurons in terms of the dot products *h*_*γ*_ taking values *y*_1_ and *y*_2_.

##### Condition of existence of non-trivial fixed points

The fixed points with *y*_1_ and *y*_2_ given by equations 68-69 will only exist in ℝ if the discriminant *b*^2^ − 4*a*_1_*a*_2_ in *α* is non-negative. Writing this discriminant explicitly,

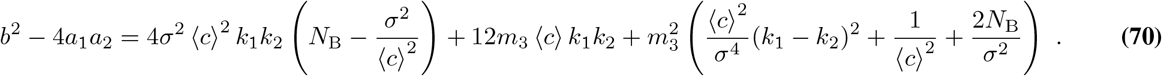

In general, this expression will be *>* 0, unless there is very high variance and low average concentration, *σ*^2^ *> N*_*B*_ ⟨*c*⟩^2^, and vanishing third moment *m*_3_ → 0. This would be an unnaturalistic setting corresponding to Gaussian, zero-average background fluctuations.

#### E. Summary of the general fixed point solutions

The fixed point solution for i.i.d. odor concentrations with mean ⟨*c*⟩, variance *σ*^2^, third moment *m*_3_, is summarized here. The fixed points of 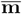 are characterized by their dot products with the *N*_B_ background odors, 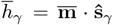. There are 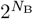 fixed points in total: one with all *h*_*γ*_ = 0, one with all *h*_*γ*_s equal to (subsection D.1)

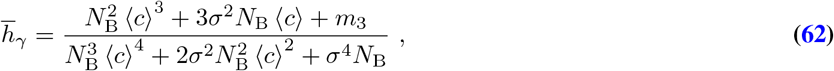

and 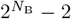 where *k*_1_ dot products are equal to *y*_1_, and *k*_2_ = *N*_B_ − *k*_1_ are equal to *y*_2_, with 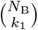 choices for each possible *k*_1_ *{*1, …, *N*_B_ − 1*}*. The values *y*_1_ and *y*_2_, where by convention *y*_1_ *> y*_2_, are calculated as follows:

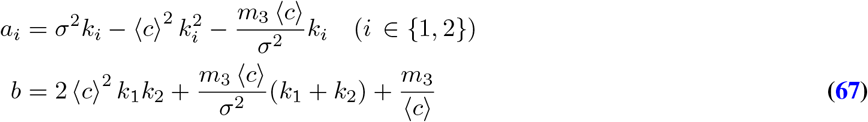

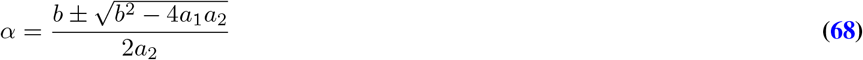

Compute solutions for each sign in *α*

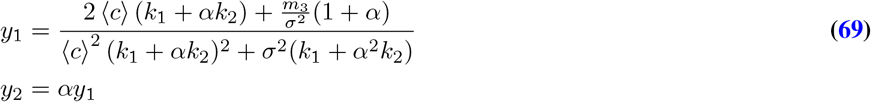

Keep the pair where *y*_1_ *> y*_2_

Our linear stability analysis, below, suggests that the *stable* fixed points of an IBCM neuron have one dot product equal to

*y*_1_ and all others equal to *y*_2_. This means the neuron becomes selective: it is specifically responding to one background odor.

There are *N*_B_ such fixed points. We can therefore call **m** *·* **ŝ**_*γ*_ = *y*_1_ ≡ *h*_sp_ (specific) for that odor, and *y*_2_ ≡ *h*_ns_ (non-specific) for all other odors. We observe close quantitative agreement between these fixed point predictions and numerical simulations in the weakly non-Gaussian background, Fig. 3C. For log-normal (Fig. S6D-E) and turbulent backgrounds (Fig. 4A-B), we also observe that IBCM neurons align with individual background odors, but due to stronger correlations between **m** and Θ, the exact dot product values do not exactly match equations 67-69.

##### G. Linear stability analysis of IBCM fixed points

To support our empirical results on IBCM neuron selectivity, we linearize the dynamical equations 51-52 around a fixed point and compute the Jacobian matrix. Then, we evaluate its eigenvalues, at least numerically, for every fixed point, and check which fixed points are stable in a number of examples. Here, we perform this analysis for a single neuron, and assume that weak coupling with other neurons in the network does not fundamentally affect the stability of single-neuron fixed points. We also assume a constant learning rate *µ*_Θ_ = *µ*, as would approximately be the case at steady-state in the Law and Cooper variant used for full simulations. Hence, the dynamical equations are

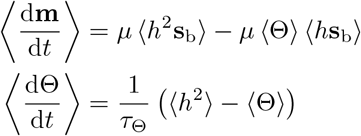

Computing the Jacobian entries, recalling that *h* = **m** *·* **s** and thus 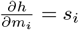, we find

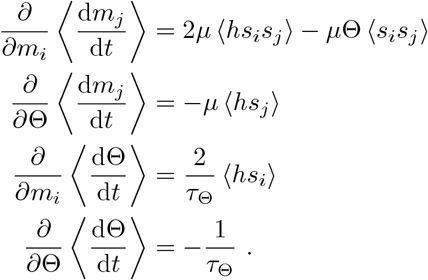

These derivatives form the different blocks of the Jacobian matrix, which is, in vector notation,

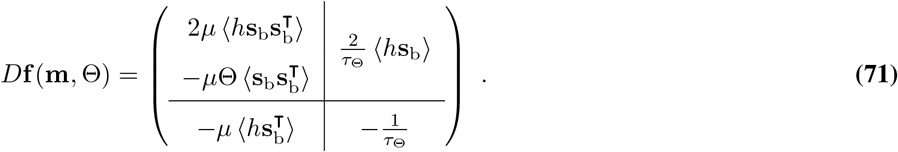

This expression is general. Now, computing more explicitly the expectation values for i.i.d. concentrations as in previous subsections,

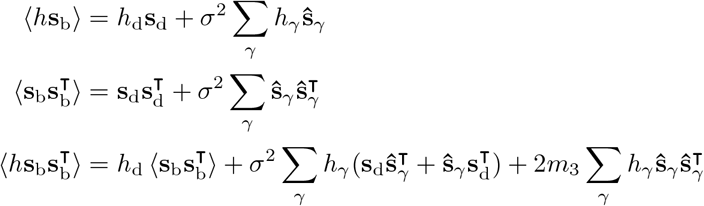

The last two lines, along with 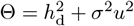, means that the main block of the matrix is

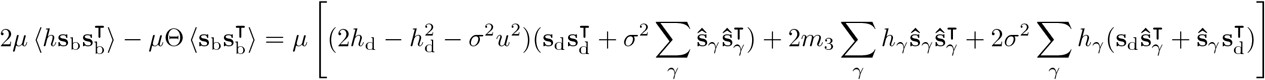

We have defined here **s**_d_ = ⟨*c*⟩ ∑_*γ*_ **ŝ**_*γ*_. We see that these moments depend on the specific odor components **ŝ**_*γ*_, making an analytical calculation of eigenvalues hard in general. However, these expressions can be evaluated easily at the analytical fixed points in several examples to check the stability in these cases; we show the eigenvalues for weakly non-Gaussian, log-normal, and turbulent background statistics in Fig. S3. In all these examples, we find that the only stable fixed points are those where the neuron has one dot product *h*_*γ*_ = *h*_sp_ (specific) and the *N*_B_−1 others are equal to *h*_ns_ (non-specific). This property is robust against OSN noise, as shown in Fig. S8. In that case, the IBCM and BioPCA models still perform habituation to the true background subspace and new odor recognition, until the OSN noise becomes comparable in magnitude with odor signals (Fig. S8H).

#### G. Analytical *W* weights with IBCM neurons

We can also derive an analytical expression for the average, steady-state values of the inhibitory weights *W* when the projection weights *M* converge to the IBCM fixed point derived above. We assume there is at least one neuron per odor, *N*_I_ ≥ *N*_B_. We call *γ*_*j*_ the background odor to which IBCM neuron *j* is specific; thus, **m**_*j*_ *·* **ŝ**_*γ*_ = *h*_sp_ if *γ* = *γ*_*j*_, *h*_ns_ otherwise. There will be in general some number *n*_*γ*_ of neurons specific to odor *γ*.

We start by working on the *W* equation in matrix form,

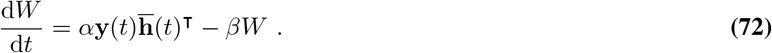

We define the (constant) matrix Γ, whose columns are the **ŝ**_*γ*_, and **c** the vector of odor concentrations. Then, **s**_b_(*t*) = Γ**c**. We also define the matrix *H* = *LM* Γ, in which row *j* gives the alignment of IBCM neuron *j* with each odor, 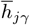. Since each neuron is selective for one odor, each row contains *h*_sp_ once and *h*_ns_ *N*_B −_ 1 times. Averaging the *W* equation over fast *c* fluctuations, and neglecting correlations between *c, W*, and *H*, we have

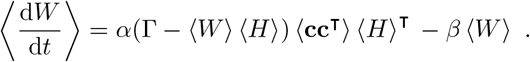

Average signs on *W* and *H* are implied below. We define *N* = ⟨**cc**^⊤^ ⟩ and evaluate it for i.i.d. odors,

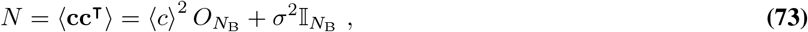

where 𝕀 is the *N*_B_ *× N*_B_ identity matrix and 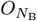 is a *N*_B_ *× N*_B_ matrix filled with ones. We now set d*W/*d*t* = 0 and focus on single columns of the equation, with the notation **w**_*j*_ for column *j* of *W*, and **h**_*j*_ for column *j* of *H*^⊤^ (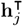 is the row *j* of *H*).

The set of equations to solve for the **w**_*j*_ is then

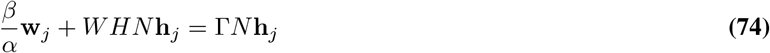

We notice here that all IBCM neurons with the same specificity *γ*_*j*_ will have the same **h**_*j*_, thus all columns **w**_*j*_ with the same *γ*_*j*_ will be identical, and we can denote them by **w**_*γ*_. This allows to rewrite sums over columns as sums over components, for instance ∑_*j*_ **w**_*j*_ = ∑*n*_*γ*_**w**_*γ*_. We moreover notice that

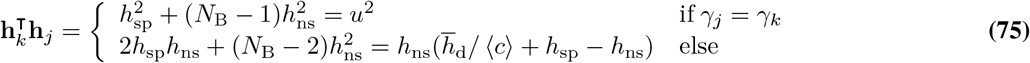

and that

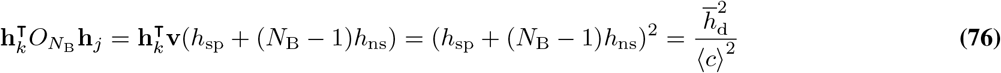

where we defined **v**, a vector filled with ones, and recognized 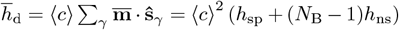 for all neurons specific to one odor. We also need to compute

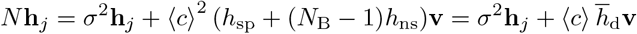

where we used Eq. (73). We use the above results to evaluate the two terms involving *N* in Eq. (74). With some algebra to combine coefficients efficiently, we find

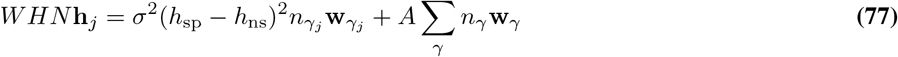

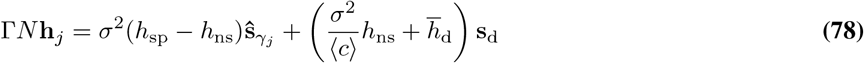

where we have used the average background expression, 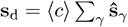, and defined a coefficient independent of *j*,

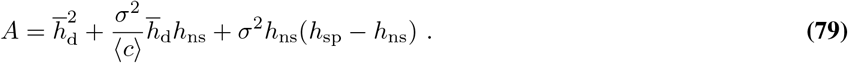

We now insert Eq. (77) and Eq. (78) into the equation Eq. (74) for **w**_*j*_ (or equivalently, 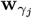), to find

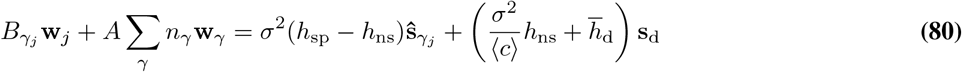

where we have defined another coefficient, different for each *j* in general, unless all 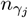are equal,

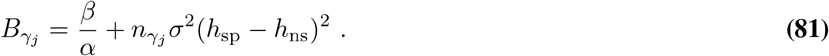

Now, to solve Eq. (80) for 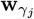, we look at the difference between the equations for columns *j* and *k*, with *γ*_*k*_ ≠ *γ*_*j*_; this eliminates common terms and allows us to isolate 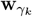 in terms of 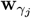 and the known vectors **ŝ**_*γ*_ only:

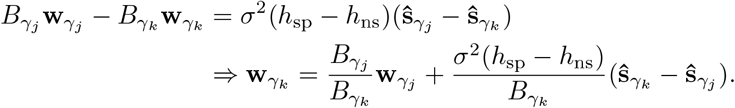

Doing this for each *γ*_*k*_ ≠ *γ*_*j*_, and noticing the expression reduces to 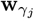 for *γ* = *γ*_*j*_, we express 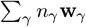 in terms of 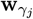 only, and insert into eq. Eq. (80) to isolate 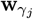. Dividing the result by 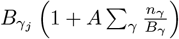, and rearranging with further algebra gives our final expression for columns of the matrix *W*,

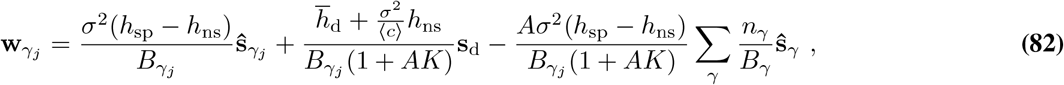

where we have defined 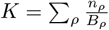. These are the analytical expressions that we compare to numerical simulations of the *W* weights in Figures 3D (weakly non-Gaussian) and S6F (log-normal), reaching close agreement at steady-state. For backgrounds with slower, stronger fluctuations where the correlations between *M, W*, and **s**(*t*) are not entirely negligible (*e*.*g*., turbulent statistics), the agreement would be less accurate. The numbers of neurons selecting each odor, *n*_*γ*_*j*, are inferred from the *M* weights numerical results, then used to evaluate Eq. (82).

Unfortunately, the expression for the instantaneous **y**(*t*) = **s**(*t*) − *W LM* **s**(*t*) with steady-state *M, M* weights is quite cumbersome in general, due to the *B*_*γ*_. However, we find a more compact expression for the average PN response: after simplifying some terms,

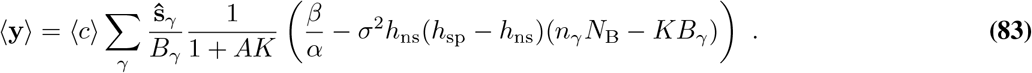

The factor *n*_*γ*_*N*_B_ − *KB*_*γ*_ is zero when all *n*_*γ*_s are equal; hence, the second term represents the bias incurred by having an uneven distribution of IBCM neurons across odor components. The threshold *n*\* at which *n*_*γ*_*N*_B_ − *KB*_*γ*_ = 0 for some *n*_*γ*_ is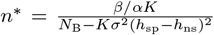 ; since we can show (using a Lagrange multiplier to enforce 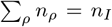) that *K* is maximized by having a uniform distribution *n*_*γ*_ = *n*_*I*_ */N*_B_, the threshold *n* < n*_*I*_ */N*_B_; hence, all components which have *n*_*γ*_ *> n*_*I*_ */N*_B_ surely have a positive factor (*n*_*γ*_*N*_B_ − *KB*_*γ*_), and since *h*_ns_ *<* 0 in general, they have a negative bias in eq. Eq. (83), i.e. they are suppressed less than other background odor components. Conversely, if some *n*_*γ*_ = 0 (no neuron specific to that odor), then this odor is still partly subtracted, due to the non-specific response of other neurons, *h*_ns_ *<* 0, but at the cost of less efficient inhibition of all other odors, and without overall reduction of fluctuations since the factor 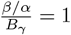 if *n*_*γ*_ = 0.

When all *n*_*γ*_s are equal, the mean PN response to the background, Eq. (83), is minimized for the IBCM network, and it reduces to a simple expression,

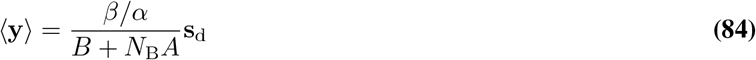

where, recall, *A* is given by Eq. (79), *B* by Eq. (81) with all *n*_*γ*_ equal, and 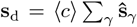 is the average background. We will use this simplified expression of the maximal habituation by IBCM neurons determine the Λ factor needed to match the BioPCA and IBCM performances (section 8B).

### 5. Analytical fixed point solutions of the BioPCA model

We assume that the average background 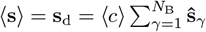 is subtracted from **s**, from the input to the BioPCA inhibitory neurons, and from **y**; hence, the effective background to consider here is 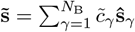, with 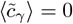 and 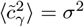 for all components *γ*. The LN activity is 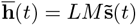 and the PN response is 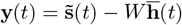. The covariance matrix, from which a PCA with *N*_B_ components can be computed, is

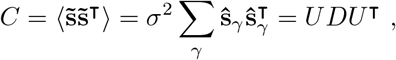

where *U* is *N*_s_ *× N*_B_, with its columns containing the *N*_B_ principal component vectors with non-zero eigenvalue, and *D* is *N*_B_ *× N*_B_ and diagonal with the principal values in it, 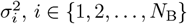. Since the **ŝ**_*γ*_ are not orthogonal in general, these eigenvalues are not all equal to the variance *σ*^2^, but should be on the same scale.

Given a background, *U* and *D* are known; we can thus express the steady-state solution of the BioPCA model in terms of these PCA matrices. From Lemma 3 in Minden *et al*., 2018 [47], we expect the BioPCA model with *N*_I_ = *N*_B_ neurons (one per background component) to have the following stationary solution:

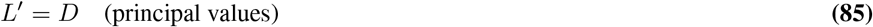

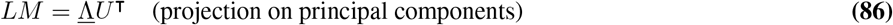

where, recall, *L*′ = *L*^−1^. The input projections give interneuron activities of

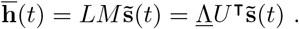

We insert the BioPCA stationary solution into the *W* equation 72, where we average over fast 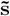 fluctuations, assuming a separation of time scales between these fluctuations and slow *M, L, W* learning, as in the IBCM case. Writing the *W* equation in matrix form, this leads to

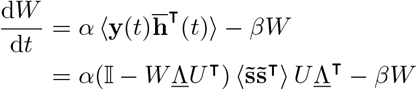

Setting the *W* derivative to zero, we can rearrange to isolate *W*,

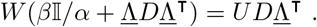

Since *D* åand <u>Λ</u> are both diagonal, the matrix in parentheses on the left-hand side is diagonal with entries 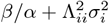. It is full-rank when *N*_I_ ≤ *N*_B_, so we can invert the equation explicitly. Since *D*<u>Λ</u>^T^ is also diagonal, we find

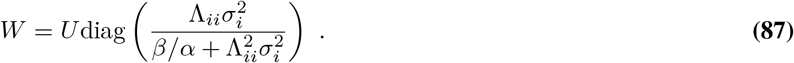

Inserting this *W* back in the expression for the PN response, 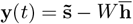, we find

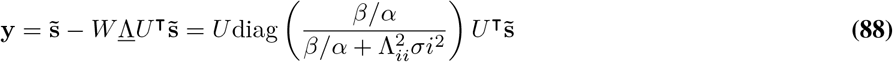

where we have used the fact that *UU*^T^ is a projector on the background subspace to write 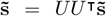, and where Λ_*ii*_ = Λ_PCA_(1 − *λ*_*r*_(*i* + 1)*/N*_B_) for *i* = 0, 1, …, *N*_I_. Hence, we see that the BioPCA network projects the inputs on the principal directions 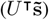, reduces the amplitude of each component by a factor 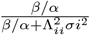, then reassembles these components (leftmost *U*). Comparing to equation 84 for the IBCM model, the latter has a better reduction by a factor of approximately (*h*_sp_ − *h*_ns_)^2^ in the denominator, hence we need to increase the Λ scale in the BioPCA network to match the performance of these models, as explained in section 8B. We check some of these predictions against numerical simulations in a background with log-normal (Fig. S6B-C) and turbulent (Fig. 4C-D) concentration statistics. We also check that these properties are relatively robust against OSN noise (Fig. S8); the first *N*_B_ neurons still capture odor directions corresponding to real odors, while additional neurons align with orthogonal OSN noise components (which are part of the full background PCA decomposition).

### 6. Analytical results for a two-odor simplified background process

To gain further analytical insight into the convergence dynamics of IBCM neurons in particular, we study the simplest non-trivial background, illustrated in Fig. S4A-B. It consists of two odors (**s**_a_, **s**_b_) with fluctuating proportion 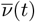 following a Ornstein-Uhlenbeck process (section 2B),

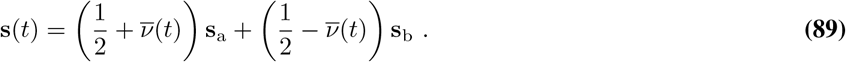

We start by calculating the average fixed points of the IBCM neurons synaptic weights,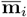. The fixed point equations 54 are identical for all neurons, so we focus on one neuron and omit index *i*. As in section 4, we assume that time scales are well separated, replace Θ = ⟨ *h*^2^⟩, and neglect correlations between 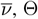 and **m**. We work with reduced weights and activities 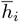, averaged over fast fluctuations, so overlines and are ⟨ ⟩ implied for the rest of the section. Individual neurons’ weights **m** can be recovered from equation Eq. (55). Moreover, for this simple background, the learning rate can be chosen constant,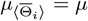.

Hence, the fixed point equation to solve here is

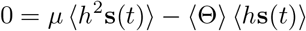

Now, we rewrite

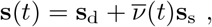

where we have defined 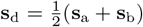 (deterministic part) and **s**_s_ = **s**_a_ *−* **s**_b_ (stochastic part). We examine the dot products of synaptic weights with these components, *h*_d_ = **m** *·* **s**_d_ and *h*_s_ = **m** *·* **s**_s_, such that 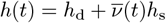. We can solve for the two dot products *h*_d_ and *h*_s_ because they specify the fixed points completely for *N*_B_ = 2 background components. In term of these quantities, the fixed point equation becomes

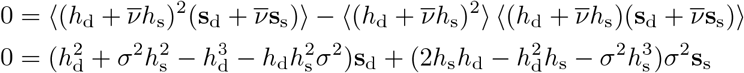

Since **s**_d_ and **s**_s_ are linearly independent, both coefficients must be zero, leading to a system of two equations for *h*_d_ and *h*_s_,

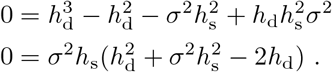

There is a trivial solution *h*_s_ = *h*_d_ = 0, which is unstable. The other solutions are, by inspection,

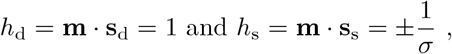

or, in terms of the dot products with **s**_a_ and **s**_b_,

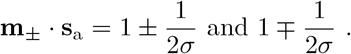

Hence, we have two different stable fixed points, which we call **m**_+_ and **m**_*−*_ to indicate which sign the dot product with **s**_s_ takes. Figure S4C shows the convergence of a two-neuron network to these fixed points. To interpret these expressions, consider the response at the fixed point to some input sample **s**(*t*):

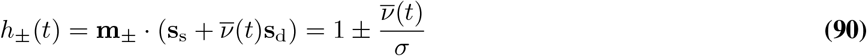

We notice that *h*_*±*_ = 0 when 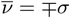, that is, the IBCM neuron is non-responsive to an odor component one standard deviation away on one side of the average background, while it responds strongly to odors on the other side of the average. Hence, in this simplified background, the specificity property of IBCM neurons translates into selecting inputs one standard deviation away from the average.

#### A. Analytical results: PN inhibition

From the steady-state solution for **m**, we can also compute the steady-state inhibitory weights **w**. We assume there are *N*_I_ = 2 neurons, one at each fixed point *±*.

Averaging the *W* equation 72 over background fluctuations, writing out **y** = **s** *− W LM* **s**, and focusing on the column for one neuron *j*, we have

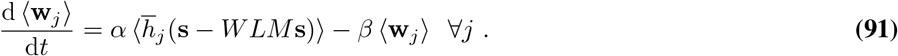

To solve for **w**_*j*_, we set the derivative to zero, and we assume there are *N*_I_ = 2 IBCM neurons, one at each fixed point *±*. We still assume a separation of time scales, and assume the 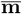 are equal to their average fixed point values, so at any time *t*, the IBCM neuron activity 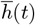 is given by Eq. (90). We assume the two neurons converge to fixed points + and *−*, respectively. We thus establish equations to solve for the **w**_*j*_ weights fixed point values, **w**_+_ and **w**_*−*_. We have

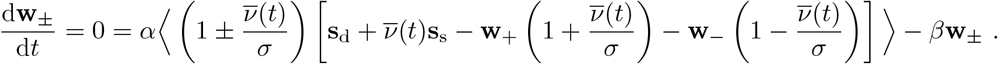

Solving for **w**_+_ and **w**_*−*_, we find answers summarized as

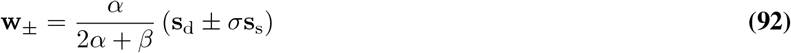

Hence, each IBCM neuron inhibits the off-average component for which it is selective,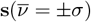. Combining the two IBCM neurons, the instantaneous PN activity is reduced to

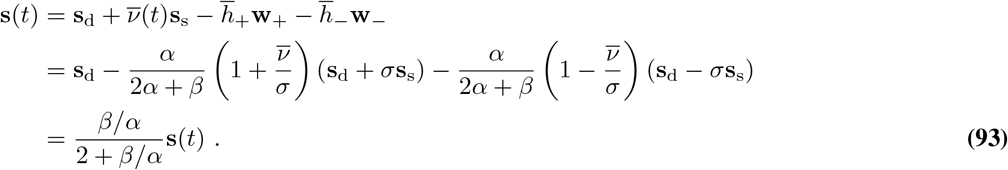

Figure S4F-G show close agreement at steady-state between numerical simulations and equations 93 and 92. Hence, by learning *N*_B_ = 2 linearly independent components **w**_*±*_ that are one standard deviation away from the average background, the network is able to suppress any **s**(*t*) from that background, in real time, to a fraction 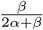 of its original amplitude. Therefore, not only the average, but also the variance of the background is reduced: background fluctuations are actively suppressed by the IBCM-inhibitory neuron pairs. However, new odors would not be suppressed in the same way, because they have a component orthogonal to the vector space of learnt background.

#### B. Analytical results: convergence time

The convergence time of the **m** weights of an IBCM neuron can be estimated analytically in the two-odor simplified background; this analysis reveals the main parameters influencing how long it takes to habituate to a fluctuating background. Numerically, we observe that with the *α, β* rates chosen, *W* weights converge at a similar pace.

We again make a quasi-static approximation on the threshold Θ, assuming it averages over the fast background fluctuations but also converges fast enough to track the slow variations of **m**,

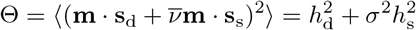

where we made use of 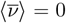 and 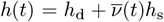. Then, we derive dynamical equations for the slow variables *h*_d_ and *h*_d_, by taking the dot product of 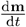 with **s**_s_ and **s**_d_, averaging over fast time scales of 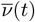, and using the quasi-static Θ above. To simplify calculations, we assume that the two odor vectors, **s**_a_ and **s**_b_, have the same norm (like the **ŝ**_*γ*_ in the general case are unit normed); in this case, the vectors **s**_d_ and **s**_s_ are orthogonal. Making use of these properties, we calculate for instance, for *h*_d_,

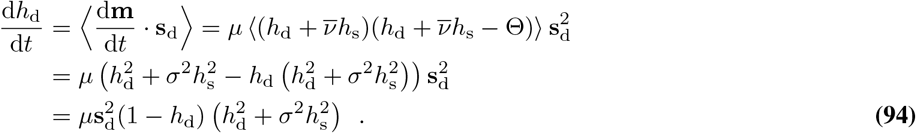

By a similar calculation, we find for *h*_s_

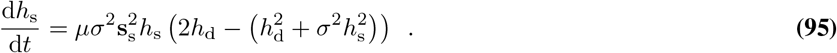

From equations Eq. (94) and Eq. (95), we can conclude there will be two phases to the dynamics if the initial values of *h*_s_(0) = *E*_s_ and *h*_d_(0) = *E*_d_ are small, and *σ*^2^ is small also. The only positive term in 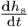 contains *h*_d_; hence, as long as *h*_d_ is small, *h*_s_ will remain close to zero. The first phase therefore consists in the growth of *h*_d_ to its steady-state value of 1, while *h*_s_ remains approximately equal to its initial value, *E*_s_. We call *t*_d_ its duration. After *h*_d_ has converged, the second phase consists in the growth of *h*_s_. We call *t*_s_ the duration of that phase. Hence, *h*_s_ reaches steady-state after a total time of *t*_d_ + *t*_s_.

We compute *t*_d_ (first phase duration) by integrating equation Eq. (94) from 0 to some fraction *ξ* (close to unity; we use *ξ* = 0.9 in practice) of the steady-state *h*_d_ = 1, with the assumption that *h*^2^ is approximately constant and sub-dominant in that phase, i.e. 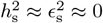. We find

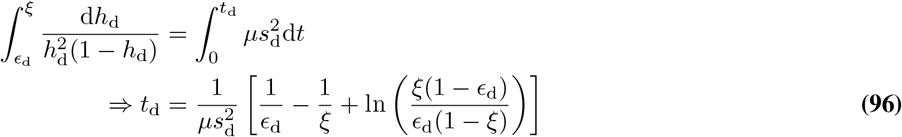

where 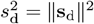.

Then, once *h*_d_ *≈* 1, the second phase starts. We neglect sub-dominant terms and we integrate from *t*_d_ (time at which *h*_d_ *≈* 1 but *h*_s_ *≈ E*_s_ still) to *t*_d_ + *t*_s_. We integrate *h*_s_ from *ϵ*_s_ to *±ξ/σ*: depending on the sign of the initial value *ϵ*_s_, the system goes to either fixed point *±*1*/σ* (same sign as the initial value). Hence,

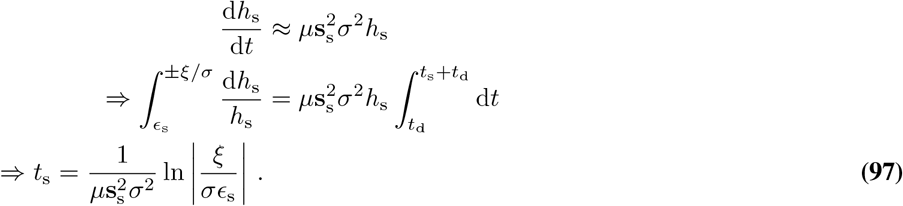

Fig. S4D and E show that the approximations Eq. (96) and Eq. (97) hold well in a range of initial values *ϵ*_s_ between 0.005 and 0.05, and *ϵ*_d_ between 0.01 and 0.1. Importantly, these analytical expressions show that convergence time is faster if initial conditions are larger (note inverse *ϵ* terms) and the noise *σ*^2^ is larger. It means that initial conditions must be chosen to avoid very small dot products with inputs initially. Then, larger background fluctuations trigger faster convergence too, at least in this simple background model. This is a surprising property of the IBCM model: fluctuations drive the dynamics.

#### C. BioPCA neuron on the two-odor simplified background

We also characterized the behavior of the BioPCA model in this simplified background. Since the background effectively has one principal direction, **s**_s_ (Fig. S7A), a single BioPCA neuron is needed to capture it. Then, the matrix *M* is only a row vector **m**^T^ and the matrix *L* is only a scalar *𝓁*, so *L′* = *𝓁 ′* = 1*/𝓁*. Likewise, *W* is a column vector **w** and the matrix <u>Λ</u> is just the scalar parameter Λ. Since we assume BioPCA receives the input with the average subtracted, the background we consider is 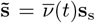. We illustrate how to solve the BioPCA steady-state equations in this simple case. With 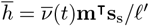, the dynamical equations simplify to

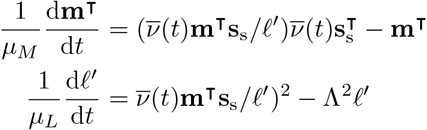

Averaging over 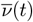 and setting the derivatives equal to zero, we have two fixed point equations,

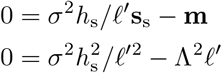

where we have defined *h*_s_ = **m**^T^**s**_s_. The first equation shows that **m** is parallel to the first principal component: **m** = ∥**m**∥ **s**_s_*/*∥ **s**_s_∥. To find its magnitude, we take the dot product of that equation with **s**_s_, which allows to factor out *h*_s_ and solve for *𝓁 ′*. We find that *𝓁 ′* does converge to the first principal eigenvalue, which is *σ*^2^ ∥**s**_s_∥^2^ in this simplified background,

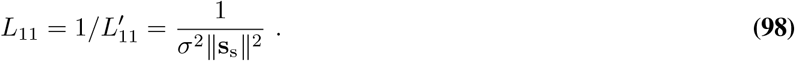

From the second equation, we then find *h*_s_ = Λ*σ*^2^ ∥**s**_s_∥^3^, which means that **m** has a norm ∥**m**∥ = ∥Λ*σ*^2^∥ **s**_s_∥^2^. Hence, we have **m** parallel to the first principal component,

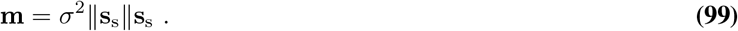

Also, the instantaneous response of this neuron to 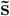 is 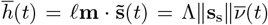. Inserting these expressions in the **w** equation for this single neuron,

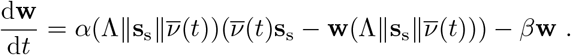

Averaging over 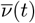 and solving for **w**, we find that it is also a vector parallel to **s**_s_,

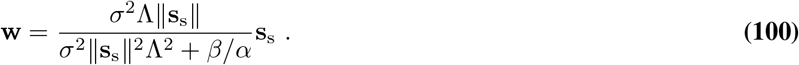

Lastly, computing the PN instantaneous activity once the BioPCA neuron has reached its learning fixed point, we find

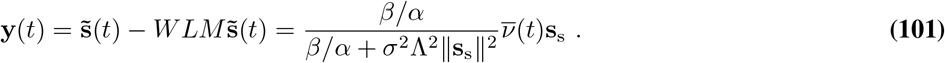

Fig. S7 shows that these analytical predictions for **m**, *L*, **w**, and **y**(*t*) match numerical simulations very well. Hence, by learning the direction of fluctuations along the first principal component, the BioPCA neuron reduces the mean and standard deviation of fluctuations by a factor 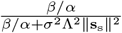, to be compared with the reduction achieved by the IBCM network, in equation 93.

### 7. Testing different *L-*norms for the *W* Hebbian learning rule

The Hebbian learning rule for *W* used in the main text (derived in *Methods*) causes our proposed models to subtract the entire component of the input lying in the background subspace. It gives them a sub-optimal performance, limited to the similarity between the new odor and its orthogonal component (Figures 5, S10). To start exploring alternative *W* rules that could better exploit the odor-specific projections learnt by IBCM neurons, we considered the effect of using different *L*^*p*^-norms in the cost function from which the *W* dynamics are derived. We write a cost function for *W* weights based on the *L*^*p*^-norm of PN activity and the entry-wise *L*^*q*^-norm of *W* (generalization of the Frobenius norm, which corresponds to *q* = 2), defined as

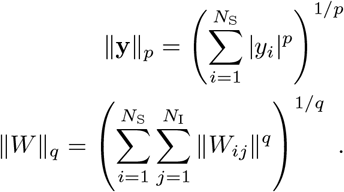

In the cost function, we square terms to preserve direct comparisons with the default *L*^2^-norm cost function:

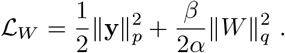

Taking gradient descent dynamics on this loss function gives a generalized Hebbian learning rule,

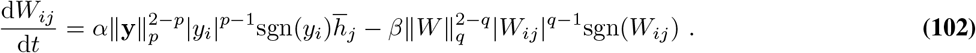

Notice that it reduces to the main text rule when *p* = *q* = 2. We performed numerical experiments of habituation and new odor recognition, analogous to Fig. 2, for various *p, q* choices in this generalized Hebbian rule. For each (*p, q*) choice, we optimized the performance with a grid search over a few *α* and *β* learning rate values, centered on relevant windows (*e*.*g*., a smaller *p* requires a smaller *α* to prevent numerical instabilities). Unfortunately, different *p, q* choices did not fundamentally alter the model performance for habituation (Fig. S9A-C) or new odor recognition (Fig. S9D-F); in fact, the default *L*^2^-norm provided the best results. Hence, more strongly nonlinear versions of manifold learning, such as online manifold tiling [57], or learning rules with positive feedbacks to learn the optimal matrix *P* of section 1, would be needed to further improve new odor recognition performance.

### 8. Projection weights scale factor, Λ

As explained in *Methods*, we defined a parameter Λ controlling the scale of *M* weights and compensating for the regularization on *W*. Here, we explain how to introduce Λ in the IBCM model, and how to set this parameter to make BioPCA and IBCM perform equivalently.

#### A. Introducing a scale factor Λ in the IBCM model

We start with the equations for a single neuron. We seek to introduce Λ where appropriate in the equations to maintain the exact same dynamics, only with the numerical values of **m** weights (including their initial values) scaled by some factor Λ. By definition, as we scale **m** *∼* Λ, then *h* = **m** *·* **s** *∼* Λ as well. The IBCM equation contains terms of the form *h −* Θ, with Θ *∼ h*^2^; to keep these terms matched, we need Θ *∼* Λ, which we achieve by letting Θ *→ h*^2^*/*Λ. Hence, we start by modifying the threshold equation to

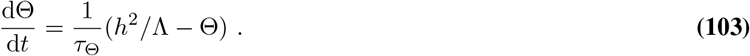

We do not need to rescale the learning rate here, because both sides have a homogeneous scaling *∼* Λ. However, in the **m** equation, we need to rescale the learning rate to preserve the dynamics, because the right-hand side has terms *∼ h*^2^. To keep both sides scaling as 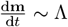, we modify the **m** equation to

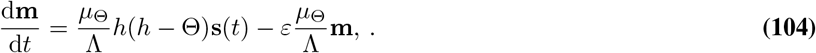

Moreover, since the scale of Θ *∼* Λ, we need to rescale it in the Θ-dependent learning rate (from our variant of the Law and Cooper version of IBCM),

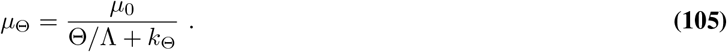

The generalization to a network of IBCM neurons is straightforward, since Λ is a unique scale parameter for all neurons. Each 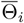 equation has the rescaled term 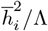 as in 103. All terms in neuron *i*’s **m** equation, including those from coupling with other neurons *j*, have their learning rates 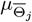 divided by Λ, as in 104. Also, 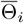 is divided by Λ in the denominator of each learning rate 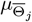 as in eq. 105.

These are the IBCM equations we use for general Λ values. In Fig. S11, we characterize the performance of the network for habituation and new odor recognition as as function of the scale Λ, and we observe that Λ_IBCM_ = 1 is large enough to maximize the performance.

#### B. Scaling the BioPCA model for performance equivalent to IBCM

As explained in *Methods*, the scale Λ is already built into the BioPCA model, in the matrix <u>Λ</u> intervening in the *L* equation. Λ_PCA_ is set to 1 by default [47], but this leads to a smaller *M* weights magnitude than by default in the IBCM model. For this reason, Fig. S11 shows that the BioPCA model requires a Λ_PCA_ *>* 1 to achieve the same performance.

For other simulations where we compare the two models, we use our analytical results on the IBCM and BioPCA models to estimate beforehand what Λ_PCA_ value should yield a comparable performance from both models. As derived in Eq. (84), the PN response is reduced, in an IBCM network, by a factor

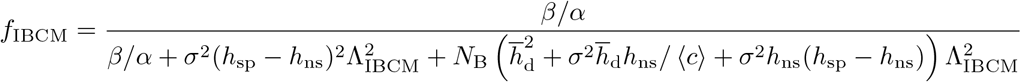

where we have explicited how Λ_IBCM_ controls this factor by multiplying the default-scale LN activities, 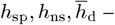 recall that these are the specific, non-specific, and average alignments of the IBCM fixed points, derived in section 4.

In comparison, Eq. (88) shows that the PN activity in a BioPCA network is reduced along each principal direction by a factor of approximately

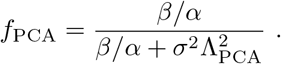

We set Λ_PCA_ to make these two factors equal, which occurs at

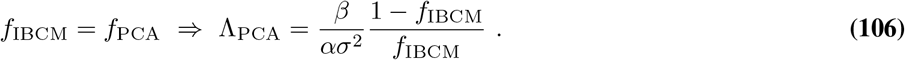

So, to set up parameters for a numerical simulation, we compute the statistics of the chosen background (⟨*c*⟩, *σ*^2^, *m*_3_), the analytical predictions for the IBCM fixed points 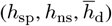, and we set Λ_PCA_ to these parameter values inserted in Eq. (106).

**Fig. S1.**
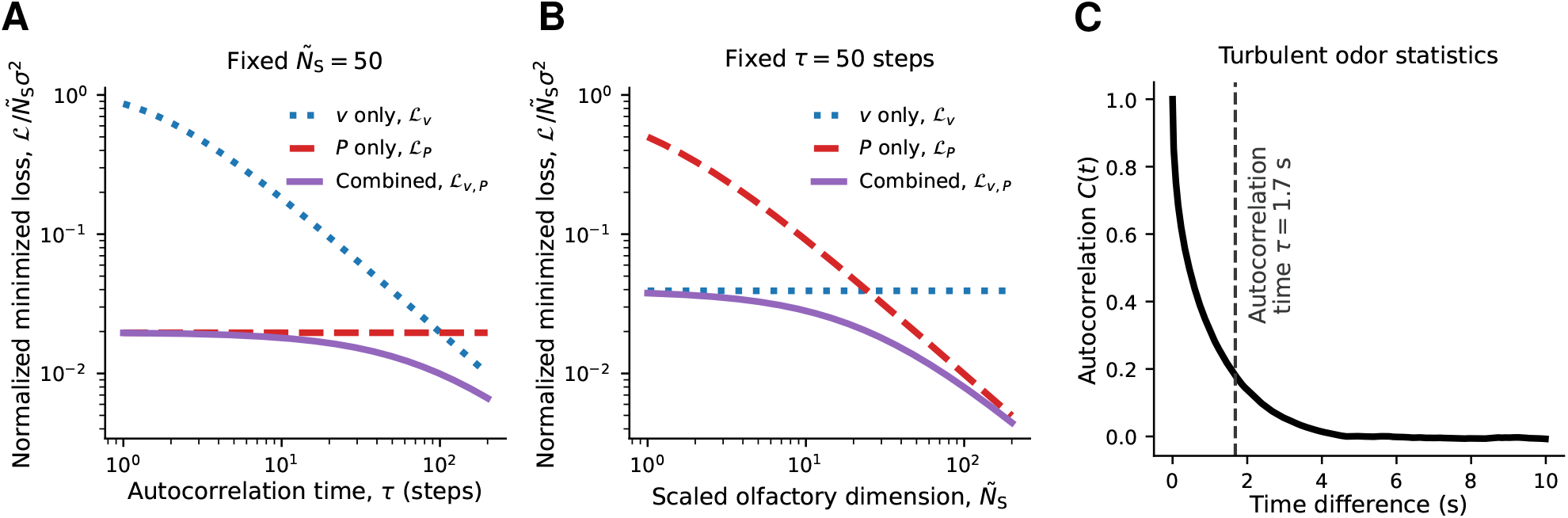
Comparing manifold learning and predictive filtering. Supplement to Fig. 1E. (**A**) Minimized loss function for new odor recognition in a simple background (Ornstein-Uhlenbeck), for the combined strategies (purple) and either single strategy (blue, red), as a function of the autocorrelation time *τ*, for fixed olfactory space dimensionality. The combined strategy is always better, but manifold learning explains essentially all the loss reduction at low autocorrelation times (ℒ_*P,v*_ *≈* ℒ_*P*_). (**B**)_new_ Same, as a function of the scaled olfactory space dimension, 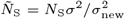, for fixed autocorrelation time *τ* = 50 steps. Most of the loss reduction comes from one strategy or the other on either side of the crossover region; manifold learning dominates in high dimensions. (**C**) Autocorrelation function of the concentration fluctuations in the turbulent background of Fig. 1B-C, showing an autocorrelation time of *∼* 1.7 s.

**Fig. S2.**
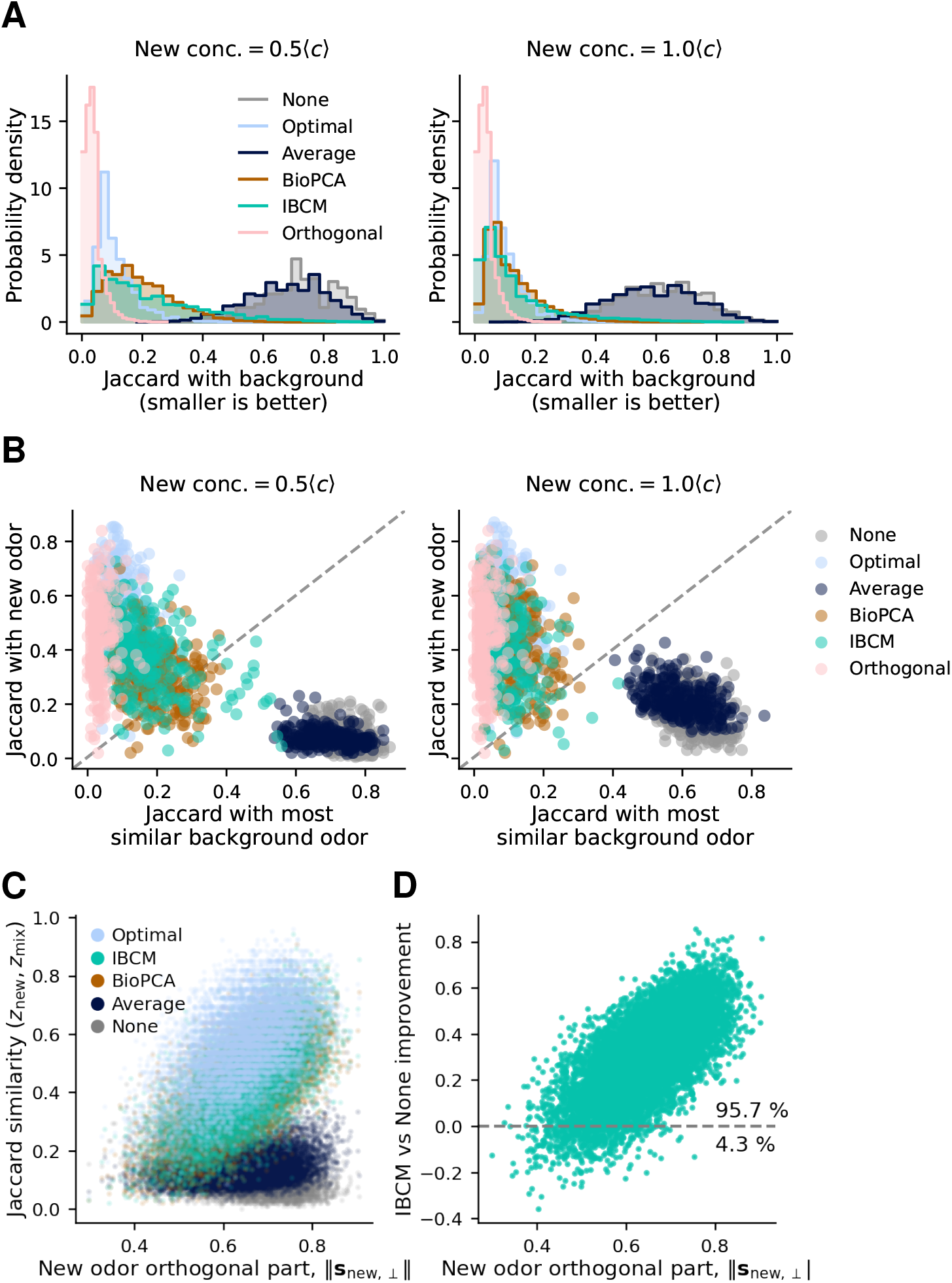
New odor recognition compared to background recognition after habituation. Supplement to Fig. 2. (**A**) Histograms of the Jaccard similarity between the response to the mixture (background and new odor) and the most similar background odor, across all repeats (new odors, background samples, etc.) described in Fig. 2A, for two tested new odor concentrations. Low similarity is better: it means that background odors are suppressed and do not dominate the response to the new odor, due to habituation. In the absence of habituation, the similarity to background odors remains high. (**B**) Scatter plots of the similarity to the new odor (y axis, larger is better) versus the similarity to the background (x axis). Manifold learning models are generally above the diagonal, meaning their response is more similar to the new odor than to the background, while habituation by average subtraction (as well as no habituation) produce responses still dominated by the background fluctuations (below diagonal). (**C**) Jaccard similarity between *z*_mix_ and *z*_new_ plotted as a function of the magnitude of the new odor component orthogonal to the background (or equivalently, of the distance between the background odors and the new odor), in each trial. For all models, the performance improves as the new odor is less similar to the background. (D) For the IBCM model, improvement in Jaccard similarity compared to no habituation (*J*_IBCM_ *− J*_None_). Habituation by manifold learning almost always (*>* 95 % of trials) provides an improvement, except for rare background samples (*<* 5 %) where background odors are all in blanks or in very dilute whiffs (then the new odor, by chance, is not masked by the background). In (B), (C), and (D), each point is the median across test times and background samples in each individual background simulation (of which there are 100 repeats).

**Fig. S3.**
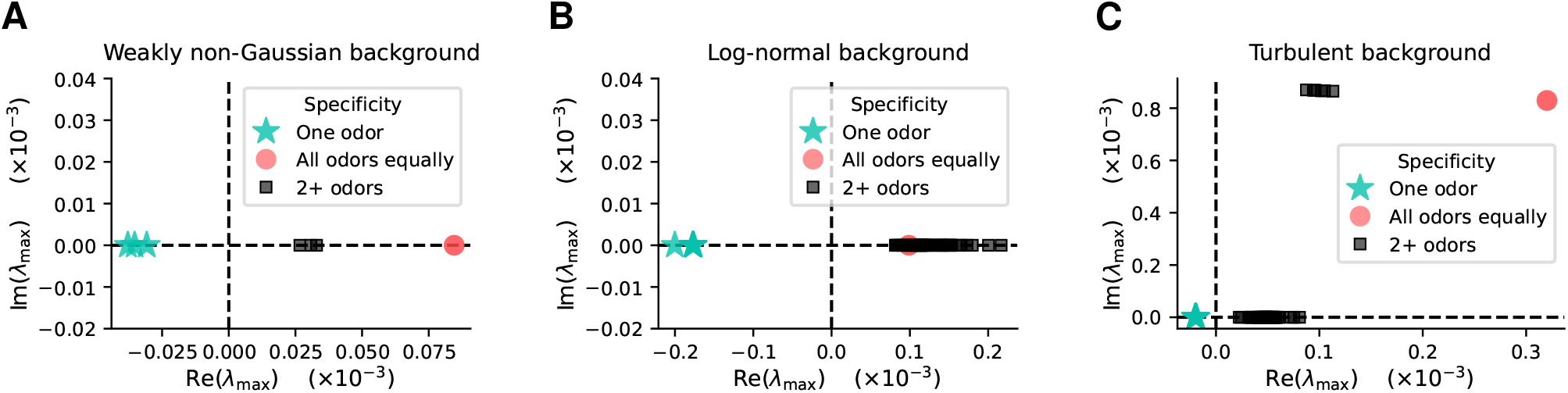
Selective states are the only stable fixed points in the IBCM model. Eigenvalue with the largest negative real part in the Jacobian matrix obtained in the linear stability analysis of the IBCM model (section 4F), evaluated for each possible fixed point in (**A**) the weakly non-Gaussian background of Fig. 3, (**B**) the log-normal background of Fig. S6, and (**C**) the turbulent background of Figs. 1B-C, 2, 5. The Jacobian matrix derived in eq. 71 is diagonalized numerically for each possible choice of specific and non-specific odors. In the three background examples considered, the only stable fixed points, *i*.*e*., fixed points where even the largest eigenvalue has a negative real part, are those where the neuron is specific to only one odor, *i*.*e*., selective states. The saddle points where the neuron is equally sensitive to all odors, in red, have some eigenvalues with positive real parts (largest is shown), and some with negative real parts.

**Fig. S4.**
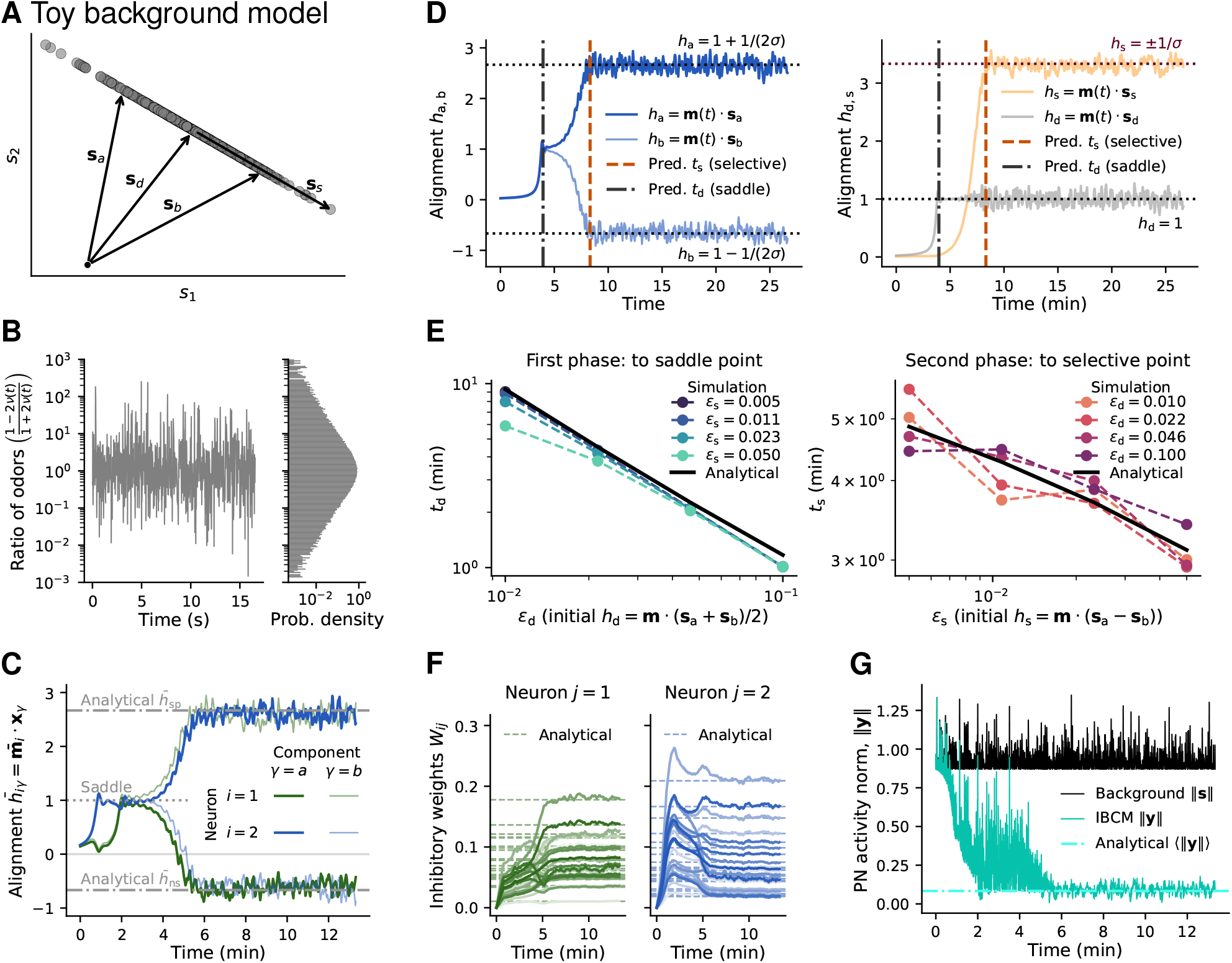
IBCM model learning dynamics on a simplified background model. (**A**) Schematic of the simplified background model described in section 6: a 1D manifold generated by two odors **s**_a_, **s**_b_ with fluctuating proportion 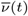 following a Ornstein-Uhlenbeck process with 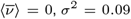, autocorrelation time *τ*_b_ = 20 ms. The deterministic part is 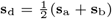 and the stochastic part, **s**_s_ = **s**_a_ − **s**_b_. The two odors are randomly sampled from the default distribution (exponential iid elements, then normalized). (**B**) Time series and stationary distribution of the ratio of odors a and b. (**C**) Alignment of the two IBCM neurons with the background odors. One neuron becomes specific to **s**_a_, the other, to **s**_b_. The average dot products at steady-state match the analytical fixed points calculated in 90, 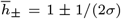. (**D**) Analysis of the two-phase dynamics of an IBCM neuron. Left: visualized in terms of the alignments (dot products) with odors a and b, **m** first reaches a saddle point, then one of the two selective stable fixed points, respectively at times *t*_d_ (eq. 96) and *t*_s_ closely matching their analytical values (equations 97 and 96). Right: visualized in terms of the alignments *h*_d_ and *h*_s_ with the deterministic and stochastic components, **s**_d_ and **s**_s_, the saddle at time *t*_d_ corresponds to the time at which the deterministic part *h*_d_ reaches its fixed point average value 1, and the final steady-state, when the stochastic part reach its fixed point value *±*1*/σ*. (**E**) Scaling of the convergence times *t*_d_ (left) and *t*_s_ (right) as a function of the initial dot product magnitudes, ϵ_d_ and ϵ_s_. The numerical simulations match the analytical expressions, valid for small ϵs. (**F**) Time series of the inhibitory weights (elements of *W*), following the Hebbian dynamics of eq. 91, for the two neurons in the network (weights of LN *j* are in column *j* of *W*). The steady-state values match closely the analytical values derived in eq. 92. (**G**) Reduction in PN activity after habituation, compared to the background norm. The steady-state reduction matches the analytical prediction (eq. 93) and is reached once the two neurons select one odor each.

**Fig. S5.**
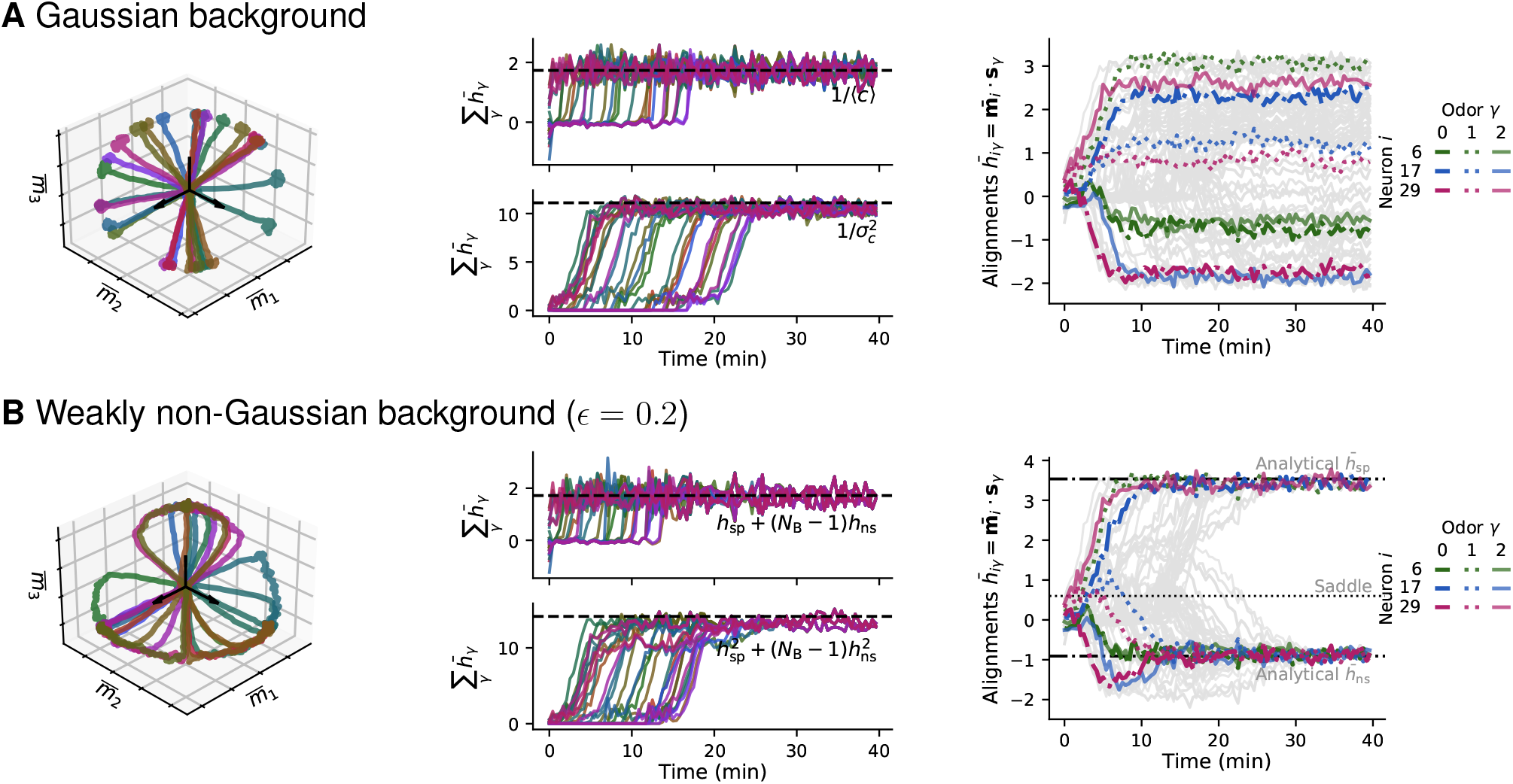
IBCM learning depends on higher-oder moments of the background distribution. (**A**) Three-odor background with Gaussian concentration fluctuations: each odor has *c*_*γ*_ = *g*_*γ*_ where *g*_*γ*_ follows a Ornstein-Uhlenbeck process (with 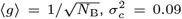, correlation time *τ* = 20 ms). Left: when the third moment is zero, synaptic weights 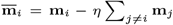 have non-isolated fixed points on the codimension-two ring defined by constraints Eq. (59) (hyperplane) and Eq. (60) (hypersphere). The figure is showing the first three dimensions. Center: All neurons converge to these constraints. Right: alignments with individual odors can take a continuous range of values and do not split into specific and non-specific odors for each neuron. (**B**) Same as (A), but with a small non-zero third moment added to the background concentrations statistics, by taking *c* = *g* + ϵ*g*^2^ for ϵ = 0.2, leading to ⟨ (*c −* ⟨ *c*⟩)^3^⟩ ≈ 0.01. Left: the fixed points become isolated, individual neurons first converge to the codimension-two ring of the Gaussian case, then slowly approach one of three selective fixed points near that ring (driven by the small third moment). Center: for each neurons, the sums of 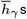 and 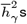 have similar values to the Gaussian case, perturbed by the small third moment. Right: alignments converging to selective fixed points with dot product values *h*_sp_ (with one odor), *h*_ns_ (with *N*_B_ *−* 1 odors); three neurons highlighted. Odor vectors were chosen to be symmetric around the origin in the first three dimensions, to clarify the geometric picture.

**Fig. S6.**
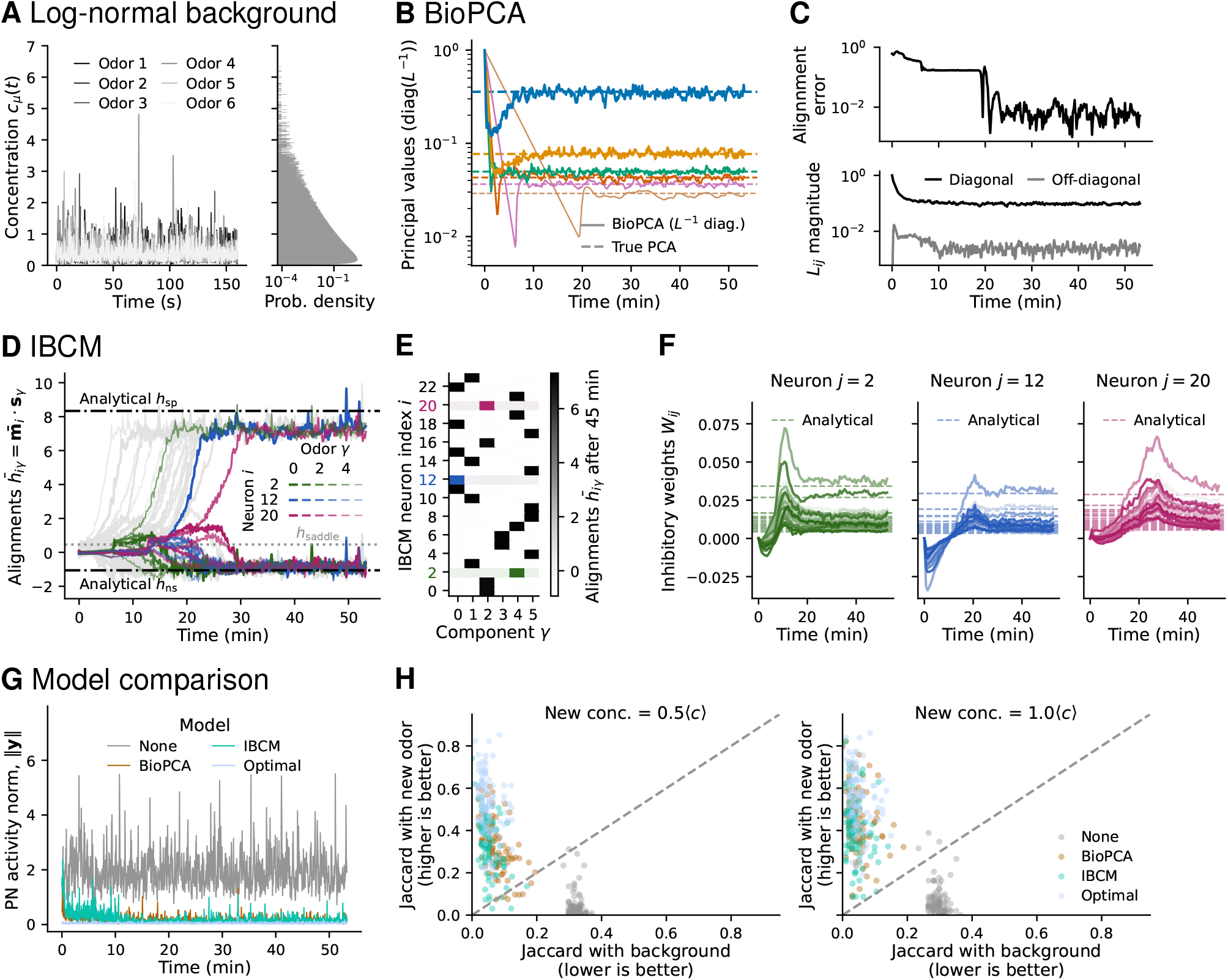
Habituation to log-normal background statistics. (**A**) (Left) Excerpt from the concentration time series of *N*_B_ = 6 odors, and (right) histogram of the concentrations showing they follow a log-normal distribution. (**B**) In a six-neuron BioPCA network, learning dynamics of the *L* matrix diagonal entries, which converge to the background’s principal values, as predicted in section 5 (horizontal dashed lines). (**C**) (Top) Alignment error (defined in *Methods*) of the BioPCA *M* weights with the background subspace, showing that the synaptic weights converge to the principal components vectors, and (bottom) time series of the average diagonal and off-diagonal elements magnitude in *L*, showing the matrix becomes nearly diagonal, as predicted. (**D**) Time series of *N*_I_ = 24 IBCM neurons’ alignment with the background odors **ŝ**_*γ*_, with three neurons highlighted (*N*_B_ = 6 dot products each). Each neuron aligns with one odor and reaches dot product magnitudes close to the analytical expressions for *h*_sp_, *h*_ns_ (section 4E). Different neurons select different odors. (**E**) Table summarizing the alignment of each IBCM neuron (with three highlighted). Plotted values are the dot products 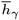 averaged over the last 15 minutes of the simulation example. (**F**) Time series of the *W* weights in the IBCM network, showing their convergence to the analytical predictions (equation 82). (**G**) Example PN time series during habituation by IBCM, BioPCA, and optimal manifold learning networks, compared to the absence of habituation (OSN input). (**H**) Performance of the different models for new odor recognition in log-normal backgrounds, tested in a sample background, across 100 new odors, 10 background samples and 10 test times in that simulation. The plot compares the Jaccard similarity between mixture and new odor responses to the similarity between mixture and background odor responses. Individual dots are medians across background samples and test times. The plot shows that after habituation by the manifold learning models, the response is more similar to the new odor (above the diagonal), while that is not the case in the absence of habituation (below diagonal).

**Fig. S7.**
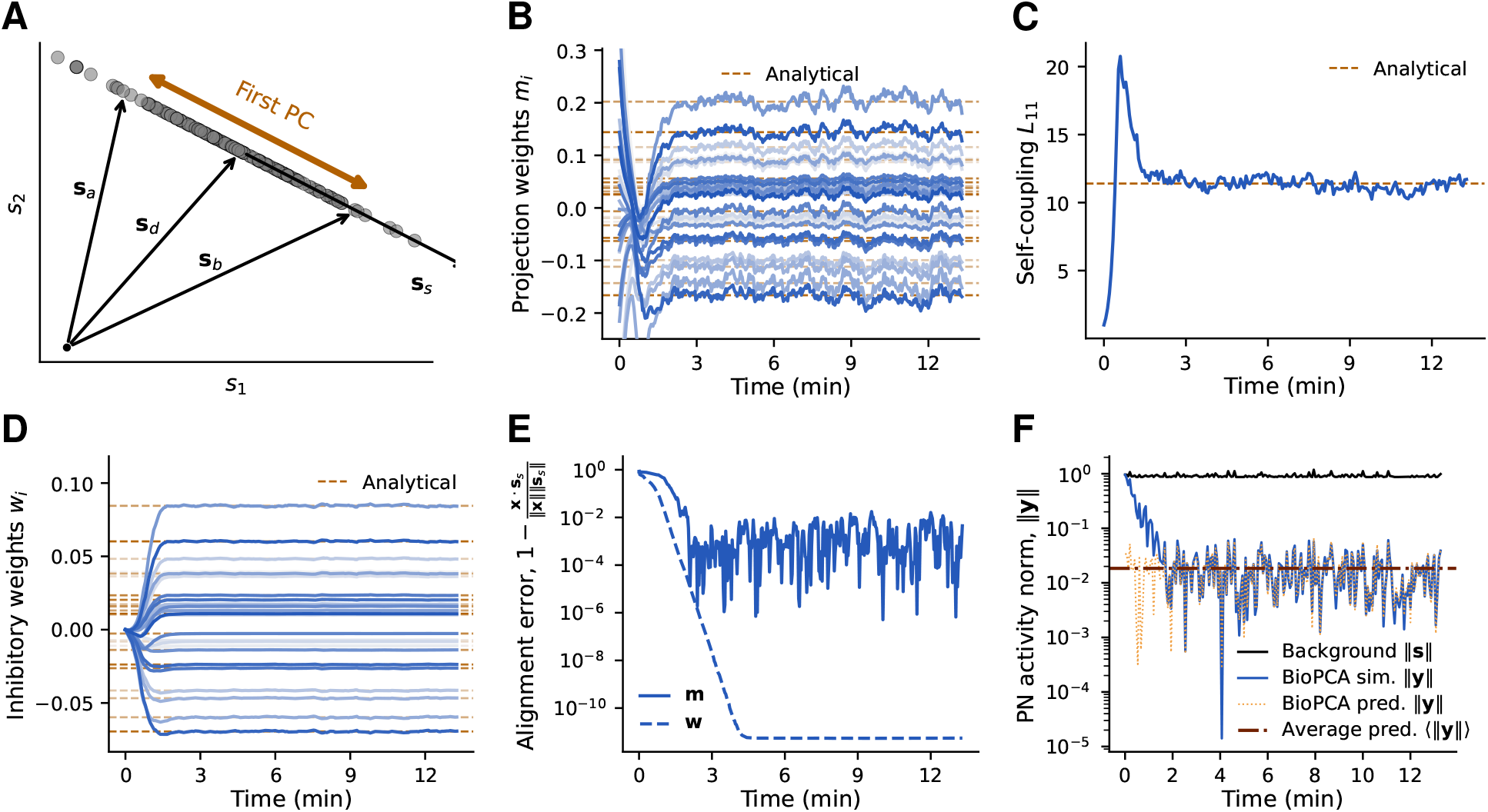
BioPCA model learning dynamics on a simplified background model. We simulate a network with one BioPCA interneuron habituating to the the same two-odor toy model used in Fig. S4. (**A**) The first principal component (PC) corresponds to the stochastic component **s**_s_ along which the background fluctuates. (**B**) Time series of the synaptic weights **m** (single-row matrix *M*) of the BioPCA neuron, which converge to the elements of the first PC vector multiplied by *σ*^2^∥**s**_s_∥ (dashed lines), as predicted in eq. 99. (**C**) Time series of the self-coupling *L*_11_ of this neuron, which converges to the inverse of the eigenvalue (dashed), eq. 98. (**D**) Time series of the inhibitory weights (single-column matrix *W*), which converge to the first PC vector with a scale given in eq. 100. (**E**) Alignment error of the **m** (solid line) and **w** (dashed) vectors with the first PC. Both alignment errors are below 0.1 %; **w** aligns especially well due to its slow Hebbian dynamics (fluctuations are amplified by nonlinearities in the BioPCA equations). (**F**) PN activity norm, ∥**y**(*t*) ∥ (solid blue line), closely matching the analytical predictions for its average (dark orange) and instantaneous (dashed orange) values given the background trajectory, derived in eq. 101. The response to the background is reduced to *∼* 2 % of its original amplitude.

**Fig. S8.**
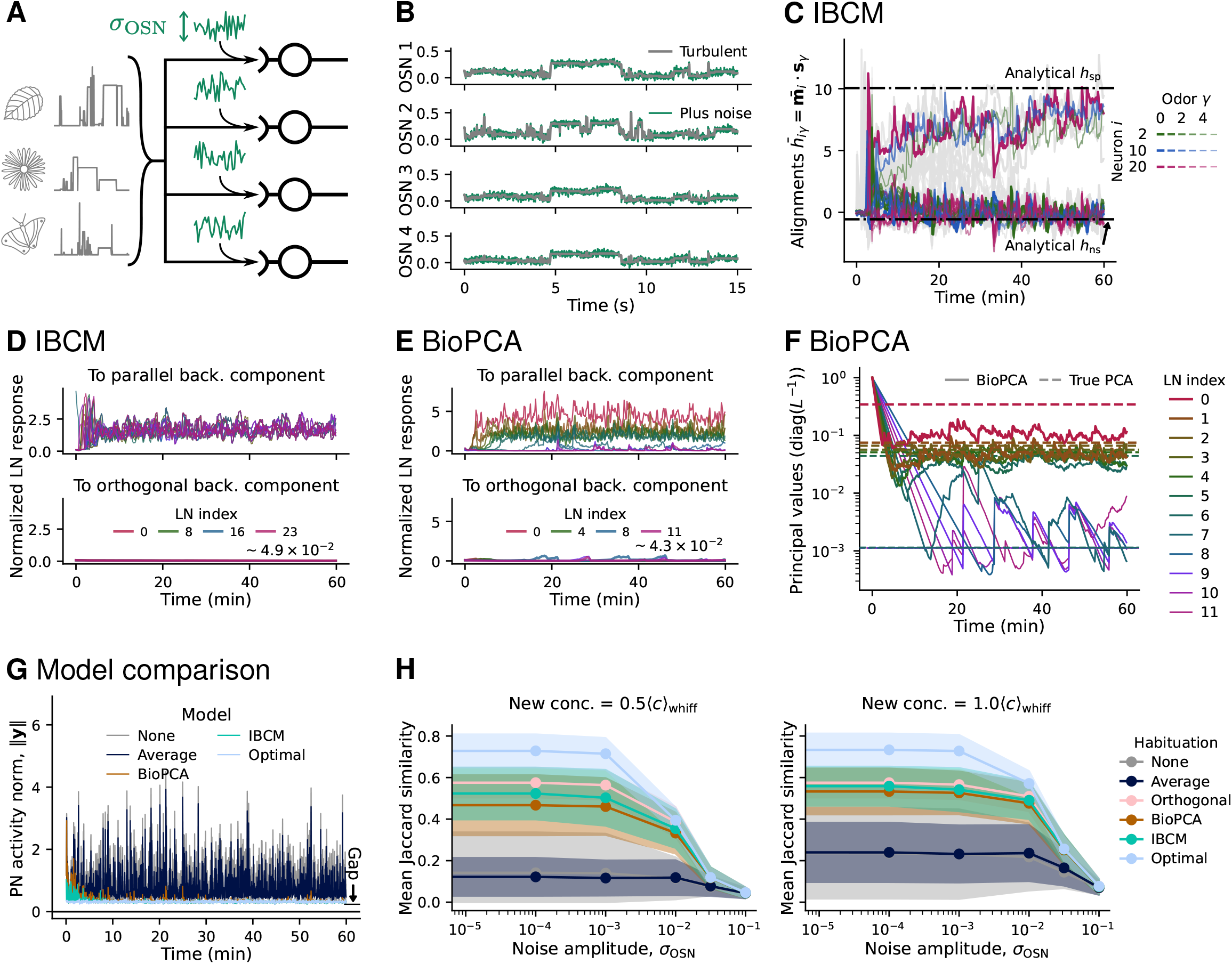
Robustness against OSN noise in the IBCM and BioPCA manifold learning models. (**A**) Illustration of the OSN noise considered. Turbulent backgrounds with six odors are simulated as usual, then independent Gaussian white noise is added to each OSN input at each time point. We consider simulations in *N*_S_ = 100 dimensions (100 OSNs). (**B**) Excerpt from the input time series of four OSNs, showing the turbulent background contribution to the input (grey) and the total input with noise added (green). We illustrate the process with noise amplitude *σ*_OSN_ = 0.01; we will consider different amplitudes in panel (H). (**C**) Time series of the alignment of IBCM neurons with background odors, showing that neurons still become selective for true odors while ignoring the added Gaussian noise components. (**D**) Response of each IBCM neuron, 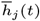, to the parallel (top) and orthogonal (bottom) components of the background, normalized by the average norm of these components, ∥**s**_*//*_∥ or ∥**s**_*⊥*_∥. The time series are smoothed with a moving average filter over a window of 3 s, to visualize the average response amplitude. The parallel background component includes the actual turbulent background odors and the component of the Gaussian noise in that subspace; the orthogonal part contains the noise components orthogonal to background odors. We see that IBCM neurons are only responsive to the true olfactory background, not to noise. (**E**) Similar to (D), but for BioPCA neurons, which also respond only to the true background subspace. (**F**) Learning dynamics of the *L* matrix diagonal elements in the BioPCA network, showing imperfect convergence to the principal values of the background (dashed lines). The first dominant principal component (PC) is missed, but there are *N*_B_ = 7 neurons roughly capturing the background subspace, while remaining neurons capture the small PCs along the pure Gaussian noise directions (eigenvalues equal to *σ*_OSN_). (**G**) Norm of the PN response to the noisy turbulent background over time, showing habituation despite the OSN noise, in all manifold learning models. (**H**) Performance of the various models for new odor recognition as a function of the OSN noise amplitude *σ*_OSN_. Numerical experiments similar to those of Fig. 2, repeated for each *σ*_OSN_ value across 64 backgrounds, 100 new odors, 5 test times and 4 background samples at each time. Lines indicate the average Jaccard similarity, and shaded areas, the standard deviation across repeats. For small noise, the performance remains unchanged, then it rapidly degrades for all models as the OSN noise becomes comparable in magnitude to the new odors. We used *N*_I_ = 24 IBCM neurons, and *N*_I_ = 12 BioPCA neurons.

**Fig. S9.**
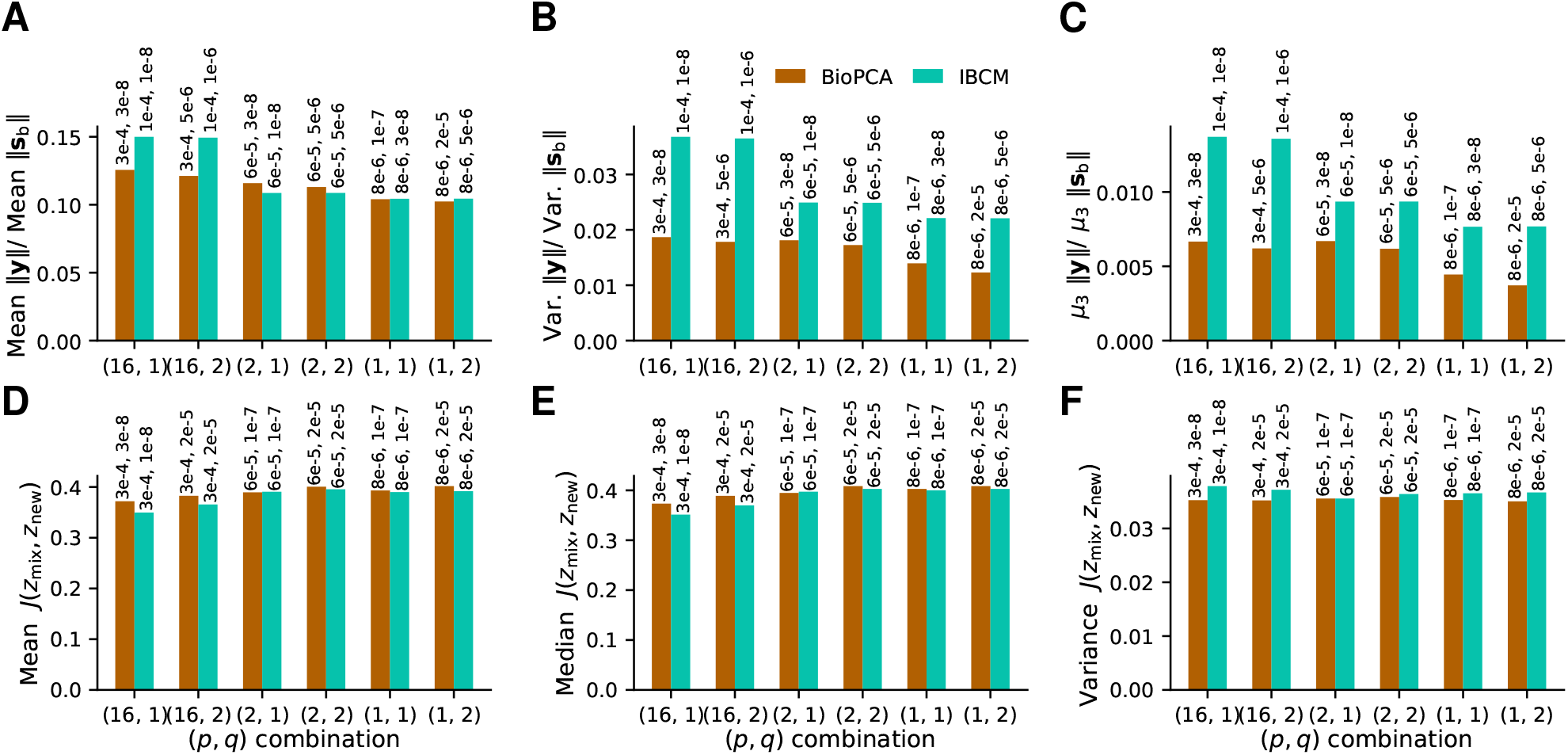
Performance for various *L*^*p*^ norms in *W* ‘s cost function learning rule. (**A**) Reduction of the mean, (**B**) variance, (**C**) and third central moment of the norm of the PN response after habituation by IBCM of BioPCA networks. We consider various *p, q* norm choices in the generalized Hebbian learning rules derived in section 7. For each *p, q* combination, we performed a grid search over learning rates *α* and *β*; the bar graph reports the best performance for each *p, q*; the *α, β* rates producing this performance are annotated above the bars. (**D**) Mean and (**E**) median Jaccard similarity between the response to mixtures and the response to the new odor alone, after habituation. We test for various *p, q* combinations as in (A)-(C). (**F**) Variance in the Jaccard similarity; lower variance is better when the mean or median similarity is high – it signifies that the network is more consistent across backgrounds and trials.

**Fig. S10.**
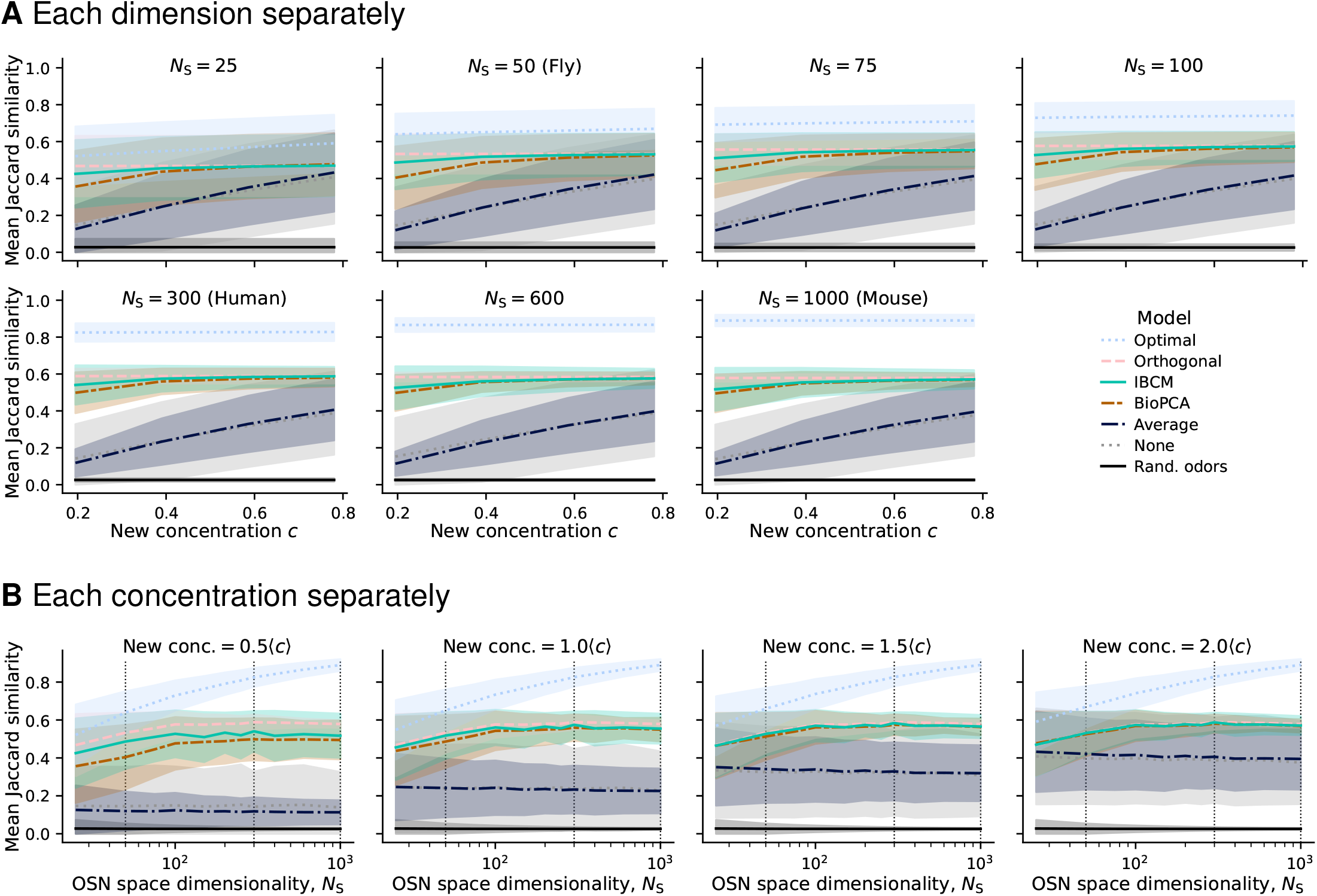
Odor recognition performance as a function of dimensionality and new odor concentration. Supplement to main Figure 5. (**A**) Jaccard similarity as a function of new odor concentration, for each tested dimensionality separately. (**B**) Jaccard similarity as a function of dimensionality, for each tested new odor concentration. Lines indicate the mean Jaccard similarity, and shaded areas, the standard deviation across repeats. “Rand. odors” indicates the similarity between two odors drawn at random, to show that the similarity of the response to the new odor is significantly higher than the similarity that would occur by chance.

**Fig. S11.**
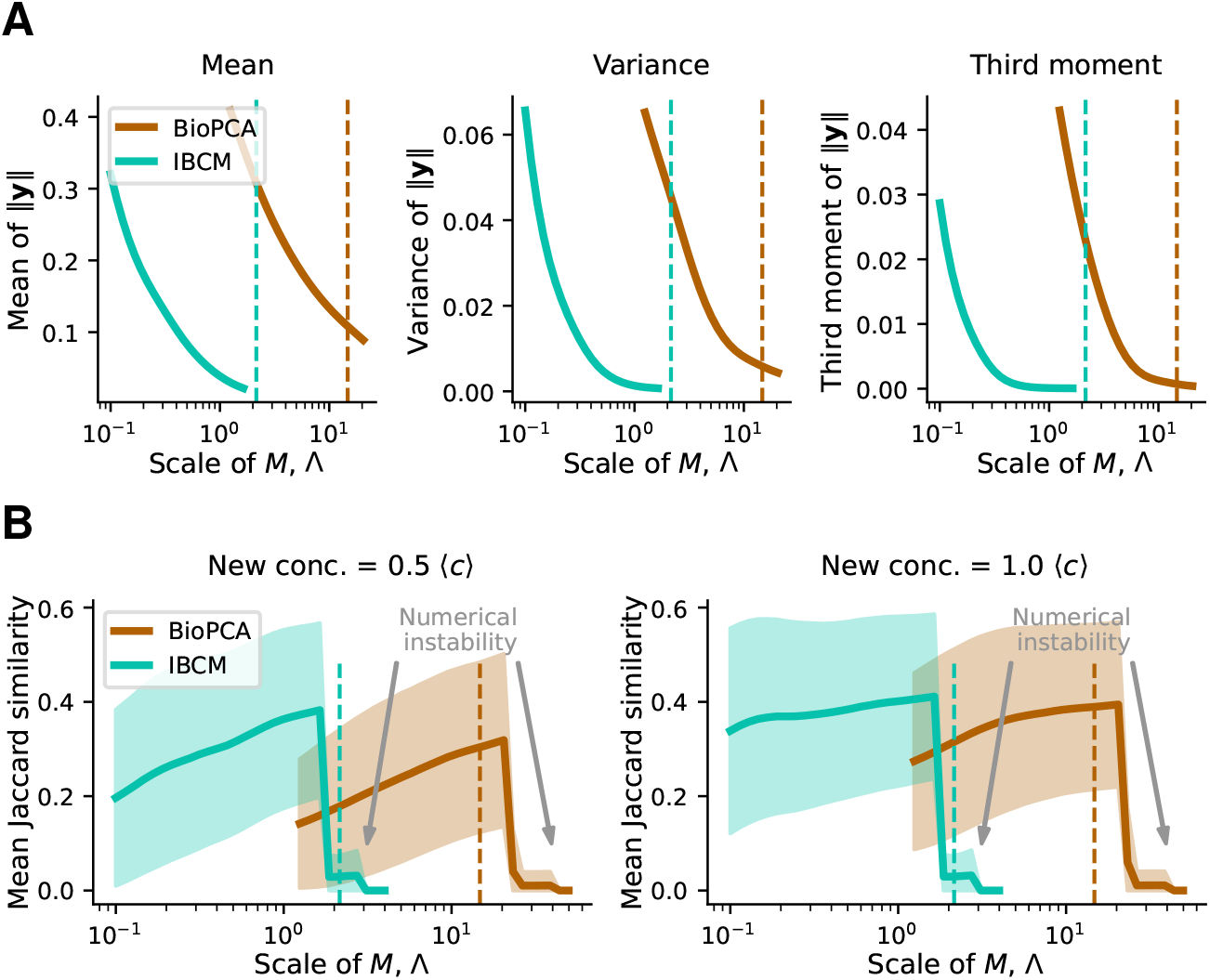
Effect of the *M* weights scaling parameter Λ, to match the IBCM and BioPCA models. (**A**) Habituation performance, quantified by reduction in the mean, variance, and third moment of the PN response to the background, **y**. Habituation runs are performed on the same background time series, with different Λ values for each model. (**B**) New odor recognition performance, quantified by the increase in mean Jaccard similarity between the response to mixtures and the new odor tag, for two new odor concentrations. Same background example as (A), testing 100 new odors at the end (shaded area: standard deviation across tested odors). BioPCA and IBCM can be made to perform equally well by setting Λ appropriately (Λ *∼* 10 times larger for BioPCA). Vertical dashed lines indicate the scale at which numerical instabilities are expected to arise according to a nonlinear stability analysis of the Euler integrator (section 2F). Simulations were run in *N*_S_ = 25 dimensions and with default parameter values.

**Table S1.**
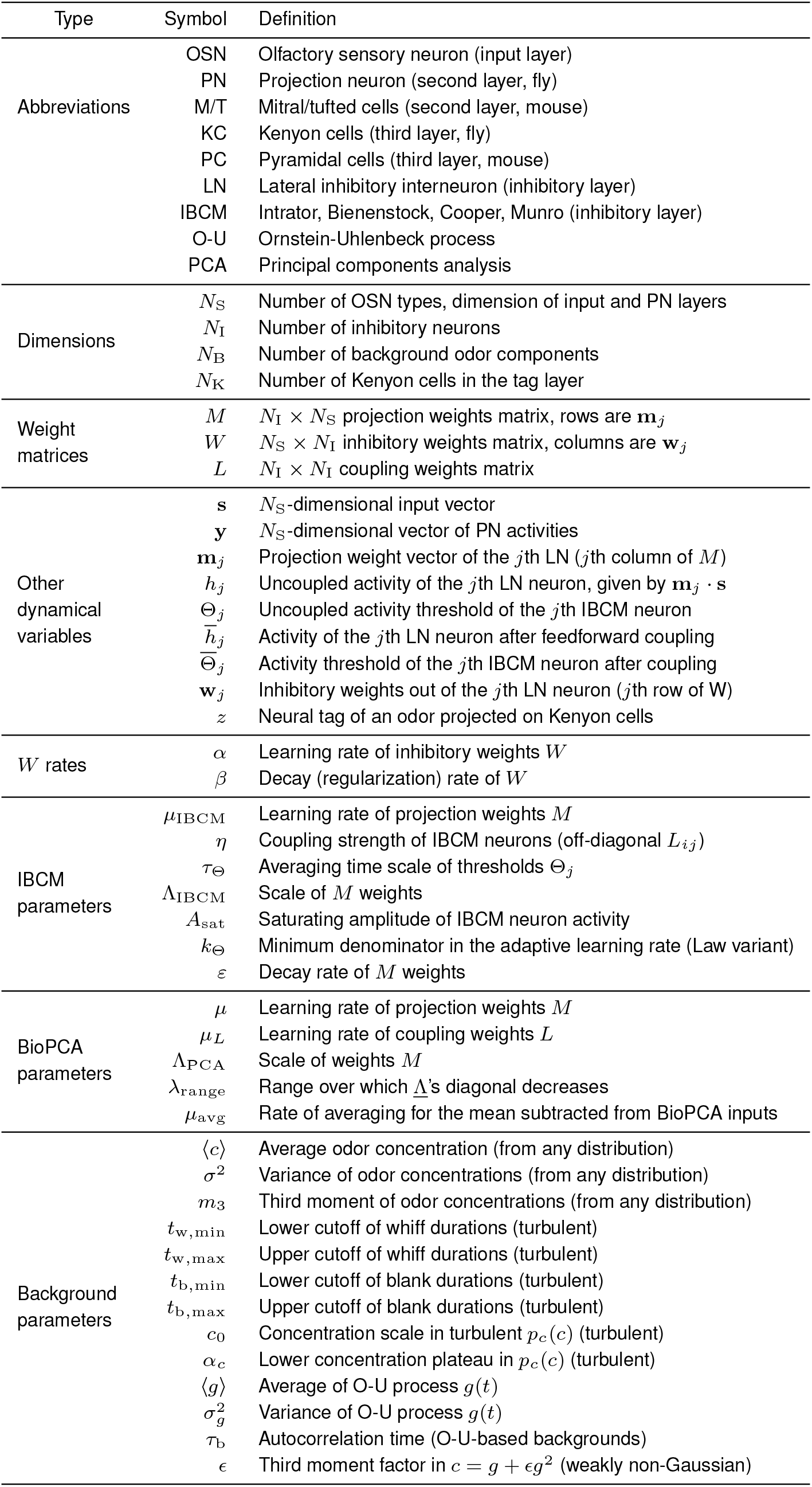
Definition of olfactory network model parameters.

**Table S2.**
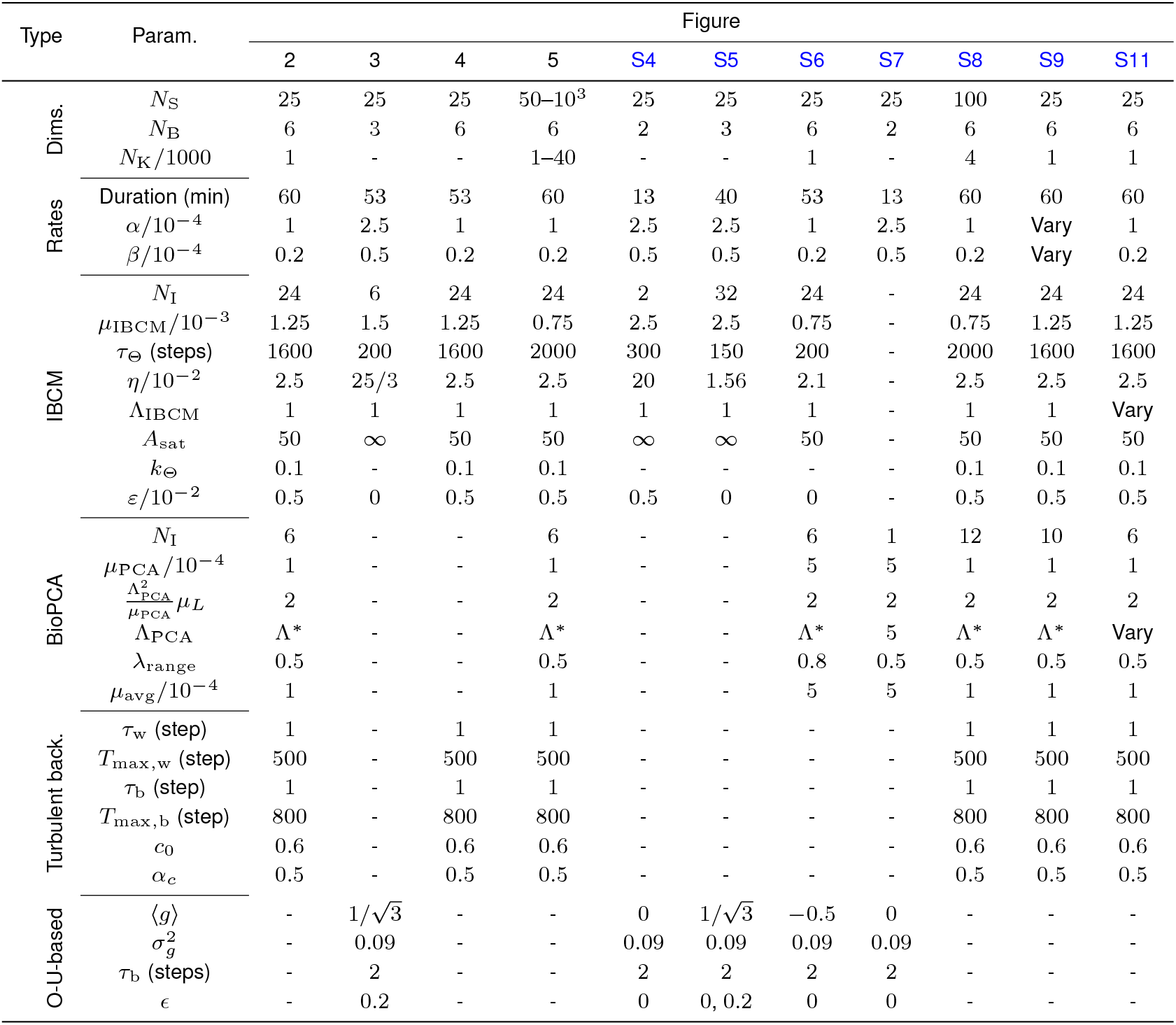
Value of model and simulation parameters. Fig. S2 uses the same parameters as 2. Fig. S3 uses the same parameters as the simulations related to each panel (A: Fig. 3, B: Fig. S6, C: Fig. 2). Fig. S10 uses the same parameters as 5. Time steps in simulations correspond to 10 ms (such that 360, 000 steps make 60 minutes). Λ^*∗*^ is the Λ_PCA_ value predicted to make it equivalent to IBCM, given in eq. 106.

## Footnotes

1 We have managed to reduce the degree from cubic to quadratic because the substitutions eliminated one root *y*_1_ = *y*_2_ =0, which was not interesting.

